# A structural model of a Ras-Raf signalosome

**DOI:** 10.1101/2020.07.15.165266

**Authors:** Venkatesh P. Mysore, Zhi-Wei Zhou, Chiara Ambrogio, Lianbo Li, Jonas N. Kapp, Chunya Lu, Qi Wang, Maxwell R. Tucker, Jeffrey J. Okoro, Gabriela Nagy-Davidescu, Xiaochen Bai, Andreas Plückthun, Pasi A. Jänne, Kenneth D. Westover, Yibing Shan, David E. Shaw

**Author notes:** These authors contributed equally to the manuscript (Venkatesh P. Mysore, Zhi-Wei Zhou, Chiara Ambrogio). To whom correspondence should be addressed. Yibing Shan, David E. Shaw.

## Abstract

The protein K-Ras functions as a molecular switch in signaling pathways regulating cell growth. In the MAPK pathway, which is implicated in many cancers, multiple K-Ras proteins are thought to assemble at the cell membrane with Ras-effector proteins from the Raf family. Here we propose an atomistic structural model for such an assembly. Our starting point was an asymmetric, GTP-mediated K-Ras dimer model, which we generated using unbiased molecular dynamics simulations and verified with mutagenesis experiments. Adding further K-Ras monomers in a head-to-tail fashion led to a compact helical assembly, a model we validated using electron microscopy and cell-based experiments. This assembly stabilizes K-Ras in its active state and presents composite interfaces to facilitate Raf binding. Guided by existing experimental data, we then positioned C-Raf, the downstream kinase MEK1, and accessory proteins (Galectin-3 and 14-3-3σ) on the helical assembly. The resulting Ras-Raf signalosome model offers an explanation for a large body of data on MAPK signaling.

## Introduction

The protein K-Ras is a crucial molecular switch in signaling pathways that regulate cell proliferation, differentiation, and survival.^1^ Gain-of-function mutations in three closely related members of the Ras family, K-Ras, H-Ras, and N-Ras, are found in approximately 25% of all human cancers, with mutations in K-Ras being particularly common.^2^ In many cases, oncogenic mutations in Ras are associated with hyperactivation of the mitogen-activated protein kinase (MAPK) signaling pathway.^3^ Dysregulation of the MAPK pathway has also been implicated in a host of hyperproliferative developmental disorders termed RASopathies.^4^ Inhibiting oncogenic or aberrant Ras is thus of great clinical interest.^5,6^ Our structural understanding of how K-Ras activates downstream MAPK signaling, however, has been incomplete.

Ras is active when GTP-bound and inactive when GDP-bound; conversions between these states are catalyzed by guanine nucleotide exchange factors (GEFs) and GTPase-activating proteins (GAPs), respectively.^7^ Active and inactive Ras conformations differ primarily at the so-called “switch” regions involved in effector binding.^8,9^ Despite some early evidence that Ras can form oligomers,^10^ activated Ras has long been thought to act in MAPK signaling as a monomeric membrane anchor for C-Raf or other effectors from the Raf kinase family (Ras itself being anchored to the plasma membrane by its lipidated C-terminal tail). A growing body of data now indicates that Ras proteins dimerize,^11–14^ that their dimerization may depend on GTP binding,^15^ and that Ras dimerization may be critical for Raf activation.^16–19^

A significant fraction of membrane-bound, GTP-bound Ras proteins have been found to organize into relatively immobile so-called “nanoclusters,” each with about eight members and a radius of ~100 Å.^20,21^ Ras dimerization appears to be crucial to nanocluster formation,^18^ and it is possible that nanoclusters comprise networks of loosely interacting, lower-order Ras structures (such as monomers and dimers); structurally well-defined, higher-order oligomers; or a mixture of both. It is known that Galectin-3 (Gal-3) is crucial for K-Ras nanoclustering and signaling,^22^ as is Galectin-1 (Gal-1) for H-Ras.^23^ Raf kinases co-localize with nanoclustered Ras proteins; it has been shown that such co-localized effectors generate the majority of the downstream MAPK signal,^21,24^ and that Ras activity can be suppressed by proteins that disrupt this clustering.^18,25^ Structural information about nanoclusters is sparse, however, because they are difficult to reconstitute in vitro and the resolution of cellular imaging is limited.

In the work reported here, we used unbiased molecular dynamics (MD) simulations to inform the construction of an atomistic structural model of a K-Ras nanocluster. We term our proposed structure a *Ras-Raf signalosome* because it is a higher-order hetero-oligomer (containing multiple monomers of K-Ras, C-Raf, Gal-3, and other proteins) with well-defined structural features and is reminiscent of “signalosomes” that underlie the activities of other signaling systems.^26^ We first generated a GTP-mediated asymmetric dimer model by directly simulating the association of two K-Ras monomers; subsequent mutagenesis experiments strongly supported this model. The asymmetric dimer model was then extended to a higher-order oligomer model by adding monomers in a head-to-tail fashion, resulting in a compact helical K-Ras assembly. This assembly promotes the stability and accessibility of active K-Ras and creates composite interfaces that facilitate Raf binding. Guided by existing experimental data and further simulations, we then positioned C-Raf dimers, the downstream kinase MEK1, and the accessory proteins Gal-3 and 14-3-3σ around the K-Ras core. Our Ras-Raf signalosome model, which synthesizes a large body of data on the MAPK pathway, may extend to other Ras and Raf isoforms and provides a structural framework to inform further investigations of MAPK signaling.

## Results

We first summarize key structural features of our Ras-Raf signalosome model for convenience before detailing its construction and resulting findings.

### An overview of the Ras-Raf signalosome model

In brief, we obtained our Ras-Raf signalosome model by proposing a GTP-mediated asymmetric K-Ras dimer model (Figure 1A), extending it to a K-Ras helical assembly (Figure 1B), and adding further components. The final model is anchored to the plasma membrane, and the Ras-binding domains (RBDs) and cysteine-rich domains (CRDs) of C-Raf, along with Gal-3, interact directly with the K-Ras helical assembly (Figure 1C), essentially covering the outside of the assembly. Unstructured linkers connect the CRDs of C-Raf proteins to their kinase domains, which are dimerized and located at the periphery of the structure (Figures 1D and 1E), in complex with MEK1 kinases and 14-3-3 dimers (Figure 1F). Although the signalosome is structurally open-ended, making a range of signalosome sizes possible, we anticipate this size range will be constrained by GAP and GEF regulation at the cell membrane. In this report we mostly focus on an 8-protomer signalosome as an example system. The signalosome (Movie S1 and Figures 1D and S1A; atomic coordinates in Dataset S1) produces a millimolar local concentration of C-Raf (see SI), which would be sufficient to ensure the dimerization-dependent activation of C-Raf.^27^ The model incorporates seven previously resolved single- or double-domain structures of the constituent proteins (Figure 1G). The following sections describe in greater detail our construction of this signalosome model, which was informed by existing experimental data, new experiments conducted as part of the current study, and previous experience with MD simulations of protein-protein and protein–small molecule association.^28–30^

**Figure 1.**
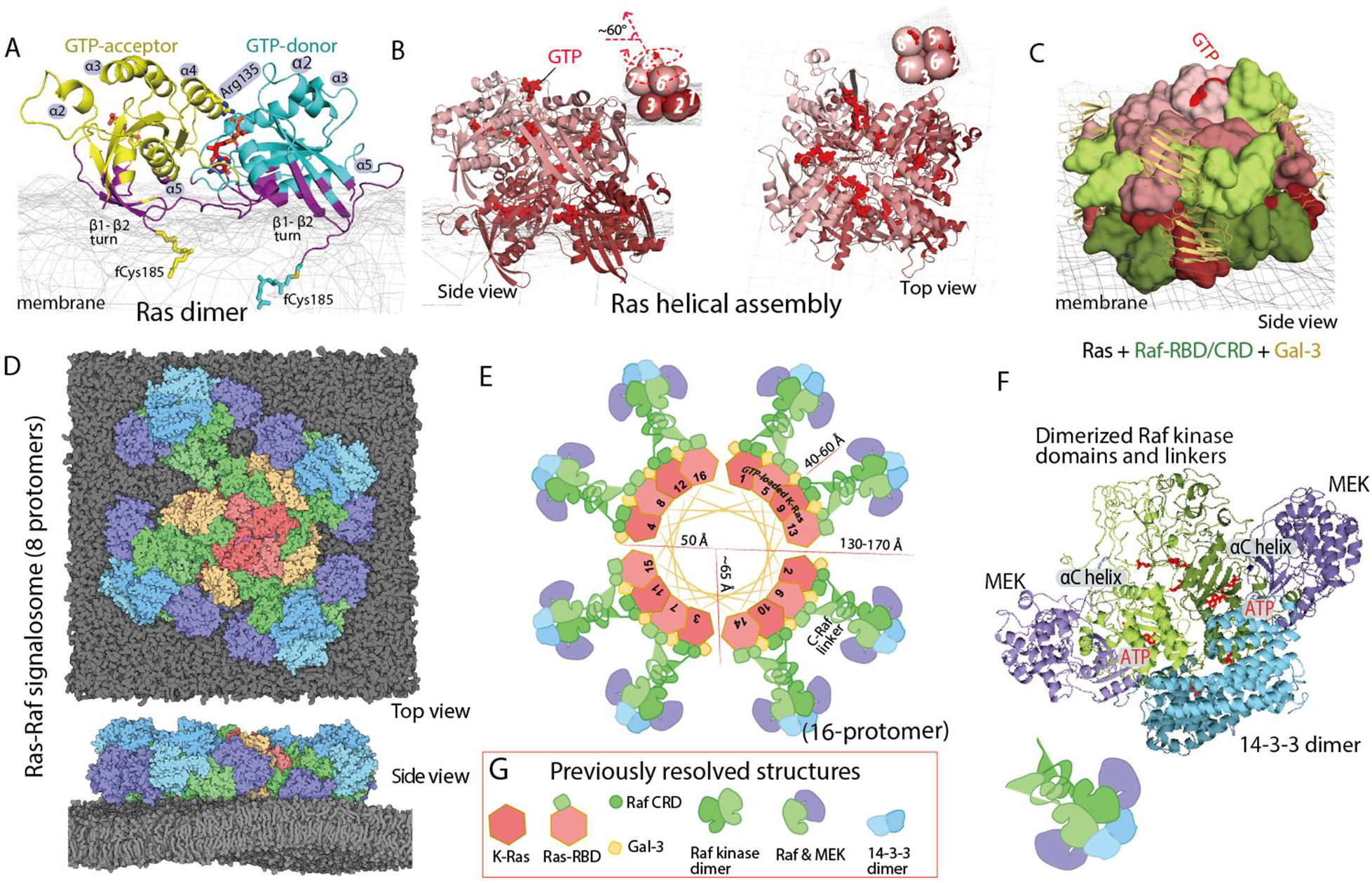
Outline of the structural model of a Ras-Raf signalosome. **A**. The GTP-mediated asymmetric K-Ras dimer model. Residues contacting the membrane (mesh) are shown in purple; the GTP at the interface is in red. **B**. The (8-protomer) K-Ras helical assembly with illustrative cartoons. **C**. The K-Ras helical assembly (pink), decorated with C-Raf RBD and CRD domains (green) and Gal-3 proteins (yellow). **D**. The signalosome model on the membrane. The 8-protomer model contains four catalytic units at the periphery. **E**. Diagram of the signalosome model with a helical wheel representing the Ras assembly at the center. To illustrate how the signalosome could grow beyond 8 protomers, 16 protomers are shown. **F**. One catalytic unit centered on a Raf kinase dimer. ATP and phosphorylated Raf residues (pSer338, pTyr341, and pSer621) are in red. **G**. Previously resolved structures used as building blocks in the modeling: the K-Ras monomer^79^ (PDB 4DSN), H-Ras/Raf RBD complex^45^ (PDB 4G0N), C-Raf CRD^80^ (PDB 1FAR), Gal-3^81^ (PDB 3ZSM), C-Raf kinase domain dimer^64^ (PDB 3OMV), B-Raf/MEK1 complex^65^ (PDB 4MNE), and 14-3-3σ dimer bound with C-Raf phosphopeptide^63^ (PDB 4IEA).

### A GTP-mediated asymmetric K-Ras dimer favors the active state

We first performed 20 simulations (680 μs total simulation time) of two GTP-bound K-Ras proteins (PDB 4DSN) in aqueous solvent (Figure S2A, left). In one simulation, the two K-Ras proteins formed stable interactions mediated in part by a bound GTP (Movie S2). This model is compelling because it provides a direct explanation for the GTP-dependence of K-Ras dimerization.^15^ Hereafter we will refer to this model as the GTP-mediated asymmetric (GMA) dimer model.

Because K-Ras dimerization occurs at the membrane, we then performed 23 simulations (363 μs total simulation time) of two GTP-bound K-Ras proteins anchored to the membrane by their farnesylated Cys185 (fCys185) residues^31^ (Figure S2A, right). In one of these membrane simulations (Figure 2A and Movie S3), the K-Ras proteins also formed the GMA dimer; the structure is virtually identical to that obtained from the solvent simulations (Figure 2B, upper panel). The GMA dimer (atomic coordinates in Dataset S2) remained stable for nearly 100 μs in both the solvent and membrane simulations (Figure 2B). We note that, because the timescale of the individual simulations (tens of microseconds) of K-Ras dimerization is smaller than the typical timescale of protein-protein association and dissociation, it is to be expected that most such simulations will not reach the native dimer structure (Figure S2A), and that taken together these simulations will not yield the correct Boltzmann weight of the K-Ras dimer (see SI).

**Figure 2.**
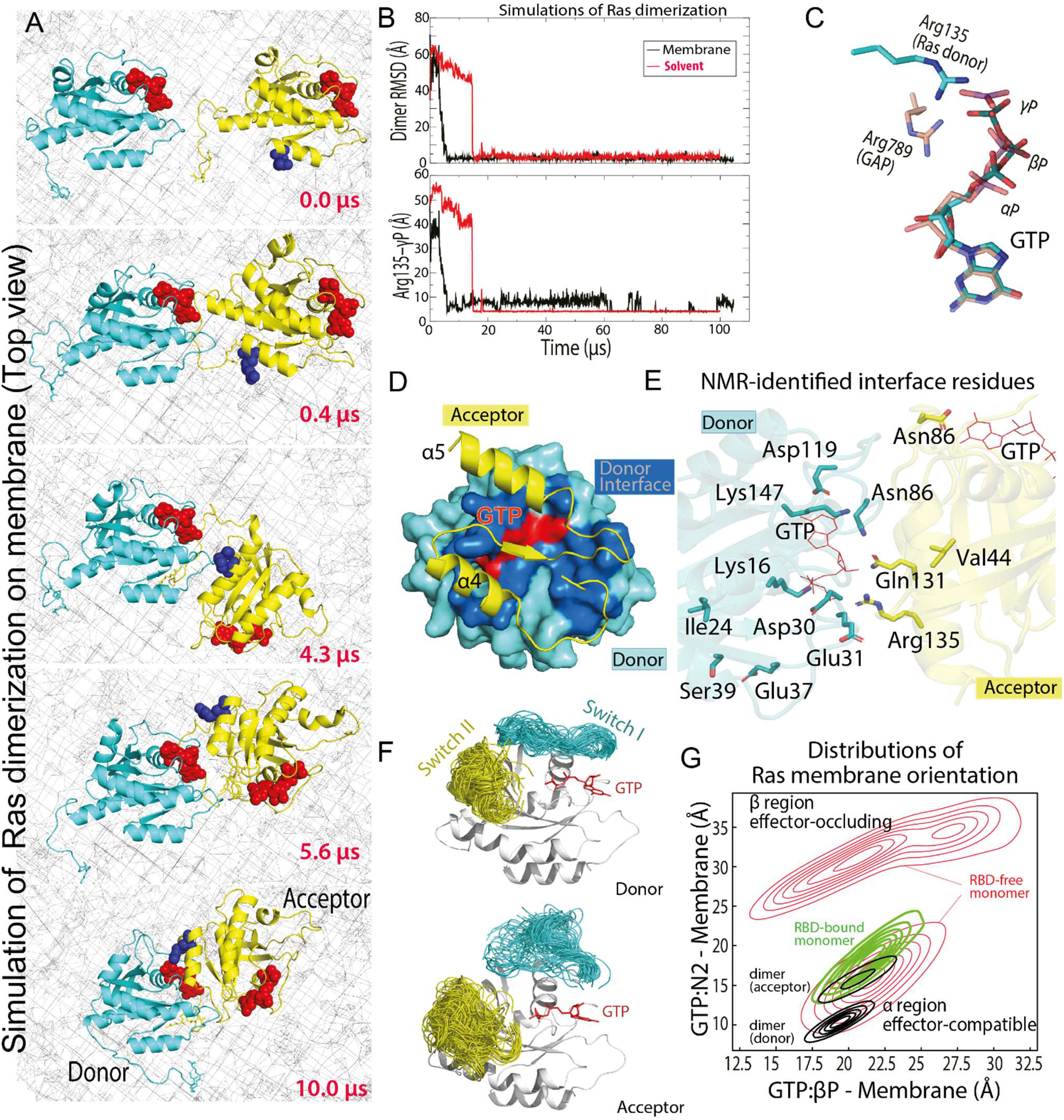
The GTP-mediated asymmetric dimer model of K-Ras. **A**. Snapshots, taken at the simulation times shown, of the GTP-mediated dimerization of membrane-anchored K-Ras proteins (top view; membrane shown as a mesh). **B**. Upper panel: RMSD of the two K-Ras proteins in the solvent and in the membranous simulation of K-Ras dimerization with respect to the final GTP-mediated asymmetric K-Ras dimer model (the snapshot at 100 μs from the simulation shown in (A)). Lower panel: the Arg135 distance to the GTP γ-phosphate at the dimer interface in the simulations. **C**. Comparison of the GTP-Arg135 (or Lys128) interaction with the GAP “arginine finger.” **D.** The dimer interface; only part of the acceptor is shown. **E**. All 12 NMR-identified residues^15^ involved in K-Ras dimerization, shown with respect to the GMA dimer interface. **F**. The conformations of the switch I and II regions in simulations of the GMA dimer. **G**. Distributions of K-Ras membrane orientations in simulations of a K-Ras monomer, an RBD-bound K-Ras monomer, and the GMA dimer in the membrane environment; the orientations are mapped to a 2D space defined by the distances of the GTP β-phosphorus (βP) and the GTP amine nitrogen of the guanine ring (N2) to the membrane plane (see Figures S2D and S2E). The α and β regions correspond to the α and β orientations,^39^ respectively; in the former, the α4 and α5 helices contact the membrane, and in the latter, the β strands do.

At the GMA dimer interface, a key interaction occurs between the GTP γ-phosphate of one K-Ras protein and Arg135 or Lys128 of the other (Figures 2B, 2C, and S2B). Hereafter we will refer to the former K-Ras protein as the GTP-donor (or more briefly, the donor) and the latter as the GTP-acceptor (or simply the acceptor). Unlike the so-called “arginine finger” interaction between Ras and GAP, whereby an arginine mediates electron transfer from the γ-phosphate to the β-phosphate^32^ and catalyzes GTP hydrolysis,^33^ the GMA dimer model has Arg135 or Lys128 interacting only with the γ-phosphate (Figure 2C), hindering electron transfer and thus GTP hydrolysis.

The GMA dimer interface has a buried surface area of ~1750 Å^2^, and pronounced electrostatic complementarity (Figure S2H). The donor interface involves parts of the switch I region and the β4-α3, β5-α4, and β6-α5 loops. The acceptor interface primarily involves the α4 and α5 helices and the β6 strand (Figure 2D), and largely overlaps with the interface for NS1 (Figure S2G), a synthetic protein that disrupts Ras dimerization.^18^ The GMA dimer is consistent with existing nuclear magnetic resonance (NMR) spectroscopy data: Residues that exhibited broadening of ^15^N-HSQC spectra upon sample dilution^15^ are predominantly located at the dimer interface (Figure 2E; see SI for further discussion). The interface is mostly polar (Figure S2C), with several salt bridges (e.g., Asp154-Lys147, Arg161-Asp30/Glu31, and Glu62-Lys128/Arg135) at its periphery (Figures 3A and 3B). In our simulations, Arg135 interacted with the GTP if Lys128 interacted with Glu62, and vice versa (Figure S2B).

**Figure 3.**
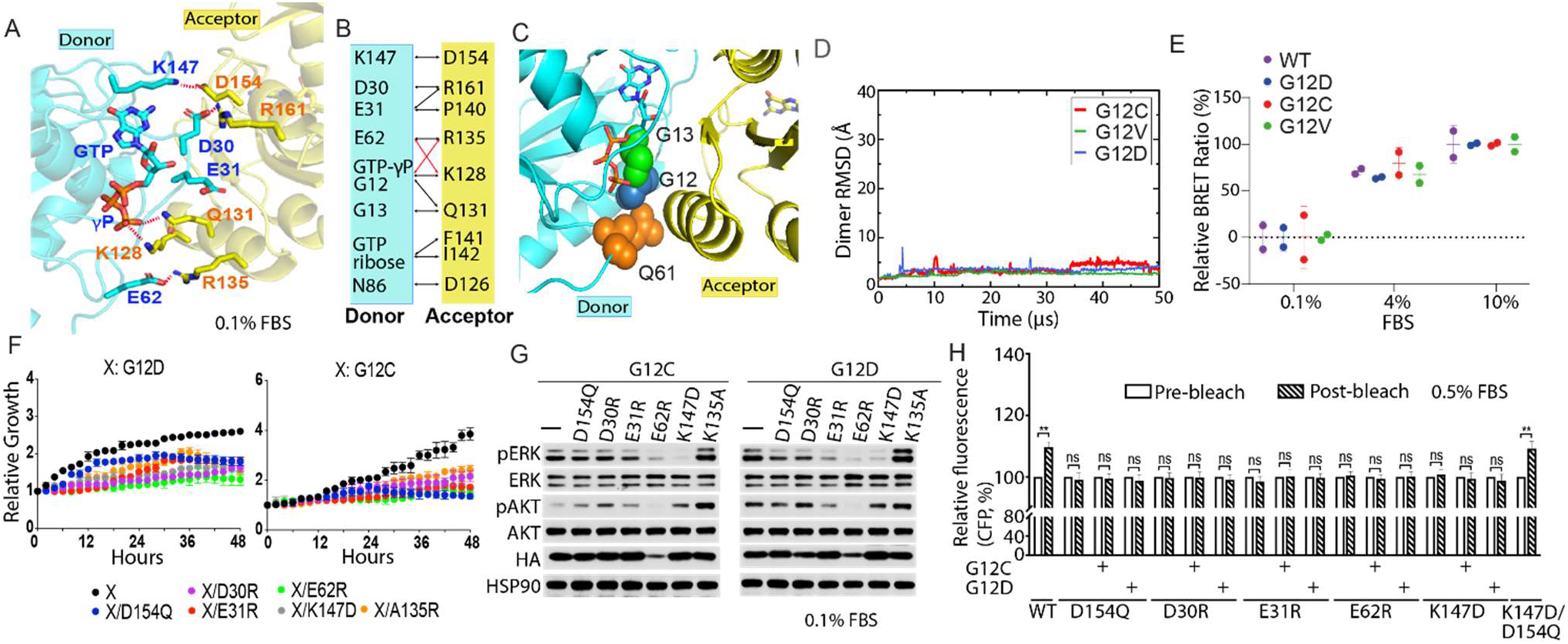
Mutagenesis validation of the GMA dimer interface. **A**. The residues involved in the salt bridges at the dimer interface. **B**. Residue contacts at the dimer interface (black and red lines). Red lines highlight E62-K128 and GTP-γP-R135 contacts, which alternate with E62-R135 and γP-K128 contacts. **C**. G12, G13, and Q61 oncogenic mutation sites at the GMA dimer interface. **D**. The stability of the G12C, G12D and G12V GMA dimers in simulations in terms of Cα RMSD. **E**. BRET signals, which provide an indication of K-Ras assembly, for WT K-Ras and three oncogenic mutants at three different FBS concentrations. . **F**. Growth rates of *K-Ras*^*lox*^/*K-RAS*^*MUT*^ cells shown as confluence values measured by IncuCyte. Results are averages of three duplicates with error bars (which are invisible if smaller than the symbols). G12C or G12D are in the background of the tested mutations. Representative images at the end point are shown in Figure S4A. Cells were kept in 0.1% FBS (see Figure S4B for curves obtained in 0.5%, 1%, and 10% FBS). **G.** Phosphorylation of ERK and AKT in *K-Ras*^*lox*^/*K-RAS*^*MUT*^ cells. Cells were lysated after 48 hours incubation. See Figure S4C for the phosphorylation at other FBS levels. **H**. CFP emission in HEK293T cells co-transfected with CFP- and YFP-fused K-Ras (wild type and various mutants), where binding between the CFP- and the YFP-fused protein would bring the fluorescent tags into close proximity, leading to an increased CFP emission after YFP bleaching. Error bars represent mean ± standard deviation (** denotes p < 0.05 by unpaired Student’s t-test; ns stands for not significant). G12C or G12D is in the background of the tested mutations.

### The GMA dimer stabilizes the active state of K-Ras

The GMA dimer may promote RBD binding by stabilizing the active Ras conformation and favoring membrane orientations that accommodate the RBD. It has been shown that GTP binding favors the active conformation^34^ but does not fully stabilize it.^9,35–38^ In our simulations, although the switch I region visited the active conformation with greater frequency in a GTP-bound K-Ras monomer than in a GDP-bound one (Figure S3A), the inactive conformation was prevalent in both (Figures S3B and S3C). In contrast, in simulations of the GMA dimer, the switch I region of the donor (but not that of the acceptor) was largely stabilized in active-like conformations (Figures 2F and S3A).

Our GMA dimer model also constrains the membrane orientation of K-Ras in a way that is consistent with prior NMR data^39^ and compatible with RBD binding. In the GMA dimer formed on the membrane (Figure 1A), the donor is in the so-called “α orientation,”^39^ and the acceptor is in a similar orientation (Figure 2G). In H-Ras, the β2-β3 strands and α5 helix form a switch region regulating membrane interactions;^40^ for both proteins in the GMA dimer, this switch region contacts the membrane. Both the donor and acceptor orientations are compatible with RBD binding (Figure S2E). This contrasts with monomeric K-Ras, which mostly adopted the β membrane orientation that occludes RBD binding (Figures 2G and S2D).

### Most oncogenic mutations are compatible with the GMA dimer

The most common cancer mutations occur at residues 12, 13, and 61; the former two residues are at the donor interface of the GMA dimer, and the latter one is immediately adjacent to it (Figure 3C). We simulated G12V, G12D, G12C, G13C, G13S, Q61L, and Q61R GMA dimers, and found that the dimers remained stable in 50-μs simulations (Figures 3D and S2I), likely because the largely polar interactions at the GMA dimer interface (which is partially hydrated) can sterically accommodate the mutations. (G13R and G13D behave differently in our simulations, disrupting the GMA dimer (Figure S2I), possibly because they are bulky and more centrally located in the interface (see SI).) To experimentally test the ability of K-Ras assemblies (which result in close proximity of the N termini of K-Ras proteins) to form in the presence of such mutations, we performed bioluminescence resonance energy transfer (BRET) using K-Ras fused with reporter molecules. The BRET experiment showed that indeed, to similar degrees, the wild type and the G12C, G12V, and G12D mutants in cells form assemblies under various fetal bovine serum (FBS) concentrations (Figure 3E).

### Experimental validation of the GMA dimer model

A so-called “*K-Ras*^*lox*^/*K-RAS*^*MUT*^” inducible system—wherein cell lines generated from Ras-less mouse embryonic fibroblasts (MEFs)^41^ are dependent on exogenous K-Ras expression for proliferation—has been reported in a previous study,^19^ along with a cell-based fluorescence resonance energy transfer (FRET) platform, to probe K-Ras/K-Ras interactions. In both experiments, K-Ras signaling is constitutive with introduction of oncogenic K-Ras mutations such as G12D or G12C. Because G12D and G12C mutations are compatible with the dimer (Figures 3D and 3E), we decided to use these two systems to test the GMA dimer. In the previous study, *K-Ras*^*lox*^/*K-RAS*^*G12C*^ or *K-Ras*^*lox*^/*K-RAS*^*G12D*^ lines were used to show that D154Q and R161E mutations in the background of either G12C or G12D impair cell fitness by disrupting K-Ras/K-Ras interactions. Although the dimer model proposed in that study differs from the GMA dimer, the GMA dimer model is also consistent with those findings, because Asp154 and Arg161 are involved in key salt bridges at the GMA dimer interface (Figure 3A).

To further validate the GMA dimer model, we have used the MEF and FRET systems to test residues (Asp30, Glu31, Glu62, and Lys147) involved in salt bridges at the GMA dimer interface (Figures 3A and 3B). We found that the mutations D30R, E31R, E62R, and K147D each impaired cell growth (Figures 3F and S4B) and reduced phosphorylation of ERK kinase, which is downstream of K-Ras in the MAPK pathway (Figures 3G and S4C). The mutations also disrupted the FRET signal of K-Ras/K-Ras interactions (Figures 3H and S5A). We further found that a D154Q/K147D double mutation, which was predicted to restore the interaction, indeed recovered the FRET signal (Figures 3H and S5A). FRET analysis also showed that mutations known to disrupt the active conformation of Ras, such as T35A, T35S,^37^ and G60A,^42^ also disrupted K-Ras/K-Ras interactions (Figures S5B and S5C), supporting the connection between GMA dimerization and the active conformation. We also mutated Arg135 at the GMA dimer interface, but the effects of R135A on dimerization were masked by the raised K-Ras nucleotide exchange rate caused by this mutation (Figures S5D and S5E).

### Extrapolation of the GMA K-Ras dimer into a K-Ras helical assembly

Adding K-Ras proteins to the GMA dimer in a head-to-tail fashion extends it into a left-handed helical assembly, which is highly compact and yet does not exhibit any steric clashes (Figure 1B). In this assembly, each K-Ras protein serves as both a donor and an acceptor (with the exception of the “head” serving only as a donor and the “tail” only as an acceptor). For convenience, we number the K-Ras proteins in the assembly from 1 to *n* starting from the head (and incrementing by one from a donor to its acceptor). The helical assembly consists of four K-Ras proteins per turn and has a radius of ~50 Å and a pitch of ~40 Å, with virtually no unoccupied space between turns or in the center. We placed the K-Ras assembly on the membrane, with the axis of the helical assembly at a ~60° angle to the membrane surface (Figure 1B, left) to maximize contact between the base tier and the membrane. K-Ras 1 to 4 are in contact with the membrane through the β2-β3 strands and the α5 helix. We refer the tier comprising K-Ras 1 to 4 as the base tier, and the four highest-numbered K-Ras proteins as the top tier. Except for the K-Ras proteins of the base and the top tiers, all K-Ras proteins in the helical assembly have essentially identical interactions with their neighboring K-Ras proteins. The helical assembly can grow in a straight tower shape, but we note that the base tier alone, without further assembly, is stable on the membrane in our simulations (Figure S3I).

We conjectured that the K-Ras helical assembly could form the core of a Ras-Raf signalosome. In the helical assembly, the entire RBD interface of each K-Ras protein is exposed and thus available for Raf recruitment. Further, the C and N termini of the K-Ras proteins are positioned outward at the surface of the assembly, leaving space for galectins to interact with the C-termini and for fluorescent proteins to fuse with the N termini. Such fusions have been used in electron microscopy experiments on Ras-Raf nanoclusters.^43^

A K-Ras protein at position *n* in our helical assembly model engages in multivalent interactions (Figure 4A), making contact with up to six other K-Ras proteins: GTP-mediated primary interactions with K-Ras *n*−1 and *n*+1, secondary (stacking) interactions with K-Ras *n*−4 and *n*+4 along the axis of the helical assembly, and tertiary interactions with K-Ras *n*−3 and *n*+3 (Figures 4C, S6A, and S6B). Each secondary and tertiary interaction has a buried surface area of ~1450 Å^2^ and ~660 Å^2^, and five and three salt bridges (Figures S6A and S6B), respectively; together these are larger than the corresponding primary interaction (~1750 Å^2^). All these interactions are asymmetric and predominantly polar.

**Figure 4.**
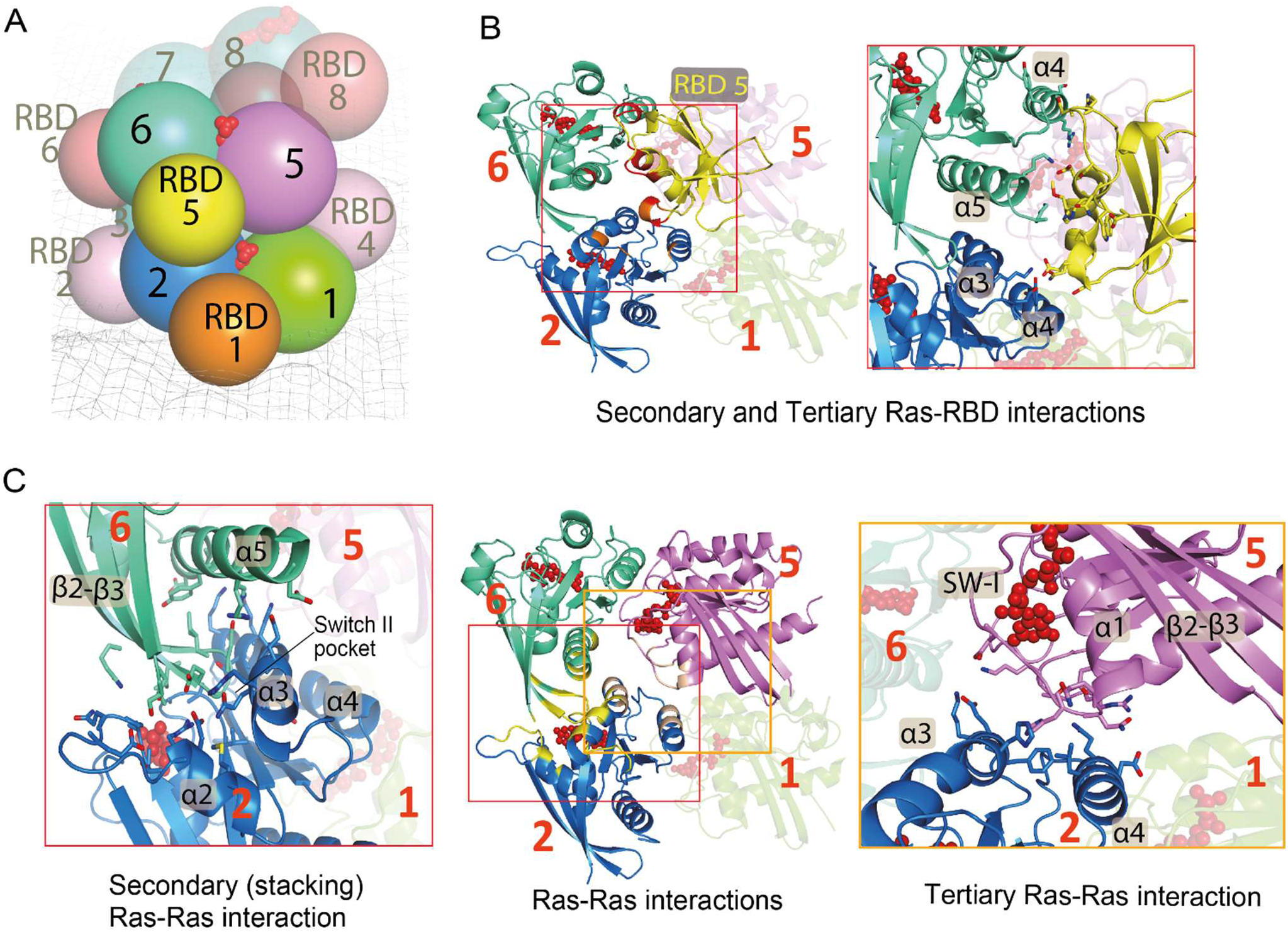
K-Ras/K-Ras and K-Ras/RBD interactions in the signalosome model. **A**. Cartoon representation of the relative positions of K-Ras proteins and RBD domains in the model; the membrane is shown as a mesh, and GTP in red. **B**. Left panel: the primary (with K-Ras 5), secondary (with K-Ras 6), and tertiary (with K-Ras 2) RBD-Ras interactions (of RBD 5); a fourth K-Ras (K-Ras 1) is shown for geometric context. The secondary and tertiary interface residues (and GTP) are colored in red and orange. Right panel: close-up of the Ras-RBD interactions. **C.** Center panel: three K-Ras proteins engaged in primary, secondary, and tertiary interactions with one another; the interface residues are shown in yellow and GTP in red. Left and right panels: close-ups of the secondary (stacking) and tertiary Ras-Ras interactions, respectively.

The secondary (stacking) interaction places the β2-β3 hairpin of K-Ras *n*+4 into the so-called “switch II pocket” of K-Ras *n*, which is exploited by covalent inhibitors of K-Ras G12C.^44^ Crystallographic data has shown that the switch II pocket can form without an occupying ligand (e.g., in PDB 4LDJ; Figure S6E), and the secondary dimer structure of K-Ras *n* and K-Ras *n+*4 to a large degree resembles a K-Ras dimer seen in crystal packing (e.g., in PDB 5UQW; Figure S6F). The G12C covalent inhibitors, by occupying the switch II pocket, are predicted to disrupt the stacking interaction (see SI), but the GMA dimer of inhibitor-bound G12C remained stable in simulations (Figure S2I). It is thus possible that the inhibitor binding is compatible with the GMA dimerization of G12C.

### Experimental validation of the helical assembly

To experimentally validate the K-Ras helical assembly, we first used negative-stain electron microscopy (EM) to image K-Ras particles in a reconstituted system. In these experiments, full-length K-Ras proteins were tethered to a lipid monolayer by Cys185, forming a chemical bond with a maleimide lipid. The K-Ras proteins were then applied to a grid, stained, and imaged (Figure 5H). To our knowledge, this is the first visualization of negatively stained Ras particles by electron microscopy. The K-Ras particles form in a maleimide-lipid dependent manner (Figures 5A–5D), consistent with the notion that K-Ras assembly requires K-Ras membrane localization. Classification of 73,282 visualized particles into 100 classes revealed a range of sizes (Figure 5E). The heterogeneity in particle shapes prevented reconstruction of the 3D structures. The size of the particles, however, is consistent with our helical assembly model. Consistent with the MEF, FRET (Figures 3F–3H), and BRET analysis (Figure 6D), the D154Q mutation disrupted particle formation (Figure 5F). The particle formation was also disrupted by K88D (Figure 5G), a mutation that disrupted the FRET and BRET signal of K-Ras assembly in cells (see Figures 6D and S5G). These mutagenesis data suggest that the helical assemblies underlie the K-Ras particles we visualized.

**Figure 5.**
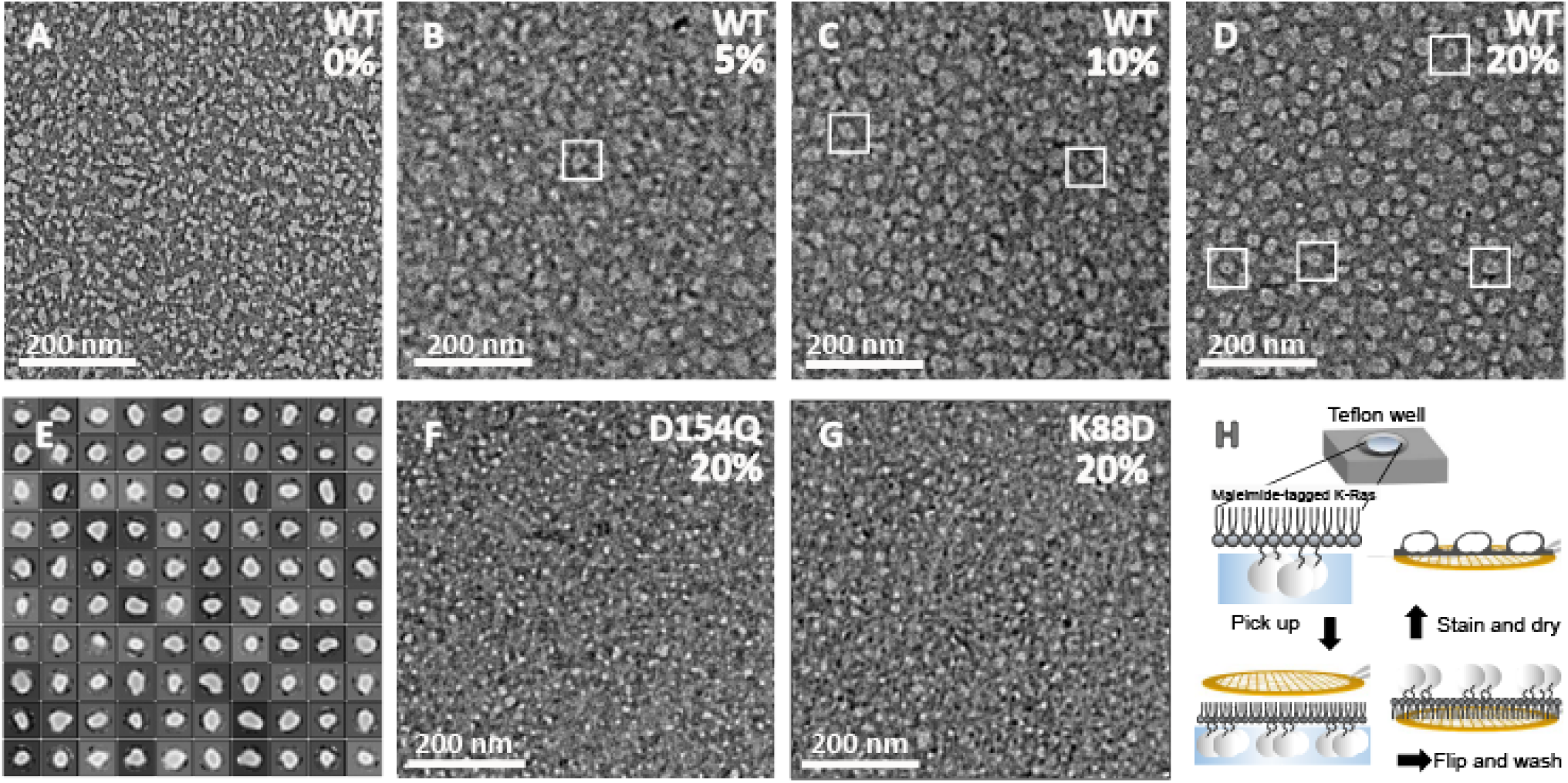
EM images with negative stain of K-Ras assembly. **A–D**. K-Ras wild type at 10 μM concentration; maleimide lipid percentage used is shown. **E**. 2D class average of the K-Ras wild-type particles. **F**. D154Q K-Ras at 10 μM concentration, 20% maleimide lipid. **G**. K88D K-Ras at 10 μM concentration, 20% maleimide lipid. **H**. Workflow of the experiment: After incubation with the monolayer lipid overnight, the K-Ras particles were moved to a carbon-coated copper grid, washed intensively, negatively stained, and examined by EM.

**Figure 6.**
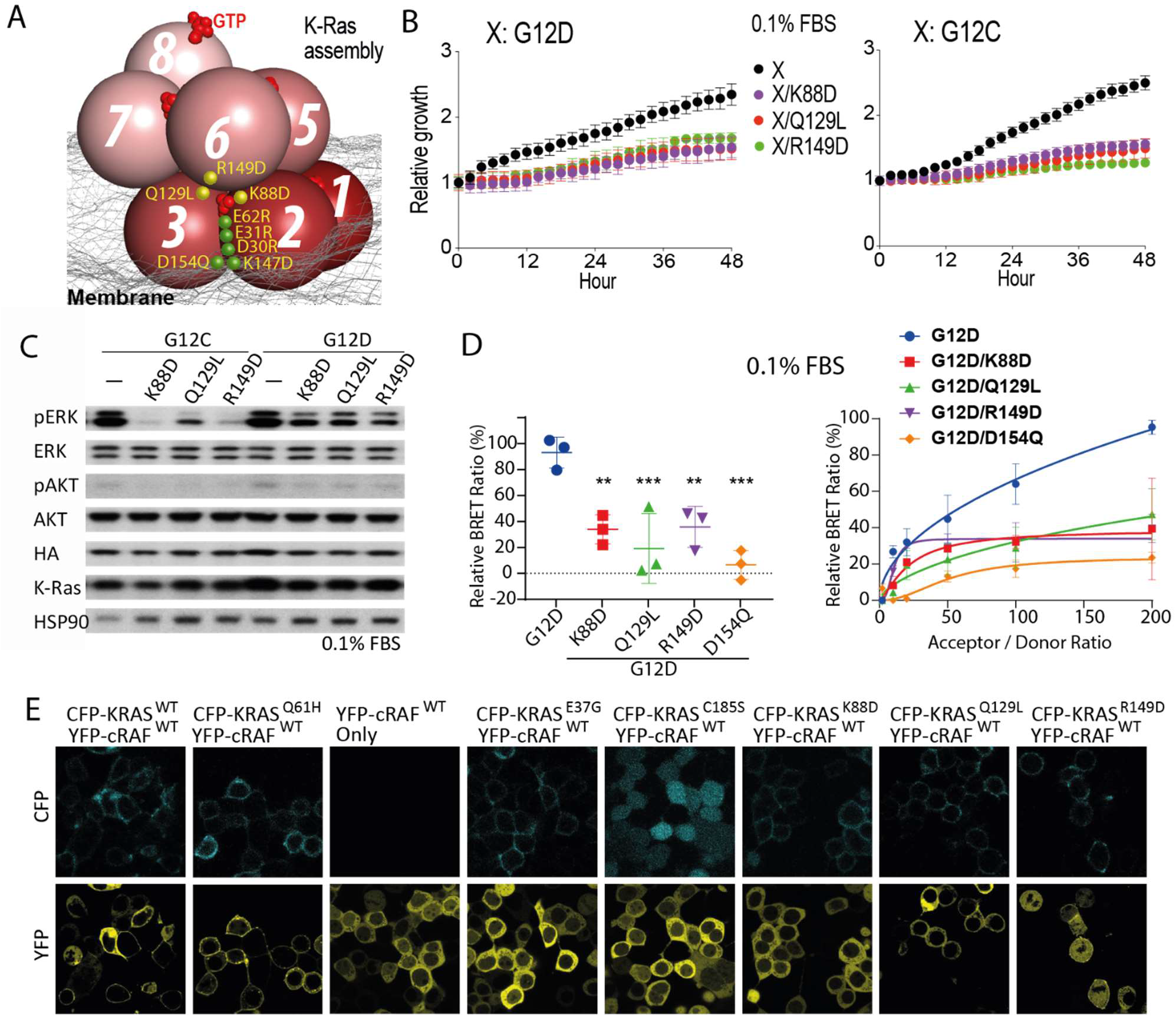
Mutagenesis validation of the secondary and tertiary K-Ras/K-Ras interactions. **A**. Illustration of the locations of the tested mutations, including those at the primary interface shown in Figure 3F–3H (green) and those at the secondary and tertiary interfaces (yellow). **B**. Proliferation rates of *K-Ras*^*lox*^/*K-RAS*^*MUT*^ cells of K88D, Q129L, and R149D in the background of G12C or G12D. Results are averages of three duplicates with error bars. Representative images at the end point are shown in Figure S4D. See Figure S4F for results obtained in 0.5% and 10% FBS. **C**. Phosphorylation of ERK and AKT in *K-Ras^lox^*/*K-RAS*^*MUT*^ cells. See Figure S4E for the phosphorylation in 0.5% and 10% FBS. **D**. Left panel: endpoint BRET measurements of K88D, Q129L, and R149D in the background of G12D, compared with the G12D mutation only; D154Q is included as a positive control. K-Ras construct (Residue 1–166) lacking the membrane-anchoring HVR was also used as a control (Figure S5F). The BRET signals were normalized to the maximum signals of G12D. An acceptor/donor expression plasmid ratio of 200:1 was chosen for comparing the K-Ras mutants. Right panel: co-transfection of increasing ratios of donor and acceptor plasmids, which allows discrimination between specific and unspecific (random collision) protein-protein interactions. **E**. Fluorescent imaging of Raf localization of K-Ras mutants K88D, Q129L, and R149D; K-Ras wild type and Q61H as positive controls, and E37G, C185S, and K-Ras-free cells as negative controls.

We also evaluated mutations predicted to disrupt the K-Ras helical assembly using the MEF, BRET and FRET systems. K88D, which was predicted to disrupt the secondary interface, and Q129L and R149D, which were predicted to disrupt the tertiary K-Ras/K-Ras interface (Figure 6A), slowed proliferation of the K-Ras dependent MEFs (Figures 6B and S4F) at low FBS and disrupted ERK phosphorylation in the MAPK pathway (Figures 6C and S4E). K88D also disrupted the FRET signal for K-Ras assembly, while Q129L and R149D did not (Figure S5G), possibly because the effect of Q129L and R149D mutations is more subtle: The GMA dimers would still form despite Q129L and R149D mutations, while K88D is located adjacent to the GMA dimer interface (although not part of the interface itself) and has the potential to disrupt the GMA dimer in addition the larger assembly (Figure 6A). To supplement the FRET experiment, we used BRET, which allows better quantification because of its larger Förster radius, to test these three mutations on the background of G12D. The BRET data confirmed that K88D, Q129L, and R149D mutations disrupt K-Ras assembly, and that their effect is weaker than D154Q at the GMA dimerization interface (Figure 6D). Although they are distal to the K-Ras interface with Raf, K88D, Q129L, and R149D mutations also hindered K-Ras recruitment of C-Raf to the cell membrane (Figure 6E), suggesting that the K-Ras assemblies play an important role in K-Ras/Raf interaction.

### C-Raf forms multivalent interactions with the K-Ras helical assembly

Using a co-crystal structure of the C-Raf RBD and H-Ras (PDB 4G0N)^45^ as a template, we docked a C-Raf RBD to each K-Ras of the helical assembly. The docked RBDs did not sterically clash with any K-Ras proteins of the helical assembly, nor with one another. The RBD that primarily interacts with K-Ras *n*, which we refer to as RBD *n*, forms a secondary contact with K-Ras *n*+1 and a tertiary contact with K-Ras *n*−3 (Figures 4B, S6C, and S6D). Together these secondary and tertiary Ras-RBD interfaces (~660 Å^2^ and ~550 Å^2^, respectively, in buried surface area) are nearly as large as the primary interface (~1600 Å^2^), and may thus confer an advantage on the helical assembly over a K-Ras monomer in terms of RBD binding. In turn, RBD binding may help stabilize the K-Ras assembly.

The 52-residue zinc-coordinated Raf CRD also interacts with Ras^46^ and imparts specificity to Ras-Raf binding.^47^ Whereas the RBD binds to GTP-bound Ras with nanomolar affinity,^48^ CRD binding is nucleotide-insensitive and weaker, with only micromolar affinity.^49^ Informed by extensive simulations of a CRD tethered to a structure of a K-Ras/RBD complex (see SI), we positioned the CRD at the C terminus of the α5 helix and β strands of K-Ras (Figure 7A), with an interface of ~1400 Å^2^. The CRD also contacts the switch II region of K-Ras (Figure S7F); this is consistent with switch II mutations (G60A and Y64W) hindering the CRD binding of H-Ras.^50^ NMR and other analyses have identified a set of CRD and Ras residues involved in the Ras-CRD interaction;^46,49,51,52^ these residues are either part of the modeled Ras-CRD interface or proximal to it (Figures S7F and S7G). By our model, a CRD interacts with a K-Ras and an adjacent RBD with extensive electrostatic complementarity (Figure S7H). We added a CRD to each K-Ras protein in the helical assembly. In the base tier, both the RBDs and CRDs interact extensively with the membrane (Figure 7A), and CRD residues Arg143, Lys144, and Lys148, interact extensively with POPS lipids (Figure S7I), consistent with previous studies.^53^

**Figure 7.**
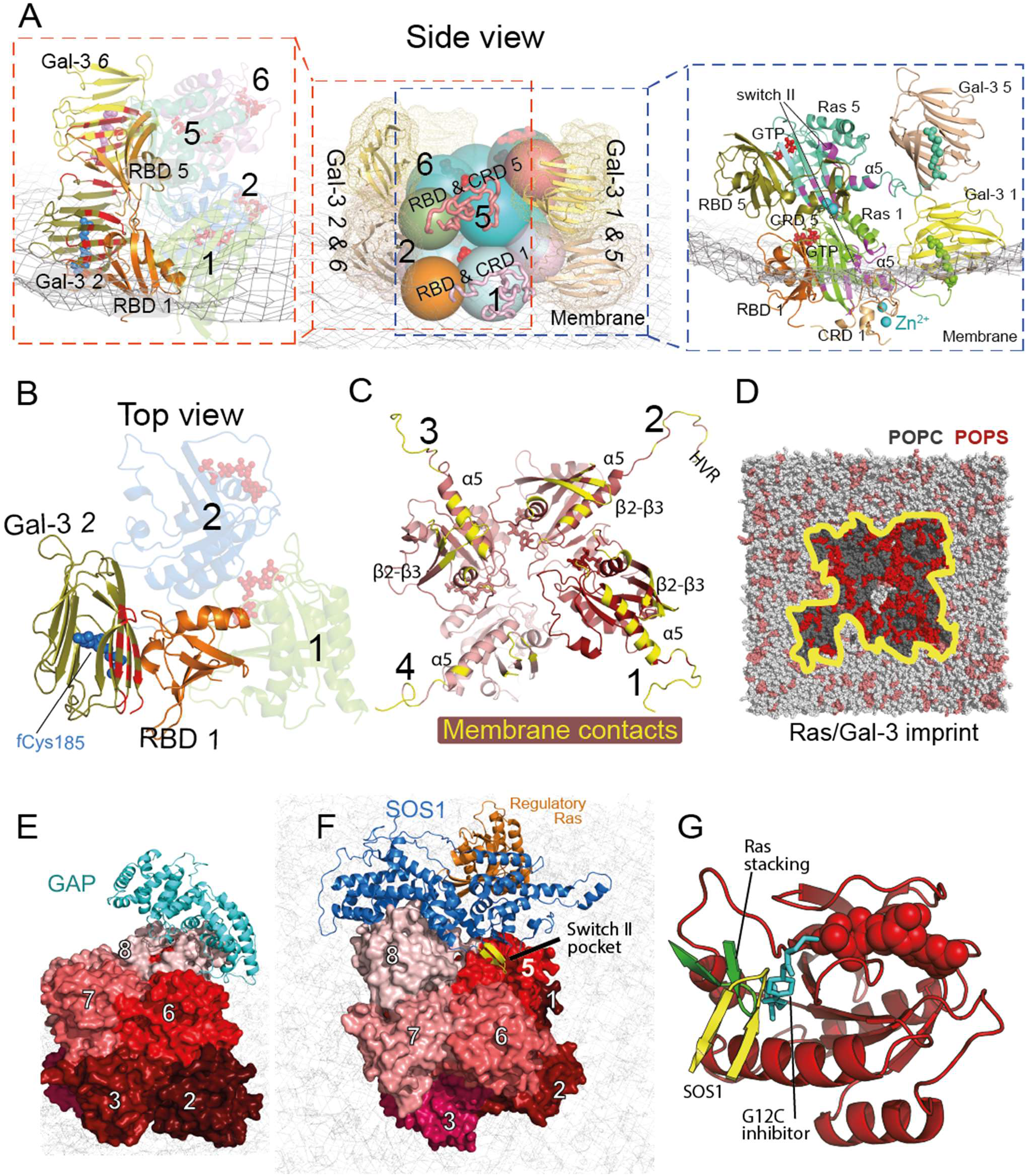
Ras interactions with Gal-3 and Raf RBD and CRD domains. **A**. Side views of the interactions of two stacking K-Ras proteins with Gal-3 and C-Raf RBD and CRD domains. Center panel: the relative positions of K-Ras, Gal-3, and Raf RBD and CRD domains. K-Ras and RBDs are shown as spheres, CRDs as ribbons, and Gal-3 proteins as ribbons surrounded by mesh. Left panel: the K-Ras proteins with RBDs and Gal-3 proteins in cartoon representation; the galectin-bound fCys185 residues of K-Ras are shown in blue and purple, and Gal-3–RBD interfaces are marked in red. Right panel: positions of CRD domains and Gal-3 proteins with respect to RBD-bound K-Ras. Ras-CRD interfaces are marked in purple. Also labeled is the α2 helix in the switch II region of K-Ras involved in the interaction with the CRD. **B**. Top view of the RBD interaction with Gal-3. Two K-Ras proteins, which form a GTP-mediated dimer, are shown as faded images. The fCys185 of K-Ras 2 is shown. **C**. K-Ras membrane-contacting residues (yellow) in the base tier. **D**. Membrane imprint of the K-Ras and Gal-3 proteins of the base tier, which is enriched in POPS lipids. **E**. GAP interacting with the tail K-Ras (K-Ras 8) of the helical assembly. **F**. SOS1 interacting with the tail K-Ras. In the secondary SOS1/K-Ras interaction the SOS1 helical hairpin (yellow) is inserted into the switch II pocket of K-Ras 5. **G**. Three interactions utilizing the switch II pocket: the secondary SOS1/K-Ras interaction, the K-Ras/K-Ras stacking involving the β2-β3 turn (green), and covalent G12C inhibitor binding (cyan).

### Gal-3 anchors the K-Ras helical assembly to the membrane and interacts with C-Raf

Although the mechanism of K-Ras membrane localization is commonly thought to be insertion of fCys185, the C-terminal tails of the helical assembly (with the exception of those in the base tier) do not contact the membrane, and thus their fCys185 residues would not readily be buried. Gal-3 can directly interact with the lipidated C-terminal tail, acting as a cap on the farnesyl group of fCys185 (and is essential for K-Ras nanoclustering,^22,54,55^ as is Gal-1 for H-Ras nanoclustering^23,56,57^). Since Gal-3 associates with membrane lipids^58^ and has been suggested to act as a so-called “prenyl receptor”^59^ to anchor K-Ras to the membrane, we considered a model in which all fCys185 residues, including those in the base tier, interact with Gal-3 proteins, and Gal-3 proteins interact with and localize the K-Ras assembly to the membrane. From unbiased MD simulations of ligand binding,^28^ we obtained a structural model of a farnesyl-bound Gal-3 (Figure S7D), in which the farnesyl is in a previously predicted^22^ hydrophobic cavity (Figure 7B). To bind fCys185, Gal-3 must be located near the K-Ras helical assembly, where limited space is available in the presence of C-Raf RBDs and CRDs.

The position of Gal-3 in this space (see SI) is tightly constrained by several considerations. First, at the base tier, the position of fCys185-bound Gal-3 proteins should allow the basic C-terminal tails to contact the membrane. Secondly, Gal-1 has been reported to interact with the C-Raf RBD, and this interface likely involves the RBD residue Asp117 and avoids the galectin carbohydrate-binding site;^60^ we assumed the same is true for Gal-3. Thirdly, both Gal-1^60^ and Gal-3^61^ dimerize, and they scaffold H-Ras^60^ and K-Ras^22,54^ nanoclustering, respectively. Finally, the Gal-3 dimerization interface should be consistent with results from mutagenesis studies.^61^

These considerations and extensive simulations (see SI) led us to a model in which Gal-3 *n* (the Gal-3 protein bound to the fCys185 residue of K-Ras *n*) interacts with RBD *n*−1 (with a buried surface area of ~1250 Å^2^; Figures 7A and 7B), including the Asp117 residue (Figure S7A). Any pair of Gal-3 *n* and *n*+4 in the model are stacked in parallel to the K-Ras stacking. The Gal-3 stacking, with an interface of ~670 Å^2^, is asymmetric, and the C-terminal tail of K-Ras *n*+4 is positioned near the interface, contacting both Gal-3 proteins (Figure 7A and S7B). As in the GMA dimer, K-Ras proteins in the base tier of the helical assembly adopt the α-like orientation, contacting the membrane with their α5 helices and β2-β3 hairpins (Figure 7C), and Gal-3 proteins interact with the membrane extensively (Figures 7A, S8A, and S8B), anchoring the K-Ras assembly. The C-Raf RBD and CRD domains in the base tier also interact with the membrane (Figures 7A and S8C), potentially contributing to the anchoring.

### A fuzzy structure for a Ras-Raf signalosome arises from disordered C-Raf linkers

In C-Raf activation in the MAPK pathway, a 14-3-3 dimer acts as a scaffold that binds to and stabilizes a C-Raf kinase domain dimer.^62^ C-Raf contains a loop C-terminal to the kinase domain that is phosphorylated at Ser621 and binds to 14-3-3 in C-Raf activation. There are multiple proteins in the 14-3-3 family; in the modeling, we incorporated 14-3-3σ into the Ras-Raf signalosome. Each C-Raf kinase domain also binds to and phosphorylates MEK1 kinase. As a step of modeling a Ras-Raf signalosome that includes full-length C-Raf, 14-3-3σ, and MEK1 proteins, we separately modeled the C-Raf kinase domain dimer complexed with a 14-3-3σ dimer and two MEK1 kinase domains (Figures 1F) using crystal structures^63–65^ that were available at the time we were constructing the model (Figure 1G) (see SI). The resulting model is overall highly consistent with cryo-EM structures of the 14-3-3 dimer in complex with a Raf kinase dimer (Figure S9A), which were later reported.^66,67^ A C-Raf protein contains a ~200-residue long, unstructured linker between its CRD and kinase domains. We modeled such linkers in order to connect the RBD and CRD domains to the kinase domains. Starting from a fully extended conformation, we simulated a linker for a total of 53 μs and obtained a diverse set of conformations with transient secondary structures (Figure S9E). We grafted a linker in a simulation-generated conformation onto each CRD in the model, then grafted onto each linker pair a C-Raf kinase domain dimer bound to 14-3-3σ and MEK1 (Figure 1F). Incorporating this model into the larger model then produced our full signalosome model.

In simulations of this full model, two paired C-Raf linkers developed extensive but nonspecific interactions with each other, with an interface of ~3400 Å^2^. The unstructured linkers introduce structural “fuzziness” to the signalosome, which is often seen in signalosome complexes^68^ and may, through the so-called “fly-casting” mechanism,^69^ produce a large capture radius for Ras-Raf and Raf-MEK interactions. Consistent with previous studies,^70,71^ the pSer338 and pTyr341 residues on the linkers developed favorable trans interactions with the αC helices of the C-Raf kinase domain dimer (Figures S9B–S9D), stabilizing the dimer. Although other pairing patterns are possible in principle, we adopted a *n*/*n+*4 Raf pairing, by which Raf *n* consistently dimerizes with Raf *n+*4 for our signalosome model (Figure 1E). In a 100-μs simulation, the 8-protomer signalosome, especially its core of K-Ras, remained stable (Figure S1B), and developed extensive interactions with the membrane, with an interface area of ~472,000 Å^2^. Many basic residues in the signalosome formed ionic interactions with the membrane, enriching negatively charged POPS lipids in the membrane at the interface (Figure 7D). This is consistent with indications that POPS depletion from the plasma membrane reduces nanoclustering of GTP-bound K-Ras.^72^

### The modeled Ras-Raf signalosome allows for regulation by GEFs and GAPs

GAPs, such as RasGAP, catalyze the hydrolysis of GTP in Ras to revert it to its GDP-bound state.^7^ Ras nanoclustering has been suggested to attenuate RasGAP-catalyzed GTP hydrolysis^54,55^ and to sequester Ras from GAP binding.^22,56^ Consistent with these results, RasGAP is sterically prevented from interacting with all but the tail K-Ras protein in the helical assembly. Based on an H-Ras/p120GAP structure,^32^ GAP can dock to a tail K-Ras protein without steric clashes, provided that the K-Ras protein is not bound to Raf (Figure 7E). GAP can then convert the tail K-Ras protein to the GDP-bound state, preventing any additional K-Raf protein from joining the helical assembly and promoting dissolution of the signalosome. Compared to the other K-Ras molecules in the signalosome, the tail K-Ras is most exposed and likely the least stable. We thus anticipate that under GAP regulation, a signalosome will eventually disassemble by one tail-position K-Ras falling off at a time.

Based on a structure of H-Ras in complex with the GEF protein SOS1,^73^ a SOS1 protein in active conformation can also dock to the tail K-Ras protein (Figure 7F). By ensuring that the tail K-Ras protein is in the GTP-bound state, ready to receive another K-Ras protein into the signalosome, SOS1 may help maintain and grow the signalosome. SOS1 is activated in part by an allosteric GTP-bound Ras protein,^74^ which our model accommodates. The model predicts, moreover, that if an active SOS1 protein docks to the tail K-Ras protein *n*, it additionally engages K-Ras *n*−3 in a secondary interaction in which a helical hairpin of SOS1 is inserted into the switch II pocket of K-Ras *n*−3 (Figures 7F and 7G). This secondary Ras-SOS1 interaction may contribute to the specificity of SOS1 in regulating MAPK signaling. One consequence of this SOS1 interaction is that it may place an upper limit on the height (and thus overall size) of the signalosome, since SOS1 is membrane-anchored by its PH domain, and thus has only limited “reach” from the membrane.

## Discussion

The Ras-Raf signalosome model we propose here entails a host of multivalent interactions, whereby a K-Ras protein may interact with up to six others simultaneously, and a Raf RBD may interact with three K-Ras proteins and one Gal-3 protein. Although many of the pairwise protein-protein interactions in the signalosome—including the GTP-mediated dimerization of K-Ras—are likely to be weak, these cooperative interactions should, by a nucleation-like process,^75^ produce a signalosome that stabilizes the active state of K-Ras, provides favorable composite interfaces for Raf recruitment, and sequesters K-Ras from down-regulation, thereby yielding a switch-like and sustained response to input signals.

It is well known that membrane localization is required for Ras activity, and furthermore that lipid composition and the local structure of the membrane also significantly affect Ras clustering.^43^ Indeed, the presence of the membrane is essential for its role in promoting GMA dimerization (by raising the local concentration of K-Ras and restricting its orientation in favor of dimerization) and thus signalosome formation, even though K-Ras farnesyl groups are not buried in the membrane in our model.

Proteins downstream of Ras may also affect signalosome formation. Certain small-molecule inhibitors of B-Raf give rise to so-called “paradoxical activation” of Raf by inducing C-Raf/B-Raf heterodimerization and even C-Raf homodimerization,^64^ and also promote Ras nanoclustering by inducing more, but not larger, Ras nanoclusters in cells.^76^ Consistent with this surprising B-Raf inhibitor effect, our model suggests a scenario in which enhanced Raf dimerization promotes nucleus formation and in turn, signalosome formation.

The K-Ras helical assembly provides a possible explanation for why wild-type K-Ras acts as a tumor suppressor in cells driven by oncogenic K-Ras mutants.^77,78^ Wild-type K-Ras, which is predominantly GDP-bound and cannot serve as a GTP donor, may inhibit the growth of a helical assembly by capping it at the tail position. This scenario is consistent with the D154Q K-Ras mutant being inactive as a tumor suppressor,^19^ since the D154Q mutation at the acceptor interface may prevent the mutant from assuming the tail position. This rationale leads us to predict that mutations at the donor interface (e.g., K147D and R161E) should not disrupt the tumor suppressor activity, as these mutants should still be able to assume the tail position and cap helical assemblies of oncogenic K-Ras mutants.

We do not expect the signalosome model to be relevant to all Ras signaling pathways, which may use a variety of other structural mechanisms. The GMA K-Ras dimer model and, by extension, the K-Ras helical assembly are almost certainly inapplicable to PI3K activation (see SI). Lys128 and Arg135, two key residues at the putative GMA dimer interface, are conserved in H-Ras and N-Ras, but overall in only 6 of the 13 human Ras isoforms,^40^ and homology analysis (Figure S2J) shows that these two residues are not well conserved in evolution, suggesting that the signalosome structure is not general to small G-proteins. Lys128 and Arg135, however, are conserved in mammals, and the signalosome model is likely to be broadly relevant to mammalian MAPK signaling. In many respects, findings on H-Ras nanoclustering echo those on K-Ras,^43^ and our signalosome model is consistent with the structures of other Ras and Raf proteins (such as N-Ras, H-Ras, A-Raf, and B-Raf) involved in MAPK signaling. With local alterations, our model may be extendable to those Ras and Raf proteins and provide a framework for understanding their overlapping, yet distinct roles in MAPK signaling.

## Acknowledgments

The authors thank Michael Eastwood for helpful discussions, Pelin Ayaz for a critical reading of the manuscript, Kevin Yuh for assistance with movies, Jessica McGillen and Berkman Frank for editorial assistance, and Claudia Scholl, Jonas V. Schaefer, and Jonas Binz for their support with the BRET experiments. This work was in supported in part by a Stand Up To Cancer (SU2C)–American Cancer Society Lung Cancer Dream Team Translational Research Grant (SU2C-AACR-DT17-15 to P.A.J.), the Gadzooks Fund (to P.A.J.), the Cammarata Family Foundation Research Fund (to P.A.J.), the Giovanni Armenise–Harvard Foundation and the Lung Cancer Research Foundation (to C.A.), the US Department of Defense (W81XWH-16-1-0106 to K.D.W.), Hale Center and Pinard Family (DFCI to K.D.W.), the Cancer Prevention and Research Institute of Texas (RP170373 to K.D.W.), and the Krebsliga Schweiz grant (KFS 4147-02-2017 to A.P.). SU2C is a program of the Entertainment Industry Foundation. Research grants are administered by the American Association for Cancer Research, the scientific partner of SU2C. P.A.J. has served as a consultant for and has received sponsored research funding from AstraZeneca, Mirati Therapeutics, and Araxes Pharmaceuticals.

## References

1. Wennerberg, K., Rossman, K.L., and Der, C.J. (2005). The Ras superfamily at a glance. J. Cell. Sci. 118(5), 843–846.

2. Hobbs, G.A., Der, C.J., and Rossman, K.L. (2016). RAS isoforms and mutations in cancer at a glance. J. Cell. Sci. 129(7), 1287–1292.

3. Yuan, T.L., Amzallag, A., Bagni, R., Yi, M., Afghani, S., Burgan, W., Fer, N., Strathern, L.A., Powell, K., Smith, B., Waters, A.M., Drubin, D., Thomson, T., Liao, R., Greninger, P., Stein, G.T., Murchie, E., Cortez, E., Egan, R.K., Procter, L., Bess, M., Cheng, K.T., Lee, C.-S., Lee, L.C., Fellmann, C., Stephens, R., Luo, J., Lowe, S.W., Benes, C.H., and McCormick, F. (2018). Differential effector engagement by oncogenic KRAS. Cell Rep. 22(7), 1889–1902.

4. Tidyman, W.E. and Rauen, K.A. (2009). The RASopathies: developmental syndromes of Ras/MAPK pathway dysregulation. Curr. Opin. Genetics Dev. 19(3), 230–236.

5. Prior, I.A., Lewis, P.D., and Mattos, C. (2012). A comprehensive survey of Ras mutations in cancer. Cancer Res. 72(10), 2457–2467.

6. Simanshu, D.K., Nissley, D.V., and McCormick, F. (2017). RAS proteins and their regulators in human disease. Cell 170(1), 17–33.

7. Cherfils, J. and Zeghouf, M. (2013). Regulation of small GTPases by GEFs, GAPs, and GDIs. Physiol. Rev. 93(1), 269–309.

8. Wittinghofer, A. and Pal, E.F. (1991). The structure of Ras protein: a model for a universal molecular switch. Trends Biochem. Sci. 16, 382–387.

9. Wittinghofer, A. and Vetter, I.R. (2011). Structure-function relationships of the G domain, a canonical switch motif. Annu. Rev. Biochem. 80(1), 943–971.

10. Santos, E., Nebreda, A.R., Bryan, T., and Kempner, E.S. (1988). Oligomeric structure of p21 ras proteins as determined by radiation inactivation. J. Biol. Chem. 263(20), 9853–9858.

11. Dementiev, A. (2012). K-Ras4B lipoprotein synthesis: biochemical characterization, functional properties, and dimer formation. Protein Expr. Purif. 84(1), 86–93.

12. Chen, M., Peters, A., Huang, T., and Nan, X. (2016). Ras dimer formation as a new signaling mechanism and potential cancer therapeutic target. Mini-Rev. Med. Chem. 16(5), 391–403.

13. Güldenhaupt, J., Rudack, T., Bachler, P., Mann, D., Triola, G., Waldmann, H., Kötting, C., and Gerwert, K. (2012). N-Ras forms dimers at POPC membranes. Biophys. J. 103(7), 1585–1593.

14. Werkmüller, A., Triola, G., Waldmann, H., and Winter, R. (2013). Rotational and translational dynamics of Ras proteins upon binding to model membrane systems. ChemPhysChem 14(16), 3698–3705.

15. Muratcioglu, S., Chavan, T.S., Freed, B.C., Jang, H., Khavrutskii, L., Freed, R.N., Dyba, M.A., Stefanisko, K., Tarasov, S.G., Gursoy, A., Keskin, O., Tarasova, N.I., Gaponenko, V., and Nussinov, R. (2015). GTP-dependent K-Ras dimerization. Structure 23(7), 1325–1335.

16. Inouye, K., Mizutani, S., Koide, H., and Kaziro, Y. (2000). Formation of the Ras dimer is essential for Raf-1 activation. J. Biol. Chem. 275(6), 3737–3740.

17. Nan, X., Tamgüney, T.M., Collisson, E.A., Lin, L.J., Pitt, C., Galeas, J., Lewis, S., Gray, J.W., McCormick, F, and Chu, S. (2015). Ras-GTP dimers activate the mitogen-activated protein kinase (MAPK) pathway. Proc. Natl. Acad. Sci. U.S.A. 112(26), 7996–8001.

18. Spencer-Smith, R., Koide, A., Zhou, Y., Eguchi, R.R., Sha, F., Gajwani, P., Santana, D., Gupta, A., Jacobs, M., Herrero-Garcia, E., Cobbert, J., Lavoie, H., Smith, M., Rajakulendran, T., Dowdell, E., Okur, M.N., Dementieva, I., Sicheri, F., Therrien, M., Hancock, J.F., Ikura, M., Koide, S., and O’Bryan, J.P. (2017). Inhibition of RAS function through targeting an allosteric regulatory site. Nat. Chem. Biol. 13(1), 62–68.

19. Ambrogio, C., Köhler, J., Zhou, Z.-W., Wang, H., Paranal, R., Li, J., Capelletti, M., Caffarra, C., Li, S., Lv, Q., Gondi, S., Hunter, J.C., Lu, J., Chiarle, R., Santamaría, D., Westover, K.D., and Jänne, P.A. (2018). KRAS dimerization impacts MEK inhibitor sensitivity and oncogenic activity of mutant KRAS. Cell 172(4), 857–868.

20. Plowman, S.J., Muncke, C., Parton, R.G., and Hancock, J.F. (2005). H-ras, K-ras, and inner plasma membrane raft proteins operate in nanoclusters with differential dependence on the actin cytoskeleton. Proc. Natl. Acad. Sci. U.S.A. 102(43), 15500–15505.

21. Tian, T., Harding, A., Inder, K., Plowman, S., Parton, R.G., and Hancock, J.F. (2007). Plasma membrane nanoswitches generate high-fidelity Ras signal transduction. Nat. Cell Biol. 9(8), 905–914.

22. Shalom-Feuerstein, R., Plowman, S.J., Rotblat, B., Ariotti, N., Tian, T., Hancock, J.F., and Kloog, Y. (2008). K-Ras nanoclustering is subverted by overexpression of the scaffold protein Galectin-3. Cancer Res. 68(16), 6608–6616.

23. Belanis, L., Plowman, S.J., Rotblat, B., Hancock, J.F., and Kloog, Y. (2008). Galectin-1 is a novel structural component and a major regulator of H-Ras nanoclusters. Mol. Biol. Cell 19(4), 1404–1414.

24. Plowman, S.J., Ariotti, N., Goodall, A., Parton, R.G., and Hancock, J.F. (2008). Electrostatic interactions positively regulate K-Ras nanocluster formation and function. Mol. Cell. Biol. 28(13), 4377–4385.

25. Sutton MN, Lu Z, Li YC, Zhou Y, Huang T, Reger A, Hurwitz AM, Palzkill T, Logsdon C, Liang X, Gray JW, Nan X, Hancock J, Wahl GM, Bast RC Jr. (2019) DIRAS3 (ARHI) blocks RAS/MAPK signaling by binding directly to RAS and disrupting RAS clusters. Cell Rep. 29(11), 3448–3459.

26. Wu, H. (2013). Higher-order assemblies in a new paradigm of signal transduction. Cell 153(2), 287–292.

27. Rajakulendran, T., Sahmi, M., Lefrançois, M., Sicheri, F., and Therrien, M. (2009). A dimerization-dependent mechanism drives RAF catalytic activation. Nature 461(7263), 542–545.

28. Shan, Y., Kim, E.T., Eastwood, M.P., Dror, R.O., Seeliger, M.A., and Shaw, D.E. (2011). How does a drug molecule find its target binding site? J. Am. Chem. Soc. 133(24), 9181–9183.

29. Shan, Y., Gnanasambandan, K., Ungureanu, D., Kim, E.T., Hammarén, H., Yamashita, K., Silvennoinen, O., Shaw, D.E., and Hubbard, S.R. (2014). Molecular basis for pseudokinase-dependent autoinhibition of JAK2 tyrosine kinase. Nat. Struct. Mol. Biol. 21(7), 579–584.

30. Plattner, N., Doerr, S., De Fabritiis, G., and Noé, F. (2017). Complete protein-protein association kinetics in atomic detail revealed by molecular dynamics simulations and Markov modelling. Nat. Chem. 9(10), 1005–1011.

31. Wright LP, Philips MR. (2006) Thematic review series: lipid posttranslational modifications. CAAX modification and membrane targeting of Ras. J. Lipid Res. 47(5), 883–891.

32. Scheffzek, K., Ahmadian, M.R., Kabsch, W., Wiesmüller, L., Lautwein, A., Schmitz, F., and Wittinghofer, A. (1997). The Ras-RasGAP complex: structural basis for GTPase activation and its loss in oncogenic Ras mutants. Science 277(5324), 333–338.

33. Ahmadian, M.R., Stege, P., Scheffzek, K., and Wittinghofer, A. (1997). Confirmation of the arginine-finger hypothesis for the GAP-stimulated GTP-hydrolysis reaction of Ras. Nat. Struct. Biol. 4(9), 686–689.

34. Vetter, I.R. and Wittinghofer, A. (2001). The guanine nucleotide-binding switch in three dimensions. Science 294(5545), 1299–1304.

35. Geyer, M., Schweins, T., Herrmann, C., Prisner, T., Wittinghofer, A., and Kalbitzer, H.R. (1996). Conformational transitions in p21^ras^ and in its complexes with the effector protein Raf-RBD and the GTPase activating protein GAP. Biochemistry 35(32), 10308–10320.

36. Ito, Y., Yamasaki, K., Iwahara, J., Terada, T., Kamiya, A., Shirouzu, M., Muto, Y., Kawai, G., Yokoyama, S., Laue, E.D., Wälchli, M., Shibata, T., Nishimura, S., and Miyazawa, T. (1997). Regional polysterism in the GTP-bound form of the human c-Ha-Ras protein. Biochemistry 36(30), 9109–9119.

37. Spoerner, M., Herrmann, C., Vetter, I.R., Kalbitzer, H.R., and Wittinghofer, A. (2001). Dynamic properties of the Ras switch I region and its importance for binding to effectors. Proc. Natl. Acad. Sci. U.S.A. 98(9), 4944–4949.

38. Araki, M., Shima, F., Yoshikawa, Y., Muraoka, S., Ijiri, Y., Nagahara, Y., Shirono, T., Kataoka, T., and Tamura, A. (2011). Solution structure of the state 1 conformer of GTP-bound H-Ras protein and distinct dynamic properties between the state 1 and state 2 conformers. J. Biol. Chem. 286(45), 39644–39653.

39. Mazhab-Jafari, M.T., Marshall, C.B., Smith, M.J., Gasmi-Seabrook, G.M., Stathopulos, P.B., Inagaki, F., Kay, L.E., Neel, B.G., and Ikura, M. (2015). Oncogenic and RASopathy-associated K-RAS mutations relieve membrane-dependent occlusion of the effector-binding site. Proc. Natl. Acad. Sci. U.S.A. 112(21), 6625–6630.

40. Abankwa, D., Hanzal-Bayer, M., Ariotti, N., Plowman, S.J., Gorfe, A.A., Parton, R.G., McCammon, J.A., and Hancock, J.F. (2008). A novel switch region regulates H-ras membrane orientation and signal output. EMBO J. 27(5), 727–735.

41. Drosten, M., Dhawahir, A., Sum, E.Y., Urosevic, J., Lechuga, C.G., Esteban, L.M., Castellano, E., Guerra, C., Santos, E., and Barbacid, M. (2010). Genetic analysis of Ras signalling pathways in cell proliferation, migration and survival. EMBO J. 29(6), 1091–1104.

42. Sung, Y.J., Carter, M., Zhong, J.M., and Hwang, Y.W. (1995). Mutagenesis of the H-ras p21 at glycine-60 residue disrupts GTP-induced conformational change. Biochemistry 34(10), 3470–3477.

43. Zhou, Y. and Hancock, J.F. (2015). Ras nanoclusters: versatile lipid-based signaling platforms. Biochim. Biophys. Acta 1853(4), 841–849.

44. Ostrem, J.M., Peters, U., Sos, M.L., Wells, J.A., and Shokat, K.M. (2013). K-Ras (G12C) inhibitors allosterically control GTP affinity and effector interactions. Nature 503(7477), 548–551.

45. Fetics, S.K., Guterres, H., Kearney, B.M., Buhrman, G., Ma, B., Nussinov, R., and Mattos, C. (2015). Allosteric effects of the oncogenic RasQ61L mutant on Raf-RBD. Structure 23(3), 505–516.

46. Hu, C.-D., Kariya, K.-I., Tamada, M., Akasaka, K., Shirouzu, M., Yokoyama, S., and Kataoka, T. (1995). Cysteine-rich region of Raf-1 interacts with activator domain of post-translationally modified Ha-Ras. J. Biol. Chem. 270(51), 30274–30277.

47. Okada, T., Hu, C.-D., Jin, T.-G., Kariya, K.-I., Yamawaki-Kataoka, Y., and Kataoka, T. (1999). The strength of interaction at the Raf cysteine-rich domain is a critical determinant of response of Raf to Ras family small GTPases. Mol. Cell Biol. 19(9), 6057–6064.

48. Nakhaeizadeh, H., Amin, E., Nakhaei-Rad, S., Dvorsky, R., and Ahmadian, M.R. (2016). The RAS-effector interface: isoform-specific differences in the effector binding regions. PLoS One 11(12), e0167145.

49. Williams, J.G., Drugan, J.K., Yi, G.-S., Clark, G.J., Der, C.J., and Campbell, S.L. (2000). Elucidation of binding determinants and functional consequences of Ras/Raf-cysteine-rich domain interactions. J. Biol. Chem. 275(29), 22172–22179.

50. Drugan, J.K., Khosravi-Far, R., White, M.A., Der, C.J., Sung, Y.-J., Hwang, Y.-W., and Campbell, S.L. (1996). Ras interaction with two distinct binding domains in Raf-1 may be required for Ras transformation. J. Biol. Chem. 271(1), 233–237.

51. Winkler, D.G., Cutler, R.E. Jr., Drugan, J.K., Campbell, S., Morrison, D.K., and Cooper, J.A. (1998). Identification of residues in the cysteine-rich domain of Raf-1 that control Ras binding and Raf-1 activity. J. Biol. Chem. 273(34), 21578–21584.

52. Thapar, R., Williams, J.G., and Campbell, S.L. (2004). NMR characterization of full-length farnesylated and non-farnesylated H-Ras and its implications for Raf activation. J. Mol. Biol. 343(5), 1391–1408.

53. Improta-Brears T, Ghosh S, Bell RM. (1999) Mutational analysis of Raf-1 cysteine rich domain: requirement for a cluster of basic aminoacids for interaction with phosphatidylserine. Mol. Cell Biochem. 198(1–2), 171–178.

54. Elad-Sfadia, G., Haklai, R., Balan, E., and Kloog, Y. (2004). Galectin-3 augments K-Ras activation and triggers a Ras signal that attenuates ERK but not phosphoinositide 3-kinase activity. J. Biol. Chem. 279(33), 34922–34930.

55. Ashery, U., Yizhar, O., Rotblat, B., Elad-Sfadia, G., Barkan, B., Haklai, R., and Kloog, Y. (2006). Spatiotemporal organization of Ras signaling: rasosomes and the galectin switch. Cell. Mol. Neurobiol. 26(4–6), 469–493.

56. Paz, A., Haklai, R., Elad-Sfadia, G., Ballan, E., and Kloog, Y. (2001). Galectin-1 binds oncogenic H-Ras to mediate Ras membrane anchorage and cell transformation. Oncogene 20(51), 7486–7493.

57. Rotblat, B., Niv, H., André, S., Kaltner, H., Gabius, H.-J., and Kloog, Y. (2004). Galectin-1 (L11A) predicted from a computed galectin-1 farnesyl-binding pocket selectively inhibits Ras-GTP. Cancer Res. 64(9), 3112–3118.

58. Lukyanov, P., Furtak, V., and Ochieng, J. (2005). Galectin-3 interacts with membrane lipids and penetrates the lipid bilayer. Biochem. Biophys. Res. Commun. 338(2), 1031–1036.

59. Marshall, C.J. (1993). Protein prenylation: a mediator of protein-protein interactions. Science 259(5103), 1865–1867.

60. Blaževitš, O., Mideksa, Y.G., Šolman, M., Ligabue, A., Ariotti, N., Nakhaeizadeh, H., Fansa, E.K., Papageorgiou, A.C., Wittinghofer, A., Ahmadian, M.R., and Abankwa, D. (2016). Galectin-1 dimers can scaffold Raf-effectors to increase H-ras nanoclustering. Sci. Rep. 6, 24165.

61. Yang, R.-Y., Hill, P.N., Hsu, D.K., and Liu, F.-T. (1998). Role of the carboxyl-terminal lectin domain in self-association of galectin-3. Biochemistry 37(12), 4086–4092.

62. Tzivion, G., Luo, Z., and Avruch, J. (1998). A dimeric 14-3-3 protein is an essential cofactor for Raf kinase activity. Nature 394(6688), 88–92.

63. Molzan, M., Kasper, S., Röglin, L., Skwarczynska, M., Sassa, T., Inoue, T., Breitenbuecher, F., Ohkanda, J., Kato, N., Schuler, M., and Ottmann, C. (2013). Stabilization of physical RAF/14-3-3 interaction by cotylenin A as treatment strategy for RAS mutant cancers. ACS Chem. Biol. 8(9), 1869–1875.

64. Hatzivassiliou, G., Song, K., Yen, I., Brandhuber, B.J., Anderson, D.J., Alvarado, R., Ludlam, M.J.C., Stokoe, D., Gloor, S.L., Vigers, G., Morales, T., Aliagas, I., Liu, B., Sideris, S., Hoeflich, K.P., Jaiswal, B.S., Seshagiri, S., Koeppen, H., Belvin, M., Friedman, L.S., and Malek, S. (2010). RAF inhibitors prime wild-type RAF to activate the MAPK pathway and enhance growth. Nature 464(7287), 431–435.

65. Haling, J.R., Sudhamsu, J., Yen, I., Sideris, S., Sandoval, W., Phung, W., Bravo, B.J., Giannetti, A.M., Peck, A., Masselot, A., Morales, T., Smith, D., Brandhuber, B.J., Hymowitz, S.G., and Malek, S. (2014). Structure of the BRAF-MEK complex reveals a kinase activity independent role for BRAF in MAPK signaling. Cancer Cell 26(3), 402–413.

66. Park E, Rawson S, Li K, Kim BW, Ficarro SB, Pino GG, Sharif H, Marto JA, Jeon H, Eck MJ. (2019) Architecture of autoinhibited and active BRAF-MEK1-14-3-3 complexes. Nature. 575(7783), 545–550.

67. Kondo Y, Ognjenović J, Banerjee S, Karandur D, Merk A, Kulhanek K, Wong K, Roose JP, Subramaniam S, Kuriyan J. (2019) Cryo-EM structure of a dimeric B-Raf:14-3-3 complex reveals asymmetry in the active sites of B-Raf kinases. Science. 366(6461), 109–115.

68. Wu, H. and Fuxreiter, M. (2016). The structure and dynamics of higher-order assemblies: amyloids, signalosomes, and granules. Cell 165(5), 1055–1066.

69. Shoemaker, B.A., Portman, J.J., and Wolynes, P.G. (2000). Speeding molecular recognition by using the folding funnel: the fly-casting mechanism. Proc. Natl. Acad. Sci. U.S.A. 97(16), 8868–8873.

70. Mason, C.S., Springer, C.J., Cooper, R.G., Superti-Furga, G., Marshall, C.J., and Marais, R. (1999). Serine and tyrosine phosphorylations cooperate in Raf-1, but not B-Raf activation. EMBO J. 18(8), 2137–2148.

71. Jambrina, P.G., Rauch, N., Pilkington, R., Rybakova, K., Nguyen, L.K., Kholodenko, B.N., Buchete, N.-V., Kolch, W., and Rosta, E. (2016). Phosphorylation of RAF kinase dimers drives conformational changes that facilitate transactivation. Angew. Chem. 55(3), 983–986.

72. Zhou, Y., Liang, H., Rodkey, T., Ariotti, N., Parton, R.G., and Hancock, J.F. (2014). Signal integration by lipid-mediated spatial cross talk between Ras nanoclusters. Mol. Cell. Biol. 34(5), 862–876.

73. Boriack-Sjodin, P.A., Margarit, S.M., Bar-Sagi, D., and Kuriyan, J. (1998). The structural basis of the activation of Ras by SOS. Nature 394(6691), 337–343.

74. Margarit, S.M., Sondermann, H., Hall, B.E., Nagar, B., Hoelz, A., Pirruccello, M., Bar-Sagi, D., and Kuriyan, J. (2003). Structural evidence for feedback activation by Ras·GTP of the Ras-specific nucleotide exchange factor SOS. Cell 112(5), 685–695.

75. Li, P., Banjade, S., Cheng, H.-C., Kim, S., Chen, B., Guo, L., Llaguno, M., Hollingsworth, J.V., King, D.S., Banani, S.F., Russo, P.S., Jiang, Q.-X., Nixon, B.T., and Rosen, M.K. (2012). Phase transitions in the assembly of multivalent signalling proteins. Nature 483(7389), 336–340.

76. Cho, K.-J., Kasai, R.S., Park, J.-H., Chigurupati, S., Heidorn, S.J., van der Hoeven, D., Plowman, S.J., Kusumi, A., Marais, R., and Hancock, J.F. (2012). Raf inhibitors target Ras spatiotemporal dynamics. Curr. Biol. 22(11), 945–955.

77. Zhang, Z., Wang, Y., Vikis, H.G., Johnson, L., Liu, G., Li, J., Anderson, M.W., Sills, R.C., Hong, H.L., Devereux, T.R., Jacks, T., Guan, K.-L., and You, M. (2001). Wildtype Kras2 can inhibit lung carcinogenesis in mice. Nat. Genet. 29(1), 25–33.

78. Singh, A., Sowjanya, A.P., and Ramakrishna, G. (2005). The wild-type Ras: road ahead. FASEB J. 19(2), 161–169.

79. Maurer, T., Garrenton, L.S., Oh, A., Pitts, K., Anderson, D.J., Skelton, N.J., Fauber, B.P., Pan, B., Malek, S., Stokoe, D., Ludlam, M.J.C., Bowman, K.K., Wu, J., Giannetti, A.M., Starovasnik, M.A., Mellman, I., Jackson, P.K., Rudolph, J., Wang, W., and Fang, G. (2012). Small-molecule ligands bind to a distinct pocket in Ras and inhibit SOS-mediated nucleotide exchange activity. Proc. Natl. Acad. Sci. U.S.A. 109(14), 5299–5304.

80. Mott, H.R., Carpenter, J.W., Zhong, S., Ghosh, S., Bell, R.M., and Campbell, S.L. (1996). The solution structure of the Raf-1 cysteine-rich domain: a novel Ras and phospholipid binding site. Proc. Natl. Acad. Sci. U.S.A. 93(16), 8312–8317.

81. Saraboji, K., Håkansson, M., Genheden, S., Diehl, C., Qvist, J., Weininger, U., Nilsson, U.J., Leffler, H., Ryde, U., Akke, M., and Logan, D.T. (2012). The carbohydrate-binding site in galectin-3 is preorganized to recognize a sugarlike framework of oxygens: ultra-high-resolution structures and water dynamics. Biochemistry 51(1), 296–306.

