## Supplemental Information for "A structural model of a Ras-Raf signalosome"

### Supplemental Information
### A structural model of a Ras-Raf signalosome

#### Materials and Methods

##### *Molecular dynamics simulations*

*Simulation protocol and force field parameters*

Molecular dynamics (MD) simulations were performed on the special-purpose supercomputer Anton 2.[1] The simulated systems ranged in size from ~37,000 to ~1,740,000 atoms, and were parameterized using the TIP3P model[2] for water molecules, the Amber99SB*-ILDN force field[3–8] for proteins, and the CHARMM36 force field[9] for lipids. Specialized force field parameters were used for farnesylated cysteine,[10] phosphorylated serine and tyrosine,[11] and GDP and GTP.[12] Simulations of the disordered C-Raf linker by itself were exceptional in that the simulation systems were parameterized using a99SB-*disp*[13] and the TIP4P water model.[14] a99SB-*disp* is a variant of the Amber force field developed to work together with TIP4P for accurate simulations of disordered (as well as ordered) protein states. Due to concerns about force field inaccuracy for $Zn^{2+}$ ions, distance restraints were applied between CRD zinc finger residues and $Zn^{2+}$ ions in simulations that included the zinc-coordinated Raf CRD domain. MD simulations were performed in the NPT ensemble with constant pressure (1 bar) and constant temperature (310 K) imposed by a Martyna-Tuckerman-Klein Nosé-Hoover chain coupling scheme,[15] which was implemented using a multigrator scheme[16] with a relaxation time of 10 ps.

Initial velocities were sampled from the Boltzmann distribution. Water molecules and all bond lengths to hydrogen atoms were constrained using an in-house implementation[17] of M-SHAKE.[18] Production simulations (for data collection) were launched after energy minimization and nanosecond-timescale MD simulation with harmonic position restraints on backbone atoms.

The protein backbone atoms were restrained to their initial positions using an initial harmonic potential with a force constant of 10 kcal $mol^{-1}$ $Å^{-2}$ for 10–500 ns as an equilibration step; the force constant was linearly reduced to zero over the course of the equilibration step. Van der Waals and short-range electrostatic interactions were cut off at 10 Å for simulations without membrane and 12 Å for simulations with membrane. Long-range electrostatic forces were calculated in k-space using a grid-based method with Gaussian spreading[19] to the grid every 7.5 fs. The simulation time step was 1 fs for the equilibration stage and 2.5 fs for production simulations; the r-RESPA integration method[20] was used, with long-range electrostatics evaluated every 7.5 fs.

*System preparation*

Structures derived from the Protein Data Bank were back-mutated into the wild type unless stated otherwise, and missing atoms and residues were built in. The system of interest was placed at the center of a simulation box that was cubic for solvent simulations and orthorhombic for membranous simulations, with a separation greater than 20 Å from any periodic image. Explicitly represented water molecules were added to fill the system, and $Na^+$ and $Cl^-$ ions were included to maintain physiological salinity (150 mM) and to obtain a neutral total charge for the system. Residue protonation states corresponded to pH 7.

In membranous simulations, a phosphotidylcholine (POPC) lipid bilayer with ~30% phosphotidylserine (POPS) in the inner layer proximal to the proteins was used. The model membrane was built from a neutral POPC lipid membrane by replacing 15% (molar) POPC with negatively charged POPS lipids. (The POPC lipids to replace were all taken from the inner bilayer, but otherwise chosen at random.) This POPS fraction was chosen to mimic the abundance of anionic lipids in the mammalian plasma membrane,[21,22] in which approximately 10% of all lipids are POPS species, with other anionic species, such as phosphoinositides, bringing the total content of anionic lipids up to 15%. The POPS lipids were introduced only in the intracellular leaflet, which is where anionic lipids are almost exclusively found in cell membranes. The lipid content was thus ~30% POPS and ~70% POPC in the intracellular leaflet and 100% POPC in the extracellular leaflet.

*Simulations of K-Ras dimerization*

We performed 20 simulations, each 10 μs long, of two GTP-bound K-Ras molecules (PDB 4DSN) in aqueous solvent. In addition, we performed 23 unbiased simulations, each at least 2 μs long, of two GTP-bound K-Ras proteins anchored to the membrane by their farnesylated Cys185 (fCys185) residues.

*FRET assays*

Full details of our fluorescence resonance energy transfer (FRET) assay are available elsewhere.[23] Briefly, HEK293T cells were co-transfected with paired CFP- and YFP-fused KRAS constructs under the conditions of 0.5% and 10% fetal bovine serum (FBS) in 4-well chambered cover glass (Lab-Tek™). After 36–48 hours, live cell imaging was performed using a Confocal/Multiphoton Zeiss LSM880 microscope. To examine the GTP-dependent Ras-Raf interaction, co-transfection of CFP-K-RasWT and YFP-C-Raf, CFP-K-RasT35A and YFP-C-Raf, CFP-K-RasG60A and YFP-C-Raf, or CFP-K-RasG13D and YFP-C-Raf plasmids was conducted. After 24 hours of transfection, cells underwent 22 hours of serum starvation, followed by epidermal growth factor (EGF) (10 ng $mL^{-1}$) exposure for 30 minutes, and then were subjected to microscopy. Data were collected from three biological repeats and 10–12 different cells in different fields from the same coverslip were selected for microscopy. Quantitation was done using ZEN software (ZEISS).

*Preparation of GMP-PNP loaded K-Ras protein for EM studies*

K-Ras (1-188) wild-type protein was expressed and purified as described previously.[24] Point mutations were generated using the GeneArt™ Site-Directed Mutagenesis System (Life Technologies). K-Ras proteins (100 μM) in buffer (20 mM Hepes pH 8.0, 150 mM NaCl, 15 mM EDTA and 200 mM $(NH_4)_2SO_4$) were loaded with 200 μM GMP-PNP at 4°C overnight. The reaction was terminated by buffer exchange to 20 mM Hepes pH 8.0, 150 mM NaCl, 200 μM GMP-PNP, 10 mM $MgCl_2$ with Zeba Spin Desalting Column (Thermo Scientific™). GMP-PNP loading was verified by back extraction of nucleotide using 6 M urea and evaluation of nucleotide peaks by HPLC using an ion-exchange column as described previously.[24]

*Preparation of lipid monolayers*

Phospholipids obtained from Avanti Polar Lipids were mixed according to the following proportions in chloroform/methanol (3:1, vol/vol): 10% DOPS, 10% DOPE, 60% Egg-PE, 20% PE-MCC.[25] Assembly of protein on lipid monolayers was patterned after prior methods.[26] Briefly, droplets (15 μL) of K-Ras (10 μM) in buffer were added to Teflon depression wells (3 mm in diameter and 0.5 mm in depth). 0.5 μL of 1 mM lipid solution was then delivered to the surface of the droplet. The wells were incubated in an airtight, humidified chamber at 4°C overnight. The lipid monolayers at the air/liquid interface were collected by hydrophobic carbon-coated grids (Cat. CF400-Cu, Electron Microscopy Sciences). After washing with buffer 3 times, the grids were blotted and stained with uranyl formate solution (1% wt/vol) 3 times.

*Transmission electron microscopy (TEM) data collection and processing*

Stained grids were examined in a FEI Tecnai G2 Spirit Biotwin Transmission Electron Microscope operated at 120 kV with a magnification of 13,000 times. Images of were collected with the pixel size of 0.78 nm. RELION 3.0 was used for all image processing.[27] For WT K-Ras reconstructions, 73,282 particles were computationally selected from 185 micrographs and subjected to 2D classification into 100 classes.

*Raf localization imaging*

Raf localization was evaluated by co-transfection of plasmids expressing CFP-K-Ras$^{K88D}$ and YFP-C-Raf, CFP-K-Ras$^{Q129L}$ and YFP-C-Raf, and CFP-K-Ras$^{R149D}$ and YFP-C-Raf. Constructs were designed as previously reported.[23] Cells were cultured in 10% FBS. 36 hours post-transfection, cells were subjected to microscopy. Images were selected from three biological replicates and 10–20 different cells in different fields from the same coverslip were used for analysis. Imaging was done using ZEN software (ZEISS).

*BRET assays*

**Construct Design:** N-terminal fusion of KRAS of either mNeonGreen or NanoLuc were constructed in a pcDNA3.1 (+) vector (Invitrogen) as reporters for Bioluminescence Resonance

Energy Transfer (BRET) assay. KRAS point mutations were introduced either by site directed mutagenesis (QuikChange, Agilent) or replacing the KRAS gene with a synthetic DNA string (Geneart) carrying the desired mutation. All sequences were confirmed by sequencing.

**Transfection:** HEK293T cells were cultured in DMEM complete medium (10% FBS) and seeded in white 96-well clear-bottom plates (Corning) 24 h pre-transfection. Co-transfections of the reporter plasmids were carried out with TransIT-293 (Mirus) using the manufacturer's suggested protocol, using ratios as indicated in the figures. 24 h post-transfection, medium was exchanged to DMEM no phenol red (Life Technologies), containing varying amounts of FCS (0.1, 4.0, 10.0%).

**BRET measurements and data analysis:** 48 hours post-transfection, BRET measurements were taken at a Victor 3 Multilabel Plate Reader after the addition of 10 µL 32 µM coelenterazine 400a (Cayman Chemical) resulting in 2.9 µM final concentration. Emission of mNeonGreen and NanoLuc was observed for 2s at 535 ± 25 nm and 460 ± 25 nm, respectively. Expression of both reporters was monitored by measuring mNeonGreen (Ex 485 nm, Em 535 nm) before the addition of luciferase substrate, and measuring total luminescence directly after the BRET measurements for 0.3 s.

BRET ratios were calculated based on the following formula for each transfected well:

**BRET Ratio** = $Em_{535\ nm} / Em_{420\ nm} - Cf$

$Cf = Em_{535\ nm} / Em_{420\ nm}$ **(Donor only samples)**

The average of three technical replicates was calculated for each of the biological duplicates or triplicates after removal of obvious outliers and subjected to statistical analysis. Statistical analyses were performed using a one-way ANOVA followed by Dunnett's post-tests (*P < 0.05, **P < 0.01, ***P < 0.001, ****P < 0.0001).

*Plasmids used in the FRET experiments*

pcDNA3-CFP (Cat#13030), pcDNA3-YFP(Cat#13033), and pBABEpuro-CRAF (Cat#51124) plasmids were purchased from Addgene. Full-length fragments of KRASWT and CRAF were inserted into vectors containing CFP or YFP to obtain CFP-KRASWT, YFP-KRASWT, and YFP-CRAF constructs. KRASG12C, KRASG12D, KRASG13D, KRASD30R, KRASE31R, KRAST35A, KRAST35S, KRASG60A, KRASE62R, KRASR135A, KRASK147D, KRASD154Q, KRASG12C/D30R, KRASG12D/D30R, KRASG12C/E31R, KRASG12D/E31R, KRASG12C/E62R, KRASG12D/E62R, KRASG12C/R135A, KRASG12D/R135A, KRASR135A/D154Q, KRASG12C/K147D, KRASG12D/K147D, and KRASK147D/D154Q mutants were generated by site-directed mutagenesis using PfuUltra II Hotstart PCR Master Mix (Cat#600850-51). The sequences were confirmed by sequencing.

*Generation of K-Ras$^{lox}$/K-RAS$^{MUT}$ cells*

Full details of our generation of *K-Ras$^{lox}$*/*K-RAS$^{MUT}$* cells are available elsewhere.[23] Briefly, K-RAS$^{D154Q}$, K-RAS$^{D30R}$, K-RAS$^{E31R}$, K-RAS$^{E62R}$, K-RAS$^{K147D}$, and K-RAS$^{A135R}$ mutations, in cis with either a G12C or G12D mutation, were created by point mutagenesis from pBABE HA-tagged K-RAS$^{WT}$ retroviral plasmid (provided by Channing Der, Addgene plasmid # 75282). Retroviruses were generated by co-transfection of pBABE plasmids together with pAmpho plasmid into HEK293T cells using FuGENE® HD Transfection Reagent (Promega). The retroviruses were transduced into *H-Ras*$^{-/-}$*; N-Ras*$^{-/-}$*; K-Ras*$^{lox/lox}$ mouse embryonic fibroblasts (MEFs) followed by 2 weeks of puromycin selection (1 μg mL$^{-1}$) in Dulbecco's modified eagle medium (DMEM) supplemented with 10% FBS, 100 μg mL$^{-1}$ penicillin, and 100 units mL$^{-1}$ streptomycin. To obtain *K-Ras*$^{lox}$/*K-RAS*$^{MUT}$ clones, we then cultured cells in the presence of 4-hydroxytamoxifen (4OHT) (Sigma, 600 nM) for another two weeks in order to achieve complete deletion of endogenous *K-Ras* alleles.

*Growth assessment by IncuCyte*

Cells (1 × 10$^3$) were seeded in 96-well plates in 150 μL DMEM complete medium. The following day, 10% FBS medium was replaced by cell starvation medium (1%, 0.5%, or 0.1% FBS) after two washes with phosphate-buffered saline (PBS). Plates were incubated in the IncuCyte Zoom for real-time imaging, with three fields imaged per well under 10x magnification every two hours. Data were analyzed using the IncuCyte Confluence version 1.5 software, which quantified cell surface area coverage as confluence values. IncuCyte experiments were performed in triplicate. A single representative growth curve is shown for each condition.

*Western blot analysis*

Cells were lysed in RIPA lysis buffer (#89900 Thermo Fisher) supplemented with protease and phosphatase inhibitor cocktail tablets (Roche). The antibodies used for western blotting included those against: HA-Tag (6E2) (Cell Signaling Cat#2367), HSP90 (H114) (Santa Cruz Biotech Cat#sc-7947), phosphorylated Akt (Ser473) (Cell Signaling Cat#4060), Akt (Cell Signaling Cat#9272), phosphorylated ERK1/2 (Cell Signaling Cat#4370), ERK1/2 (Cell Signaling

Cat#4695), phosphorylated S6 (Ser235/236) (Cell Signaling Cat#4858), S6 ribosomal protein (Cell Signaling Cat#2217), anti-rabbit IgG, HRP-linked secondary antibody (Cell Signaling Cat#7074P2), ECL Sheep anti-Mouse IgG, HRP-linked secondary antibody (GE Healthcare Cat#NA931V), ECL Donkey anti-Rabbit IgG, and HRP-linked secondary antibody (GE Healthcare Cat#NA934V).

*Nucleotide exchange assay*

Our SOS1-mediated Ras nucleotide exchange assay was performed at Icagen (Arizona). The purified K-Ras mutant R135A was diluted in assay buffer (40 mM HEPES at pH 7.5, 1.5 μM Mant-GDP, 10 mM $MgCl_2$, 0.05% CHAPS, and 0.01% NP40) to a final concentration of 1 μM. The exchange reaction was triggered by adding purified human SOS1 (residues 564–1049, 0.05 μM) to the reaction mix. Kinetic readings were taken to measure fluorescence (excitation at 360 nm, emission at 450 nm) for 30 minutes at 30-second intervals using a 384-well plate reader.

*Bioinformatics analysis*

The K-Ras sequences used for evolution analysis were compiled using protein-protein BLAST searches in the NCBI non-redundant protein database, with the human K-Ras sequence (residues 1–166) as reference. The initial search generated 20,000 sequences with a minimal sequence identity of 29.84%. K-Ras sequences of mammals, birds, and fish were then extracted from the initial search results based on taxonomy IDs. The sequence pool was then filtered based on the following rules: 1) duplicate entries or near identical (sequence identity greater than 95%) entries were removed; 2) sequences with large (greater than 25 amino acids) insertions or deletions compared to human K-Ras were removed; 3) sequences labeled as "partial," "synthetic," or "predicted" were removed. After the filtering, 334 sequences remained and were used for multiple-sequence alignment. Sequence alignment of these sequences was obtained from the NCBI server using default settings, and sequence logos were generated using the online tool WebLogo3. The final figures were manually adjusted from the WebLogo3 result to show only the residues corresponding to human K-Ras (Residue 1-166).

#### Supplemental details of model construction

##### *Preparation of simulations by type*

###### *Simulations of K-Ras monomer in solvent, on the membrane, and in a lattice*

Wild-type K-Ras4B (residues 1–169) was prepared from the crystal structure PDB 4DSN with either GDP or GTP bound. Multiple simulations were launched, and the simulations in which GDP or GTP appeared unstable were discarded. For the membrane simulations, wild-type K-Ras4B (residues 1–169) was prepared from PDB 4DSN, loaded with either GTP or GDP, with the hypervariable region (HVR) residues 170–185 added to the structure in an initially extended conformation in which Cys185 was farnesylated (fCys185). The N terminus was charged. K-Ras was initially positioned in proximity to the membrane. This procedure was adopted for both the GDP- and GTP-bound K-Ras structures. For the crystal lattice simulation, GTP-bound K-Ras4B (residues 1–167) was arranged in an orthorhombic lattice, with 24 copies in the simulation cell, which was constructed based on PDB 3GFT.

*Simulations of K-Ras dimerization in solvent and on the membrane*

In solvent simulations, two copies of GTP-bound K-Ras proteins were placed in the simulation box without contact with one another and in arbitrary relative orientations, which differed in different simulations. In simulations of dimerization on the membrane, two copies of GTP-bound K-Ras were positioned on the membrane and not in contact with one another, with their HVR farnesyl groups buried in the membrane; the two K-Ras proteins were in the same orientation on the membrane initially. The initial membrane orientations were generated from simulations of monomeric GTP-bound K-Ras on the membrane (snapshots at 1 μs).

The GTP-mediated asymmetric (GMA) K-Ras dimer model was simulated in solvent. At the beginning of each simulation, the switch I regions in both the GTP donor and acceptor were adjusted to be consistent with the crystal structure PDB 4DSN.

*Simulations of an RBD-bound K-Ras monomer and dimer*

A model of monomeric GTP-bound K-Ras, bound in turn with the Ras-binding domain (RBD) of C-Raf, was constructed based on a crystal structure (PDB 4G0N) of H-Ras bound with the C-Raf RBD. This model was then simulated in solvent and on the membrane.

Based on the crystal structure PDB 4G0N, a C-Raf RBD was positioned on each protomer of the GMA K-Ras dimer and then simulated. At the beginning of each simulation, the switch I region of each K-Ras protein was adjusted to adopt the active conformation exhibited by the crystal structure PDB 4DSN. We simulated the RBD-bound K-Ras dimer both in solvent and on the membrane. In the setup of the simulations of RBD-bound K-Ras dimer on the membrane, the

K-Ras dimer was positioned in the membrane orientation from the simulation of K-Ras dimerization on the membrane (Figure 1A).

*Unbiased binding simulations of Gal-3 and farnesyl*

The Gal-3 carbohydrate-binding domain (residues 113–250) was prepared from the crystal structure PDB 3ZSM. Three copies of farnesylated cysteine, each capped on both ends (with N-terminal acetyl and C-terminal N-methyl amide residues), were initially placed in solvent at random positions inside the simulation box; the Gal-3 protein was positioned at the center. This simulation protocol is similar to one described previously.[28]

*Simulations of a C-Raf RBD and CRD with K-Ras in solvent and on the membrane*

Starting from the RBD-bound K-Ras structure, the nuclear magnetic resonance (NMR) structure of the cysteine-rich domain (CRD) of C-Raf (residues 136–187) (PDB 1FAR) was linked to the structure of the C-Raf RBD by a short linker (residues 132–135), and positioned in arbitrary orientations proximal to the K-Ras protein.

*Simulations of monomeric and dimeric C-Raf linker in solvent*

The C-Raf linker (residues 188–339) was constructed in an extended conformation and allowed to collapse in simulations. In simulations that included two copies of the C-Raf linker, a harmonic flat-bottom distance restraint was employed to restrict the distance between the Ser339 residues of the two strands to ~15 Å. This restraint was devised so that the linker dimer would be geometrically compatible with the C-Raf kinase domain (KD) dimer (residues 340–615), in which the distance between the two Tyr340 residues is 11.5 Å according to the crystal structure PDB 3OMV.

##### ***Construction of the Ras-Raf signalosome model***

###### *Building the K-Ras octamer*

The K-Ras octamer was constructed by repeating the GMA dimer interactions in a string of eight K-Ras proteins, starting from the K-Ras dimer on the membrane that was formed in the simulations of K-Ras dimerization, with the donor of the GMA dimer being the head of the octamer (Figure 1B). The switch I and II regions in each K-Ras protein were rebuilt to be consistent with the conformation in PDB 4DSN and to maintain the key interactions between GTP and Tyr32, Lys16, and Thr35. The interaction between the GTP γ-phosphate and either Arg135 or Lys128 that was observed in the GMA dimer model was conserved in constructing the octamer model. The switch II conformation was adjusted to create a groove in that region in the base-tier K-Ras proteins; this groove is important in the stacking interactions (to accommodate the β2–β3 hairpin of a K-Ras protein from the second tier). The series of polar interactions at the stacking interface (Lys165–Asp98, Lys172–Glu105, Arg161–Asp91, and Glu154–Arg88) were also maintained by minor adjustments to the side-chain conformations.

###### *Adding a C-Raf RBD to each K-Ras protein*

A C-Raf RBD (residues 54–131) was placed on each K-Ras unit, based on a structure of H-Ras in complex with the RBD (PDB 4G0N).

*Adding a Galectin-3 protein to each K-Ras protein*

The farnesylated cysteine (fCys185) of K-Ras was inserted into Gal-3 based on the complex structure of farnesylated cysteine-bound Gal-3, which was generated by unbiased binding simulations. The K-Ras HVR (residues 170–184) was built in with an extended conformation and allowed to relax in simulations. A Gal-3 protein at index *n* was positioned near K-Ras *n* such that it bound to the fCys185 and HVR of K-Ras, and its Lys210, Glu205, and Gln220 residues interacted with the Asp117, Arg100, and Asn74 residues of C-Raf RBD *n−1*, which itself made a primary interaction with K-Ras *n−1* (the GTP donor to K-Ras *n*). The Gal-3 protein bound to the fCys185 residue of K-Ras *1* was positioned in a similar orientation, although it did not interact with an RBD in the same manner. The four Gal-3 proteins that bound to the fCys185 residues of K-Ras *1–4* at the base tier were positioned on the membrane such that the first (residues 113–126) and last (residues 242–250) β strand of each Gal-3, and the K-Ras HVRs bound to each Gal-3 protein, were proximal to the membrane. The other Gal-3 proteins were positioned in an orientation similar to the Gal-3 proteins of the base tier: Gal-3 *n+4* stacked onto Gal-3 *n*, with β strands of the former (residues 242–250) and latter (residues 183–190) resembling an antiparallel β sheet. We considered only the structurally resolved carbohydrate-binding domain of Gal-3, not the unresolved N-terminal region.

*Adding a C-Raf CRD to each K-Ras protein*

We performed 24 simulations (totaling 122 μs) of the three-domain system in solvent (CRD tethered to a K-Ras–RBD complex), in which the CRD was initially in contact with neither the K-Ras protein nor the RBD. The simulations generated a set of structural models (Figure S7E). A CRD pose was considered only if, once positioned on the K-Ras helical assembly, it did not clash with any K-Ras proteins or the membrane. Based on the successful poses, we constructed models of membrane-bound K-Ras bound to both the RBD and CRD for further simulation (31 simulations of 140 μs in aggregate). In one of these simulations, the K-Ras–CRD structure adopted a stable pose in which the Glu174, Arg143, Phe151, Lys157, and Leu160 residues of the CRD were positioned proximally to the Arg41, Asp54, Met170, Glu76, and Ile163 residues, respectively, of K-Ras, and a CRD $Zn^{2+}$ interacts with Glu3 of K-Ras. The K-Ras/CRD interaction with this CRD pose exhibits electrostatic complementarity (Figure S7H).

In our signalosome model we used this K-Ras–CRD structure: The CRDs bound to the K-Ras proteins of the base tier also interact with the membrane, as do the $Zn^{2+}$ ions bound to the CRDs. The stability of the Glu76-Lys157 and Glu3-$Zn^{2+}$ interactions in CRD/K-Ras pairs, over the course of a nearly 100-μs long and restraint-free simulation of the complete signalosome model (Figure S7F), is indicative of the stability of the CRD/K-Ras pose in the model. In this model, the CRDs of the second tier interact with K-Ras molecules with the same pose, although they do not contact the membrane.

*Adding the C-Raf linkers and kinase domains to the model*

Our simulations of the C-Raf linker in solvent suggested that it may adopt a diverse set of conformations, with a large variation in the distance between the N and C termini. We selected eight different C-Raf linker conformations from these simulations. In the signalosome model, each linker was attached to a CRD (ending at Trp187), and the linker of C-Raf *n* (bound to K-Ras *n*) was positioned adjacent to the linker of C-Raf *n+4* (bound to K-Ras *n+4*). (Other C-Raf pairing patterns may also be possible, as discussed in the following section.) In the model, the KD of C-Raf *n* was arranged to dimerize with the KD of C-Raf *n+4*. The structure of the KD dimer (PDB 3OMV) was placed slightly above the membrane, with the Tyr340 residues facing inward toward the K-Ras assembly. An ATP and one $Mg^{2+}$ ion were placed at the ATP binding site of each KD. The C termini of the linkers were adjusted so that linker *n* and *n+4* were connected to a KD dimer. Steric clashes and interweaving of the linkers were avoided in the modeling. Phosphorylation of Ser338 and Tyr341 was also introduced. In subsequent simulations of the model, the two linkers developed extensive contact with one another (Figure 1F).

*Constructing an 8-protomer Ras-Raf signalosome model with alternative C-Raf pairing*

Although there are no clear experimental observations favoring a particular pattern of C-Raf pairing, given the length and flexibility of the C-Raf linker, it is structurally feasible for C-Raf *n* to pair with C-Raf *n*+1, C-Raf *n*+3, or C-Raf *n*+5, rather than with C-Raf *n*+4. C-Raf *n* pairing with C-Raf *n*+2 or CRD *n*+6 can be excluded, because the CRD C-termini separation for such a pair is too large (~150 Å). The separation between the CRD C termini is 80–100 Å for an *n/n+1*, *n/n+3*, or an *n/n+5* C-Raf pairing, while the separation is only 35–50 Å for an *n*/*n*+4 pairing. The signalosome model does not dictate a uniform pattern of Raf dimerization, but the *n/n+4* pairing (Figure 1F) would appear more likely if we consider that the other pairings require a significant part of a C-Raf linker to adopt an extended loop conformation to circumvent the Gal-3 proteins (Figure S9F, lower right panel). Further investigation is required to clarify whether C-Raf dimerization follows a uniform pattern and what that pattern might be. To demonstrate that other C-Raf dimerization patterns may be possible, we developed an alternative Ras-Raf signalosome model in which C-Raf *n* is paired with C-Raf *n*+1 (Figure S9F).

*Adding the 14-3-3σ dimer to each C-Raf KD dimer*

C-Raf was extended C-terminally to residue 625, with Ser621 phosphorylated. A 14-3-3σ dimer was attached to each KD dimer based on the crystal structure PDB 4IEA, in which each 14-3-3σ is bound with a C-Raf phospho-peptide (residues 618–625) including pSer621. The orientation of the 14-3-3σ dimer with respect to the KD dimer is unknown, but limited flexibility is afforded by the two residues (Ile616 and Asn617) that connect the KD and the phospho-peptide bound to 14-3-3σ. Under this constraint, together with an assumption of symmetry that ensures the two 14-3-3σ proteins interact with the two KDs in the same way, we constructed a model of the C-Raf KD dimer bound with the 14-3-3σ protein. Based on this hetero-tetrameric C-Raf KD–14-3-3σ model, 14-3-3σ proteins were incorporated into the signalosome model.

*Adding MEK1 to each C-Raf KD*

Based on a structure of a B-Raf KD bound with MEK1 (PDB 4MNE), we positioned a MEK1 KD (residues 66–382) abutting each C-Raf KD as a substrate protein in the signalosome model. This also positioned the N-lobe of MEK1 against 14-3-3σ. The missing residues of MEK1 (residues 275–305) were modeled as an extended loop, which collapsed towards the C-lobe of the MEK1 KD in simulations.

*Simulating the eight-protomer Ras-Raf signalosome model*

The simulated signalosome model consists of eight protomers, each of which includes a K-Ras, a Gal-3, a C-Raf, a 14-3-3σ, and a MEK1 protein. These components were incorporated into the model in a stepwise fashion: 1) by extending the GMA dimer, a K-Ras octamer was constructed and simulated. A model with consistent K-Ras/K-Ras interactions, and stability in a simulation of ~10 μs, was used in further modeling; 2) 8 C-Raf RBD domains were added to the K-Ras octamer based on an RBD-Ras crystal structure, and the resulting model was simulated. The K-Ras/RBD model was stable in these μs-long simulations; 3) based on our simulation studies of Gal-3 interactions with the farnesylated K-Ras tail, with full-length K-Ras protein, and with another Gal-3, 8 Gal-3 proteins were added to the K-Ras/RBD octamer model, and the resulting model was simulated. At the end, a K-Ras/RBD/Gal-3 model, stable for 20 μs in a simulation, was chosen for further modeling; 4) after our simulation study of CRD interaction with an RBD-bound K-Ras, 8 C-Raf CRD domains were added to the model and simulated. In the event that a Gal-3 molecule or a CRD deviated from the initial modeled position in the simulation, we manually repositioned the Gal-3 or CRD, and then relaunched the simulation. A model with consistent CRD/K-Ras and Gal-3/K-Ras interactions, stable in μs-long simulations, was chosen; 5) after simulations of C-Raf linkers and their interaction with one another in isolation, and modeling of the two-protomer C-Raf/MEK/14-3-3σ complex, each C-Raf in the octamer model was extended to full length, with the linkers paired with one another, and with the kinase domains dimerized and interacting with 14-3-3σ dimers and MEK kinases. This model was then simulated to nearly 100 μs, and remained stable in the simulation without any artificial restraints (Figure S1B).

In step 5, the simulation of a complete signalosome model consisted of five phases. In the first phase, weak harmonic positional restraints (with a spring constant of 1 kcal $mol^{-1}$ $Å^{-2}$) were applied to the backbone atoms of the CRD and the KD domain of each C-Raf for 1 μs. This allowed the disordered C-Raf linkers to equilibrate. In the second phase, weak harmonic flat-bottom distance restraints were employed for ~1 μs to facilitate structural relaxation at the predicted protein-protein interfaces, particularly the CRD interface with K-Ras and the Gal-3/Gal-3 interface. This was designed to allow these protein-protein interactions to equilibrate after more stringent harmonic positional restraints were removed. In the third phase, all restraints were removed, and extensive equilibrium MD simulations were performed. In the fourth phase, the complex was inspected for overall stability and symmetry in the interaction surfaces. Any Gal-3 protein, C-Raf CRD, or MEK1 kinase that had deviated substantially from the initial model was remodeled, and then the simulation was relaunched. These four phases were repeated, until a stable signalosome model was achieved. In the fifth phase, the final signalosome model was simulated for nearly 100 μs without any restraint applied to any protein-protein interactions.

#### Supplemental analyses

##### *Simulations of GTP-bound K-Ras in a crystal packing*

In X-ray crystal structures of wild-type Ras, although binding to GTP analogs appears to be more stabilizing to the active conformation of the switch regions than GDP binding, conformational heterogeneity remains (Figure S3E and S3F). In crystal structures of the GTP-bound Ras mutant T35S, the heterogeneity is more pronounced (Figure S3G), consistent with the fact that T35S is an inactivating mutation. The conformation of Ras bound to GTP analogs in crystals at the switch regions (Figure S3E) appears much less varied than in our simulations of GTP-bound K-Ras (Figure S3B). To reconcile this apparent discrepancy, we set up a crystal lattice with 24 copies of wild-type GTP-bound K-Ras in a simulation box, based on the crystal lattice underlying PDB 3GFT, and simulated this for 4 μs. From the 24 copies, we took a conformational snapshot every 1 μs. We found that K-Ras was significantly less flexible in the lattice than in solvent; in the lattice the relative population for the active conformation at the switch one region was 45%, as opposed to 11% in the solvent (Figure S3A and S3H). This result suggests that crystal packing may mask the inherent conformational fluctuation of GTP-bound K-Ras. We also simulated RBD-bound K-Ras, and found that the switch regions were most stable in the active conformation upon RBD binding (Figure S3D).

***Timescale of protein dimerization and the length of our simulations***

In the absence of a favorable electrostatic steering force, an upper bound for the "on" rate of protein-protein binding is ~$10^6$ $M^{-1}s^{-1}$.[29] At the millimolar protein concentration relevant for our simulations, the average time required for a binding event is thus ~1 ms. Since the timescale of our individual simulations of K-Ras dimerization (~10 μs) is significantly shorter than that of protein-protein, we would not expect most of the simulations to converge to a single dimer model. Of the tens of K-Ras/K-Ras binding simulations we conducted, indeed only two (one in solvent and one with membrane) arrived at the GMA K-Ras dimer structure. (We did not obtain any other dimer from these simulations more than once, although this itself is not sufficient evidence to support the GMA dimer.) To obtain a Boltzmann ensemble of K-Ras dimer conformations, the simulations need to sample both the formation and the dissociation of the GMA dimer adequately. The former event has only been sampled twice, and the latter has not been sampled. We thus do not expect that the snapshot from all the simulations combined can generate the Boltzmann ensemble of K-Ras dimers, or a correct relative population of the dimers.

***Analogous Arg-GTP interaction in Toc33-Toc34 dimers***

A trans interaction similar to that in the GMA K-Ras dimer model has been reported[30] between arginine and GTP phosphate in GTP-mediated dimers of Toc33 and Toc34 (Figure S2F, PDB

3BB1), two isoforms of a GTPase receptor protein in the outer envelope of chloroplasts; indeed, these homodimers do not promote GTP hydrolysis.[31]

##### *$^{15}$N-HSQC broadening profile of K-Ras and K-Ras dimerization*

Twelve residues exhibited broadening of $^{15}$N-HSQC spectra upon sample dilution of GTP-γS-bound truncated K-Ras-4B (Residue 1–166).[32] Eight of the twelve residues (Lys16, Asp30, Glu31, Asn86, Asp119, Gln131, Arg135, Lys147) are located at the GMA dimer interface, and four are not (Ile24, Glu37, Ser39, Val44) (Figure 2E). The broadening of the eight residues can be easily explained by the dimerization. The broadening of Ile24 (immediately N-terminal to switch I) and Glu37 and Ser39 (immediately C-terminal to the switch I) is consistent with the notion that GMA dimerization stabilizes the donor's active conformation of switch I. Val44 is adjacent to the α5 helix of the acceptor interface, and the broadening may be a result of the altered α5 dynamics in the GMA dimer.

NMR chemical shift perturbations (CSPs) were also observed upon sample dilution of full-length GTP-γS-bound K-Ras4B for nine residues. The CSP profile of the full-length construct is not identical to that of the tail-truncated construct (residues 1–166), potentially because in solution the HVR tail interacts with the catalytic domain and affects K-Ras dimerization.[33] The CSPs are broadly consistent with the GMA dimer as well. Of the nine CSP residues, six residues are part of the GTP-acceptor interface (Ile84, Val125, Ala130, and Glu143) or immediately adjacent to the interface (Val114 and Tyr157); Glu91 is adjacent the GTP-donor interface; Asp38 is immediately C-terminal to the switch I; Ala18 is secluded and interacts intimately with the mediating GTP, and may be indirectly affected by the GTP-mediated dimerization.

***Intriguing questions with regard to G13D and G13R oncogenic mutations***

Unlike the other oncogenic mutations we tested, in our simulations G13R and G13D disrupted the GMA K-Ras dimer (Figure S2I). This is corroborated by our preliminary BRET data, which showed that G13D disrupts K-Ras assembly (Figure S5F). Further investigation is required to address whether these two mutations indeed are incompatible with the GMA dimer and if so, what might be the biological implications.

***Complicating factors associated with the R135A mutation in K-Ras***

In light of the interaction between Arg135 and GTP predicted by the GMA dimer model, we tested the R135A mutation, with the expectation that its effect would be similar to the other four mutations we tested. We did not, however, observe increased CFP emission after YFP bleaching for R135A (Figure S5E), and its effect on *K-Ras*$^{lox}$/*K-RAS*$^{MUT}$ cell growth was moderate (Figure 3F and Figure S4B). R135A also did not disrupt ERK phosphorylation, although it boosted AKT phosphorylation (Figure 3G and Figure S4C). To reconcile these results, we considered the possibility that R135A might alter the biochemical properties of K-Ras. Indeed, although Arg135 is distal from the nucleotide binding site, we found that in the absence of any GEF protein, the inherent nucleotide exchange rate of R135A was significantly higher than that of the wild-type (Figure S5D). This may have increased the population of GTP-bound K-Ras in the cells and masked the effect of R135A on K-Ras dimerization.

##### *R135A/R128A double mutation*

The double mutation R135A/R128A attenuates H-Ras nanoclustering.[34] In addition to reducing effector recruitment and MAPK signaling, the double mutation is thought to affect nanoclustering primarily by altering the orientation of Ras on the membrane.[35] If similar helical assemblies of H-Ras underlie the nanoclustering, our model suggests that the R135A/R128A mutation would attenuate H-Ras nanoclustering and signaling by disrupting GMA H-Ras dimers. Given the connection between the GMA dimerization and the orientation of Ras on the membrane (Figure 2G), the observed change in Ras orientation may in part be associated with a double mutation–induced change in Ras dimerization.

##### *The E62R mutation abolishes AKT phosphorylation*

The E62R mutation is unique among the mutations we tested in this study, in that it abolishes AKT phosphorylation (Figure 3G) and is located at the switch II region. Raf primarily binds at the switch I region, but PI3K kinase binds with both the switch I and II regions.[36] Although E62R may disrupt the GMA dimerization and hinder ERK phosphorylation, it may more effectively disrupt Ras-PI3K binding and affect AKT phosphorylation in the PI3K-AKT pathway. Further investigation is needed to understand the effect of the E62R mutation on this pathway. Basal expression levels of the E62R mutant were lower than those of the other mutants, in agreement with previous reports that demonstrated that some mutations in the switch II region of K-Ras are hypomorphic.[37,38]

***Simulations investigating membrane orientations of K-Ras***

In the NMR-identified α orientation, a bound GTP tends to have the guanine moiety closer than the phosphates to the membrane, but in the NMR-identified β orientation the opposite is the case, and the GTP is also further removed from the membrane. We thus used the guanine and phosphate distances to the membrane to represent a K-Ras orientation (Figures S2D and S2E). We simulated a GTP-bound K-Ras monomer on the membrane for 67 μs in total and a (GTP-bound) K-Ras monomer bound with a C-Raf RBD on the membrane for 50 μs in total, and compared the distributions of K-Ras orientations in these simulations with the distribution in a simulation of the GMA dimer. The RBD-free monomer adopted two clusters of orientations, one in the RBD-compatible α region, and one in the RBD-occluding β region (Figure 2G). In contrast, the RBD-bound monomer only adopted orientations of the α region, as did the two K-Ras proteins in the simulations of a GMA dimer. In the α orientation, as suggested by a previous study,[39] Met170 at the C terminus of the α5 helix tended to be partially buried in the membrane in our simulations. This analysis indicates that the membrane interaction of K-Ras may promote GMA dimerization by restraining the K-Ras orientation, and that GMA dimerization may promote K-Ras recruitment of Raf proteins by imposing membrane orientations favorable to RBD binding.

***Estimate of the local concentration of constituent proteins in the Ras-Raf signalosome***

Our modeled eight-protomer signalosome is approximately 140 Å in radius and 80 Å in height, and thus about $5 \times 10^6$ Å$^3$ in volume. This translates to a local concentration of approximately 2.7 mM for each of the eight proteins constituent to the signalosome.

***Additional details of the secondary and tertiary Ras-Ras interactions***

In the secondary Ras-Ras (stacking) interaction, the α3 helix and N-terminal part of the switch II region of a K-Ras protein at position *n* packs with the α5 helix and β2-β3 hairpin of K-Ras *n*+4 (Figure 4C). The stacking interface of K-Ras *n*+4 largely overlaps with the membrane interface of a K-Ras protein in the base tier (Figure 1B). The stacking interaction of two K-Ras proteins involves a buried interface with an area of about ~1450 $Å^2$ and a number of salt bridges at the interface, including Glu98-Lys165 and Asp105-Lys172 (Figure S6A). This may explain a recent report that E98K/D105K and K165E/K172D mutations individually disrupt K-Ras nanoclustering but together rescue it.[40] The stacking is also consistent with K-Ras nanoclustering being disrupted by K101A/R102A and H94A/H95A mutations,[41] as these residues are also located at the stacking interface in the helical assembly. The tertiary interface is smaller, with a buried interface area of ~660 $Å^2$, and involves the α4 helix of K-Ras *n* and the α1 helix and switch I region of K-Ras *n*+3 (Figures 4C, right panel, and Figure S6B).

***The locations of various reported Ras mutations in the signalosome model***

The surface of a K-Ras protein in the signalosome model is almost entirely covered by interactions with other proteins, and so it is not surprising that many mutations affecting the function of K-Ras are located at the interfaces between Ras and other proteins in the signalosome model. We did not consider these mutations in the modeling. Both the K-Ras helical assembly and interactions of K-Ras with the membrane were determined by the membrane-anchored GMA K-Ras dimer model generated by simulations, and the RBD interactions with the Ras assembly were determined by the resolved crystal structure of the Ras-RBD complex.

Many cancer mutations and RASopathy mutations, including T50I, C51Y, R164Q, G48R, E49K, D47A/E49A, E153V, and F156L, are located at the β2-β2 turn and the α4 and α5 helices (the so-called "switch III" region[35]) and alter Ras nanoclustering.[42] At the base tier of the signalosome model, these mutations are involved in the Ras-membrane interaction (Figure S8A). This is consistent with the possibility that the adverse effect of these mutations comes from altering the interactions of Ras with the membrane.[42,43] Additionally, in the signalosome model these mutations are located at the upper interface of the Ras-Ras stacking (i.e., the interface of Ras *n+4* stacked with Ras *n*), which primarily involves the β2-β2 turn and α5 helix (Figure 4C, right panel). The functional effects of these mutations and their connections with RASopathy and cancer thus may partially arise from their effects on Ras-Ras stacking in addition to Ras-membrane interactions. Two H-Ras mutations, G48R and D92N, are found together in spitzoid tumor samples and are known to elevate H-Ras nanoclustering.[42] Assuming the signalosome model is broadly applicable to H-Ras, we suggest that Gly48 and Asp92 may be adjacent to one another in *trans* in Ras stacking (Gly48 at the upper stacking interface of Ras *n+4* and Asp92 at the lower stacking interface of Ras *n* (Figure S6A)), and these two mutations may interact with one another, potentially strengthening the stacking and promoting the formation of the Ras signalosome.

K101E, E107K, K101A/R102A, H94A/H95A,[40] E98K/D105K, and K165E/K172D[41] are Ras mutations that were identified to test different Ras dimer and oligomer models from those we report here. These mutations were found to disrupt the Ras nanoclustering and, intriguingly, are located at the stacking interfaces (Figure S6A). Making either the K101E/E107K double mutation[40] or the E98K/D105K/K165E/K172D quadruple mutation[41] appears to recover the nanoclustering. This could be explained by the previously proposed models, but it could also be explained by the stacking interaction of Ras in the signalosome model: Since residues belonging to these two sets of mutations interact directly at the stacking interface, single-residue mutations that individually disrupt salt bridges at the interface may restore them when combined.

***Modeling of Gal-3 and Raf-CRD***

In the modeling, we attempted to accommodate the findings that the RBD mutations D117A and D117R weaken the nanoclustering and RBD-galectin binding;[44] that the Gal-3 mutation V126A does not appear to affect the farnesyl binding, but does affect Gal-3 colocalization with K-Ras at the membrane;[45] and that a thiodigalactoside-derived Gal-1 inhibitor does not affect the interaction between Gal-1 and the C-Raf RBD.[44] In the resulting signalosome model, the Asp117 residue of the RBD is located at the Gal-3–RBD interface, the thiodigalactoside inhibitor binding site of Gal-3 is positioned away from that interface, and Val126 is proximal to the membrane (Figure S7A). At the base tier, the Gal-3 position relative to K-Ras is broadly consistent with the position of similar farnesyl capping proteins (such as PDE6δ) relative to small G proteins (Figure S7C). The unstructured N-terminal segment of Gal-3 may be crucial to Gal-3 oligomerization;[46] although we did not include this in our model, ample space is available to accommodate it.

A number of Gal-3 mutations have been reported to disrupt Gal-3 dimerization (W181L),[47] Gal-3 co-localization with K-Ras (G182A),[46] and K-Ras nanoclustering (V126A).[45] Additionally, the Gal-1 quadruple mutation C3S/L5Q/V6D/A7S—corresponding to Gal-3 residues Pro113, Ile115, Val116, and Pro117—has been reported to disrupt Gal-1 dimerization and H-Ras nanoclustering.[44] We accommodated these findings in our modeling of Gal-3–Gal-3 interactions, and in the signalosome model these Gal-3 residues are located at the Gal-3 stacking interfaces (Figure S7B).

A CRD is connected with an RBD by a linker of only four residues, and limited exposed K-Ras surface area is available for interacts with a CRD in the helical assembly after RBD binding. The CRD pose is largely constrained by RBD-CRD connectivity, the interactions of the two zinc ions in contact with the membrane, and the limited available space. CRD mutations at Ser177, Thr182, Met183,[48] Phe151, Leu149,[49] and Ras mutation at Val45[50] have been shown to affect CRD-Ras binding. In the Ras-CRD pose adopted in the model, these mutations are located at or near the Ras-CRD interface (Figure S7F) or at the CRD interface with the membrane. Further, in the model the CRD directly interacts with the switch II region of Ras, which is consistent with the finding that mutations in the switch II regions (G60A and Y64W) may disrupt CRD binding.[51] Although the Ras-CRD pose in the model is consistent with existing experimental data, it remains to be experimentally validated; we note, however, that this pose is not an essential element of the overall structure of the Ras-Raf signalosome model.

***Modeling of the C-Raf kinase dimer in complex with a 14-3-3σ dimer***

A dimer structure of 14-3-3σ, in which each monomer is bound to a peptide representing the phosphorylated C-terminal loop (residues 618–625) of C-Raf, has previously been resolved.[52] We modeled a complex of the 14-3-3σ dimer and C-Raf kinase domain dimer (PDB 4IEA).[53] This was largely constrained by the fact that only two unresolved C-Raf residues connect the kinase domain and the resolved 14-3-3σ–bound C-terminal loop. Based on a crystal structure of MEK1-bound C-Raf kinase domain,[54] we added two MEK1 kinases to the complex structure as substrates of C-Raf (Figure 1F). (A Raf/MEK/14-3-3 complex structure was reported by Eck and colleagues[55] while this manuscript was under revision. Our model, at a high level, is largely consistent with this cryo-EM structure (Figure S9A), although in our model the Raf kinase domains make more direct contact with 14-3-3 proteins than in the cryo-EM structure.)

##### *The signalosome model may not be applicable to other Ras effectors*

It is far from clear whether the Ras-Raf signalosome model is applicable to other effectors beyond Raf kinases. This model is almost certainly not applicable to PI3K signaling: A PI3K protein would clash with the membrane at the base tier of the Ras helical assembly and lose access to its phosphatidylinositol 4,5-bisphosphate (PIP2) substrate lipid at any higher tiers of the helical assembly. Raf and PI3K are laterally segregated on the membrane in nanoclusters.[56] An increase in the concentration of Gal-1 has been shown to lead to an increase in C-Raf activation by H-Ras[57] but a decrease in PI3K activation.[58] In light of PI3K preferentially binding to Ras nanoclusters in which Ser181 is phosphorylated,[59] we conjecture that distinct complex structures underlie Ras-Raf and Ras-PI3K nanoclusters. Consistent with this notion, with the exception of E62R, the mutations we designed to disrupt GMA K-Ras dimerization do not reduce phosphorylation of AKT in the PI3K pathway (Figures 3G and S4C).

On the other hand, the reported signalosome model, if proven correct for K-Ras, may be broadly relevant to MAPK signaling: In many respects, findings on H-Ras nanoclustering echo those on K-Ras, and the signalosome model is consistent with the structures of other Ras and Raf proteins (such as N-Ras, H-Ras, A-Raf, and B-Raf) involved in MAPK signaling. With local alterations, the model may be extendable to those Ras and Raf proteins and provide a framework for understanding their overlapping yet distinct roles in MAPK signaling.

##### *Interactions involving the switch II pocket of K-Ras*

SOS1 carries four basic residues (Lys949, Arg950, His951, and Lys953) at the turn of the helical hairpin that are not conserved beyond the SOS family. The helical hairpin is inserted into the switch II pocket of K-Ras in the putative secondary SOS1 interaction with K-Ras in the helical assembly. This could potentially explain the specificity of SOS1 in up-regulating MAPK signaling.

##### *The effect of the D154Q/R161E double mutation*

Ambrogio et al. reported[23] that D154Q and R161E single mutations disrupt K-Ras dimerization. These findings were used to support a symmetric K-Ras dimer model that uses the α4-α5 helices as the dimer interface. Because the GTP acceptor in the GMA dimer model uses the same interface, and D154 and R161 are involved in salt bridges at the GMA dimer interface (Figure 3B), the findings are highly consistent the GMA dimer model. The previous study also showed that D154Q/R161E recovers K-Ras nanoclustering, and this was interpreted as supporting evidence for the symmetric K-Ras dimer model. Our simulation of an octameric D154Q/R161E helical assembly showed that these two mutations may be compensatory in the K-Ras helical assembly. The simulation showed that D154Q disrupts the D154-K147 salt bridges at the primary interfaces, but repositions the α5 helix, enabling the formation a compensatory E161-R102 salt bridge at the secondary interface (Figure S6G).

##### *SOS1 and GAP access to K-Ras molecules beyond the base tier*

Our structural analysis of SOS1 suggests that the linker connecting the GEF and the PH domains of SOS1 (residues 546–579) is sufficiently long and flexible to allow the SOS1-GEF domain to dock to any K-Ras of at least the second tier of the helical assembly while the SOS1-PH domain is anchored to the membrane in a favorable membrane orientation [60] (Figure S9G). This analysis suggests that the helical assembly cannot grow beyond the second tier, which is consistent with a previous estimate that a K-Ras nanocluster contains 5–8 K-Ras proteins.[61] It is conceivable, however, that the helix portion of the PH-GEF linker (PDB entry 1XD4) could unwind, allowing greater reach for the GEF domain and thereby larger K-Ras assemblies. Similar analysis also suggested that the linker connecting the GAP domain and the membrane-anchoring C2 domain of Ras-GAP protein (residues 677–747) is amply long to allow the GAP domain access to K-Ras proteins in the second (and possibly higher) tiers of the helical assembly.

#### Supplemental References


1. Shaw, D.E., Grossman, J.P., Bank, J.A., Batson, B., Butts, J.A., Chao, J.C., Deneroff, M.M., Dror, R.O., Even, A., Fenton, C.H., Forte, A., Gagliardo, J., Gill, G., Greskamp, B., Ho, C.R., Ierardi, D.J., Iserovich, L., Kuskin, J.S., Larson, R.H., Layman, T., Lee, L.-S., Lerer, A.K., Li, C., Killebrew, D., Mackenzie, K.M., Mok, S. Y.-H., Moraes, M.A., Mueller, R., Nociolo, L.J., Peticolas, J.L., Quan, T., Ramot, D., Salmon, J.K., Scarpazza, D.P., Schafer, U.B., Siddique, N., Snyder, C.W., Spengler, J., Tang, P.T.P., Theobald, M., Toma, H., Towles, B., Vitale, B., Wang, S.C., and Young, C. (2014). Anton 2: Raising the bar for performance and programmability in a special-purpose molecular dynamics supercomputer. In Proc. International Conference for High Performance Computing, Networking, Storage and Analysis (SC14), pp. 41–53 (IEEE).

2. Jorgensen, W.L., Chandrasekhar, J., Madura, J.D., Impey, R.W., and Klein, M.L. (1983). Comparison of simple potential functions for simulating liquid water. J. Chem. Phys. *79(2)*, 926–935.

3. Joung, I.S. and Cheatham III, T.E. (2008). Determination of alkali and halide monovalent ion parameters for use in explicitly solvated biomolecular simulations. J. Phys. Chem. B *112(30)*, 9020–9041.

4. Wang, J., Cieplak, P., and Kollman, P.A. (2000). How well does a restrained electrostatic potential (RESP) model perform in calculating conformational energies of organic and biological molecules? J. Comput. Chem. *21(12),* 1049–1074.

5. Duan, Y., Wu, C., Chowdhury, S., Lee, M.C., Xiong, G., Zhang, W., Yang, R., Cieplak, P., Luo, R., Lee, T., Caldwell, J., Wang, J., and Kollman, P.A. (2003). A point-charge force field for molecular mechanics simulations of proteins based on condensed-phase quantum mechanical calculations. J. Comput. Chem. *24(16),* 1999–2012.

6. Hornak, V., Abel, R., Okur, A., Strockbine, B., Roitberg, A., and Simmerling, C. (2006). Comparison of multiple Amber force fields and development of improved protein backbone parameters. Proteins *65(3)*, 712–725.

7. Best, R.B. and Hummer, G. (2009). Optimized molecular dynamics force fields applied to the helix-coil transition of polypeptides, J. Phys. Chem. B *113(26)*, 9004–9015.

8. Lindorff-Larsen, K., Piana, S., Palmo, K., Maragakis, P., Klepeis, J.L., Dror, R.O., and Shaw, D.E. (2010). Improved side-chain torsion potentials for the Amber ff99SB protein force field. Proteins *78(8)*, 1950–1958.

9. Klauda, J.B., Venable, R.M., Freites, J.A., O'Connor, J.W., Tobias, D.J., Mondragon-Ramirez, C., Vorobyov, I., MacKerell Jr., A.D., and Pastor, R.W. (2010). Update of the CHARMM all-atom additive force field for lipids: validation on six lipid types. J. Phys. Chem. B *114(23)*, 7830–7843.

10. Khoury, G.A., Thompson, J.P., Smadbeck, J., Kieslich, C.A., and Floudas, C.A. (2013). Forcefield_PTM: ab-initio charge and AMBER force field parameters for frequently occurring post-translational modifications. J. Chem. Theory Comput. *9(12)*, 5653–5674.

11. Steinbrecher, T., Latzer, J., and Case, D.A. (2012). Revised AMBER parameters for bioorganic phosphates. J. Chem. Theory Comput. *8(11)*, 4405–4412.

12. Meagher, K.L., Redman, L.T., and Carlson, H.A. (2003). Development of polyphosphate parameters for use with the AMBER force field. J. Comput. Chem. *24(9)*, 1016–1025.

13. Robustelli, P., Piana, S., and Shaw, D.E. (2018). Developing a molecular dynamics force field for both folded and disordered protein states. Proc. Natl. Acad. Sci. U.S.A. *115(21)*, E4578–E4766.

14. Abascal, J.L.F. and Vega, C. (2005). A general purpose model for the condensed phases of water: TIP4P/2005. J. Chem. Phys. *123(23)*, 234505.

15. Martyna, G.J., Tobias, D.J., and Klein, M.L. (1994). Constant pressure molecular dynamics algorithms. J. Chem. Phys. *101(5)*, 4177–4189.

16. Lippert, R.A., Predescu, C., Ierardi, D.J., Mackenzie, K.M., Eastwood, M.P., Dror, R.O., and Shaw, D.E. (2013). Accurate and efficient integration for molecular dynamics simulations at constant temperature and pressure. J. Chem. Phys. *139(16*), 164106.

17. Lippert, R.A., Bowers, K.J., Dror, R.O., Eastwood, M.P., Gregersen, B.A., Klepeis, J.L., Kolossvary, I., and Shaw, D.E. (2007). A common, avoidable source of error in molecular dynamics integrators. J. Chem. Phys. *126(4)*, 046101.

18. Kräutler, V., Van Gunsteren, W.F., and Hünenberger, P.H. (2001). A fast SHAKE algorithm to solve distance constraint equations for small molecules in molecular dynamics simulations. J. Comput. Chem. *22(5)*, 501–508.

19. Shan, Y., Klepeis, J.L., Eastwood, M.P., Dror, R.O., and Shaw, D.E. (2005). Gaussian split Ewald: a fast Ewald mesh method for molecular simulation. J. Chem. Phys. *122(5)*, 054101.

20. Tuckerman, M., Berne, B.J., and Martyna, G.J. (1992). Reversible multiple time scale molecular dynamics. J. Chem. Phys. *97(3)*, 1990–2001.

21. Zachowski, A. (1993). Phospholipids in animal eukaryotic membranes: transverse asymmetry and movement. Biochem. J. *294(1)*, 1–14.

22. Van Meer, G., Voelker, D.R., and Feigenson, G.W. (2008). Membrane lipids: where they are and how they behave. Nat. Rev. Mol. Cell Biol. *9(2)*, 112–124.

23. Ambrogio, C., Köhler, J., Zhou, Z.-W., Wang, H., Paranal, R., Li, J., Capelletti, M., Caffarra, C., Li, S., Lv, Q., Gondi, S., Hunter, J.C., Lu, J., Chiarle, R., Santamaría, D., Westover, K.D., and Jänne, P.A. (2018). KRAS dimerization impacts MEK inhibitor sensitivity and oncogenic activity of mutant KRAS. Cell *172(4)*, 857–868.

24. Lu J, Harrison RA, Li L, Zeng M, Gondi S, Scott D, Gray NS, Engen JR, Westover KD. (2017). KRAS G12C drug development: discrimination between switch II pocket configurations using hydrogen/deuterium-exchange mass spectrometry. Structure *25*, 1442–1448.

25. Gureasko J, Galush WJ, Boykevisch S, Sondermann H, Bar-Sagi D, Groves JT, Kuriyan J. Membrane-dependent signal integration by the Ras activator Son of sevenless. Nat Struct Mol Biol. 2008 May;15(5):452-61.

26. Shen QT, Bai XC, Chang LF, Wu Y, Wang HW, Sui SF. Bowl-shaped oligomeric structures on membranes as DegP's new functional forms in protein quality control. Proc Natl Acad Sci U S A. 2009 Mar 24;106(12):4858-63.

27. Scheres SH. RELION: implementation of a Bayesian approach to cryo-EM structure determination. J Struct Biol. 2012 180(3):519-30.

28. Shan, Y., Kim, E.T., Eastwood, M.P., Dror, R.O., Seeliger, M.A., and Shaw, D.E. (2011). How does a drug molecule find its target binding site? J. Am. Chem. Soc. *133(24)*, 9181−9183.

29. Schreiber, G., Haran, G., and Zhou, H.-X. (2009). Fundamental aspects of protein-protein association kinetics. Chem. Rev. *109(3)*, 839–860.

30. Gasper, R., Meyer, S., Gotthardt, K., Sirajuddin, M., and Wittinghofer, A. (2009). It takes two to tango: regulation of G proteins by dimerization. Nat. Rev. Mol. Cell Biol. *10(6)*, 423–429.

31. Koenig, P., Oreb, M., Rippe, K., Muhle-Goll, C., Sinning, I., Schleiff, E., and Tews, I. (2008). On the significance of Toc-GTPase homodimers. J. Biol. Chem. *283(34)*, 23104–23112.

32. Muratcioglu, S., Chavan, T.S., Freed, B.C., Jang, H., Khavrutskii, L., Freed, R.N., Dyba, M.A., Stefanisko, K., Tarasov, S.G., Gursoy, A., Keskin, O., Tarasova, N.I., Gaponenko, V., and Nussinov, R. (2015). GTP-dependent K-Ras dimerization. Structure 23(7), 1325–1335.

33. Chavan, T.S., Jang, H., Khavrutskii, L., Abraham, S.J., Banerjee, A., Freed, B.C., Johannessen, L., Tarasov, S.G., Gaponenko, V., Nussinov, R. and Tarasova, N.I. (2015). High-affinity interaction of the K-Ras4B hypervariable region with the Ras active site. Biophys J, *109(12)* 2602–2613.

34. Guzmán, C., Šolman, M., Ligabue, A., Blaževitš, O., Andrade, D.M., Reymond, L., Eggeling, C., and Abankwa, D. (2014). The efficacy of Raf-recruitment to H-ras depends on H-ras membrane conformer specific nanoclustering. J. Biol. Chem. *289(14)*, 9510−99533.

35. Abankwa, D., Hanzal-Bayer, M., Ariotti, N., Plowman, S.J., Gorfe, A.A., Parton, R.G., McCammon, J.A., and Hancock, J.F. (2008). A novel switch region regulates H-ras membrane orientation and signal output. EMBO J. *27(5)*, 727–735.

36. Pacold, M.E., Suire, S., Perisic, O., Lara-Gonzalez, S., Davis, C.T., Walker, E.H., Hawkins, P.T., Stephens, L., Eccleston, J.F., and Williams, R.L. (2000). Crystal structure and functional analysis of Ras binding to its effector phosphoinositide 3-kinase γ. Cell *103(6)*, 931–944.

37. Shieh, A., Ward, A.F., Donlan, K.L., Harding-Theobald, E.R., Xu, J., Mullinghan, C.G., Zhang, C., Chen, S.-C., Su, X., Downing, J.R., Bollag, G.E., and Shannon, K.M. (2013). Defective K-Ras oncoproteins overcome impaired effector activation to initiate leukemia in vivo. Blood *121(24)*, 4884–4893.

38. Shankar, S., Pitchiava, S., Malik, R., Kothari, V., Hosono, Y., Yocum, A.K., Gundlapalli, H., White, Y., Firestone, A., Cao, X., Dhanasekaran, S.M., Stuckey, J.A., Bollag, G.,

Shannon, K., Walter, N.G., Kumar-Sinha, C., and Chinnaiyan, A.M. (2016). KRAS engages AGO2 to enhance cellular transformation. Cell Rep. *14(6)*, 1448–1461.

39. Neale C, García AE. (2018) Methionine 170 is an environmentally sensitive membrane anchor in the disordered HVR of K-Ras4B. J Phys Chem B. *122(44)*, 10086–10096.

40. Sarkar-Banerjee, S., Sayyed-Ahmad, A., Prakash, P., Cho, K.-J., Waxham, M.N., Hancock, J.F., and Gorfe, A.A. (2017). Spatiotemporal analysis of K-Ras plasma membrane interactions reveals multiple high order homo-oligomeric complexes. J. Am. Chem. Soc. *139(38)*, 13466–13475.

41. Prakash, P., Sayyed-Ahmad, A., Cho, K.-J., Dolino, D.M., Chen, W., Li, H., Grant, B.J., Hancock, J.F., and Gorfe, A.A. (2017). Computational and biochemical characterization of two partially overlapping interfaces and multiple weak-affinity K-Ras dimers. Sci. Rep. *7*, 40109.

42. Šolman, M., Ligabue, A., Blaževitš, O., Jaiswal, A., Zhou, Y., Liang, H., Lectez, B., Kopra, K., Guzmán, C., Härmä, H., Hancock, J.F., Aittokallio, T., and Abankwa, D. (2015). Specific cancer-associated mutations in the switch III region of Ras increase tumorigenicity by nanocluster augmentation. eLife *4*, e08905.

43. Mazhab-Jafari, M.T., Marshall, C.B., Smith, M.J., Gasmi-Seabrook, G.M., Stathopulos, P.B., Inagaki, F., Kay, L.E., Neel, B.G., and Ikura, M. (2015). Oncogenic and RASopathy-associated K-RAS mutations relieve membrane-dependent occlusion of the effector-binding site. Proc. Natl. Acad. Sci. U.S.A. *112(21)*, 6625–6630.

44. Blaževitš, O., Mideksa, Y.G., Šolman, M., Ligabue, A., Ariotti, N., Nakhaeizadeh, H., Fansa, E.K., Papageorgiou, A.C., Wittinghofer, A., Ahmadian, M.R., and Abankwa, D. (2016). Galectin-1 dimers can scaffold Raf-effectors to increase H-ras nanoclustering. Sci. Rep. *6*, 24165.

45. Shalom-Feuerstein, R., Plowman, S.J., Rotblat, B., Ariotti, N., Tian, T., Hancock, J.F., and Kloog, Y. (2008). K-Ras nanoclustering is subverted by overexpression of the scaffold protein Galectin-3. Cancer Res. *68(16)*, 6608–6616.

46. Lin, Y.-H., Qiu, D.-C., Chang, W.-H., Yeh, Y.-Q., Jeng, U.-S., Liu, F.-T., and Huang, J.-R. (2017). The intrinsically disordered N-terminal domain of galectin-3 dynamically mediates multisite self-association of the protein through fuzzy interactions. J. Biol. Chem. *292(43)*, 17845–17856.

47. Yang, R.-Y., Hill, P.N., Hsu, D.K., and Liu, F.-T. (1998). Role of the carboxyl-terminal lectin domain in self-association of galectin-3. Biochemistry *37(12)*, 4086–4092.

48. Daub, M., Jöckel, J., Quack, T., Weber, C.K., Schmitz, F., Rapp, U.R., Wittinghofer, A., and Block, C. (1998). The RafC1 cysteine-rich domain contains multiple distinct regulatory epitopes which control Ras-dependent Raf activation. Mol. Cell Biol. *18(11)*, 6698–6710.

49. Williams, J.G., Drugan, J.K., Yi, G.-S., Clark, G.J., Der, C.J., and Campbell, S.L. (2000). Elucidation of binding determinants and functional consequences of Ras/Raf-cysteine-rich domain interactions. J. Biol. Chem. *275(29)*, 22172–22179.

50. Hu, C.-D., Kariya, K.-I., Tamada, M., Akasaka, K., Shirouzu, M., Yokoyama, S., and Kataoka, T. (1995). Cysteine-rich region of Raf-1 interacts with activator domain of post-translationally modified Ha-Ras. J. Biol. Chem. *270(51)*, 30274–30277.

51. Drugan, J.K., Khosravi-Far, R., White, M.A., Der, C.J., Sung, Y.-J., Hwang, Y.-W., and Campbell, S.L. (1996). Ras interaction with two distinct binding domains in Raf-1 may be required for Ras transformation. J. Biol. Chem. *271(1)*, 233–237.

52. Molzan, M., Kasper, S., Röglin, L., Skwarczynska, M., Sassa, T., Inoue, T., Breitenbuecher, F., Ohkanda, J., Kato, N., Schuler, M., and Ottmann, C. (2013).

Stabilization of physical RAF/14-3-3 interaction by cotylenin A as treatment strategy for RAS mutant cancers. ACS Chem. Biol. *8(9)*, 1869–1875.

53. Hatzivassiliou, G., Song, K., Yen, I., Brandhuber, B.J., Anderson, D.J., Alvarado, R., Ludlam, M.J.C., Stokoe, D., Gloor, S.L., Vigers, G., Morales, T., Aliagas, I., Liu, B., Sideris, S., Hoeflich, K.P., Jaiswal, B.S., Seshagiri, S., Koeppen, H., Belvin, M., Friedman, L.S., and Malek, S. (2010). RAF inhibitors prime wild-type RAF to activate the MAPK pathway and enhance growth. Nature *464(7287)*, 431–435.

54. Haling, J.R., Sudhamsu, J., Yen, I., Sideris, S., Sandoval, W., Phung, W., Bravo, B.J., Giannetti, A.M., Peck, A., Masselot, A., Morales, T., Smith, D., Brandhuber, B.J., Hymowitz, S.G., and Malek, S. (2014). Structure of the BRAF-MEK complex reveals a kinase activity independent role for BRAF in MAPK signaling. Cancer Cell *26(3)*, 402–413.

55. Park E, Rawson S, Li K, Kim BW, Ficarro SB, Pino GG, Sharif H, Marto JA, Jeon H, Eck MJ. (2019) Architecture of autoinhibited and active BRAF-MEK1-14-3-3 complexes. Nature. 575(7783), 545–550.

56. Cho, K.-J., Kasai, R.S., Park, J.-H., Chigurupati, S., Heidorn, S.J., van der Hoeven, D., Plowman, S.J., Kusumi, A., Marais, R., and Hancock, J.F. (2012). Raf inhibitors target Ras spatiotemporal dynamics. Curr. Biol. *22(11)*, 945–955.

57. Belanis, L., Plowman, S.J., Rotblat, B., Hancock, J.F., and Kloog, Y. (2008). Galectin-1 is a novel structural component and a major regulator of H-Ras nanoclusters. Mol. Biol. Cell *19(4)*, 1404–1414.

58. Elad-Sfadia, G., Haklai, R., Ballan, E., Gabius, H.-J., and Kloog, Y. (2002). Galectin-1 augments Ras activation and diverts Ras signals to Raf-1 at the expense of phosphoinositide 3-kinase. J. Biol. Chem. *277(40)*, 37169–37175.

59. Barceló, C., Paco, N., Morell, M., Alvarez-Moya, B., Bota-Rabassedas, N., Jaumot, M., Vilardell, F., Capella, G., and Agell, N. (2014). Phosphorylation at Ser-181 of oncogenic KRAS is required for tumor growth. Cancer Res. *74(4)*, 1190–1199.

60. Wang, Q., Pechersky, Y., Sagawa, S., Pan, A. C., Shaw, D. E. (2019). Structural mechanism for Bruton's tyrosine kinase activation at the cell membrane. Proceedings of the National Academy of Sciences, *116(19)*, 9390–9399.

61. Zhou, Y. and Hancock, J.F. (2015). Ras nanoclusters: versatile lipid-based signaling platforms. Biochim. Biophys. Acta 1853(4), 841–849.

62. Egea, P.F., Shan, S.O., Napetschnig, J., Savage, D.F., Walter, P., and Stroud, R.M. (2004). Substrate twinning activates the signal recognition particle and its receptor. Nature, *427(6971)*, 215.

63. Spencer-Smith, R., Koide, A., Zhou, Y., Eguchi, R.R., Sha, F., Gajwani, P., Santana, D., Gupta, A., Jacobs, M., Herrero-Garcia, E., Cobbert, J., Lavoie, H., Smith, M., Rajakulendran, T., Dowdell, E., Okur, M.N., Dementieva, I., Sicheri, F., Therrien, M., Hancock, J.F., Ikura, M., Koide, S., and O'Bryan, J.P. (2017). Inhibition of RAS function through targeting an allosteric regulatory site. Nat. Chem. Biol. *13(1)*, 62–68.

64. Shima, F., Ijiri, Y., Muraoka, S., Liao, J., Ye, M., Araki, M., Matsumoto, K., Yamamoto, N., Sugimoto, T., Yoshikawa, Y., Kumasaka, T., Yamamoto, M., Tamura, A., and Kataoka, T. (2010). Structural basis for conformational dynamics of GTP-bound Ras protein. J. Biol. Chem. *285(29)*, 22696–22705.

65. Bum-Erdene, K., Gagarinov, I.A., Collins, P.M., Winger, M., Pearson, A.G., Wilson, J.C., Leffler, H., Nilsson, U.J., Grice, I.D., and Blanchard, H. (2013). Investigation into the feasibility of thioditaloside as a novel scaffold for Galectin-3-specific inhibitors. ChemBioChem *14(11)*, 1331–1342.

66. Dharmaiah, S., Bindu, L., Tran, T.H., Gillette, W.K., Frank, P.H., Ghirlando, R., Nissley, D.V., Esposito, D., McCormick, F., Stephen, A.G., and Simanshu, D.K. (2016). Structural basis of recognition of farnesylated and methylated KRAS4b by PDEδ. Proc. Natl. Acad. Sci. U.S.A. *113(44)*, E6766–E6775.

67. Ismail, S.A., Chen, Y.-X., Rusinova, A., Chandra, A., Bierbaum, M., Gremer, L., Triola, G., Waldmann, H., Bastiaens, P.I.H., and Wittinghofer, A. (2011). Arl2-GTP and Arl3-GTP regulate a GDI-like transport system for farnesylated cargo. Nat. Chem. Biol. *7(12)*, 942–949.

68. Hoffman, G.R., Nassar, N., and Cerione, R.A. (2000). Structure of the Rho family GTP-binding protein Cdc42 in complex with the multifunctional regulator RhoGDI. Cell *100(3)*, 345–356.

#### Supplemental Figures

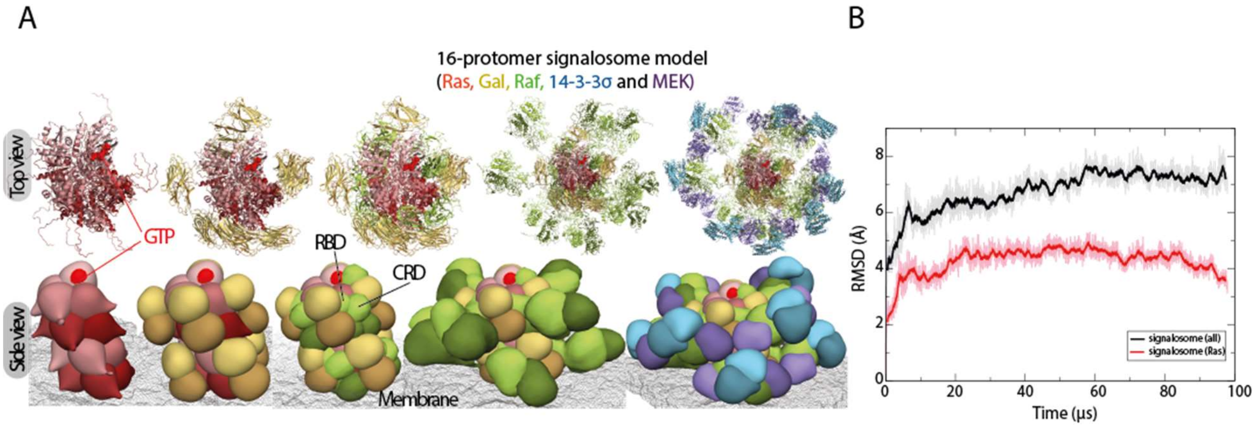


**Figure S1. Construction of the Ras-Raf signalosome model in 16-protomer form. A.** Top and side views of the stepwise addition (left to right) of Gal-3 (yellow), C-Raf (green), 14-3-3σ (blue), and MEK1 (violet) to the K-Ras helical assembly (red), producing the 16-protomer signalosome model. The membrane is shown as a mesh. Each C-Raf KD dimer binds to a 14-3-3σ dimer and two MEK1 kinases. **B.** The Cα atom RMSD (w.r.t. the starting structure) of a membrane-anchored 8-protomer Ras-Raf signalosome and its K-Ras octamer core in a 100-μs simulation. As shown, the signalosome—especially the K-Ras core—was stable in the course of the simulation.

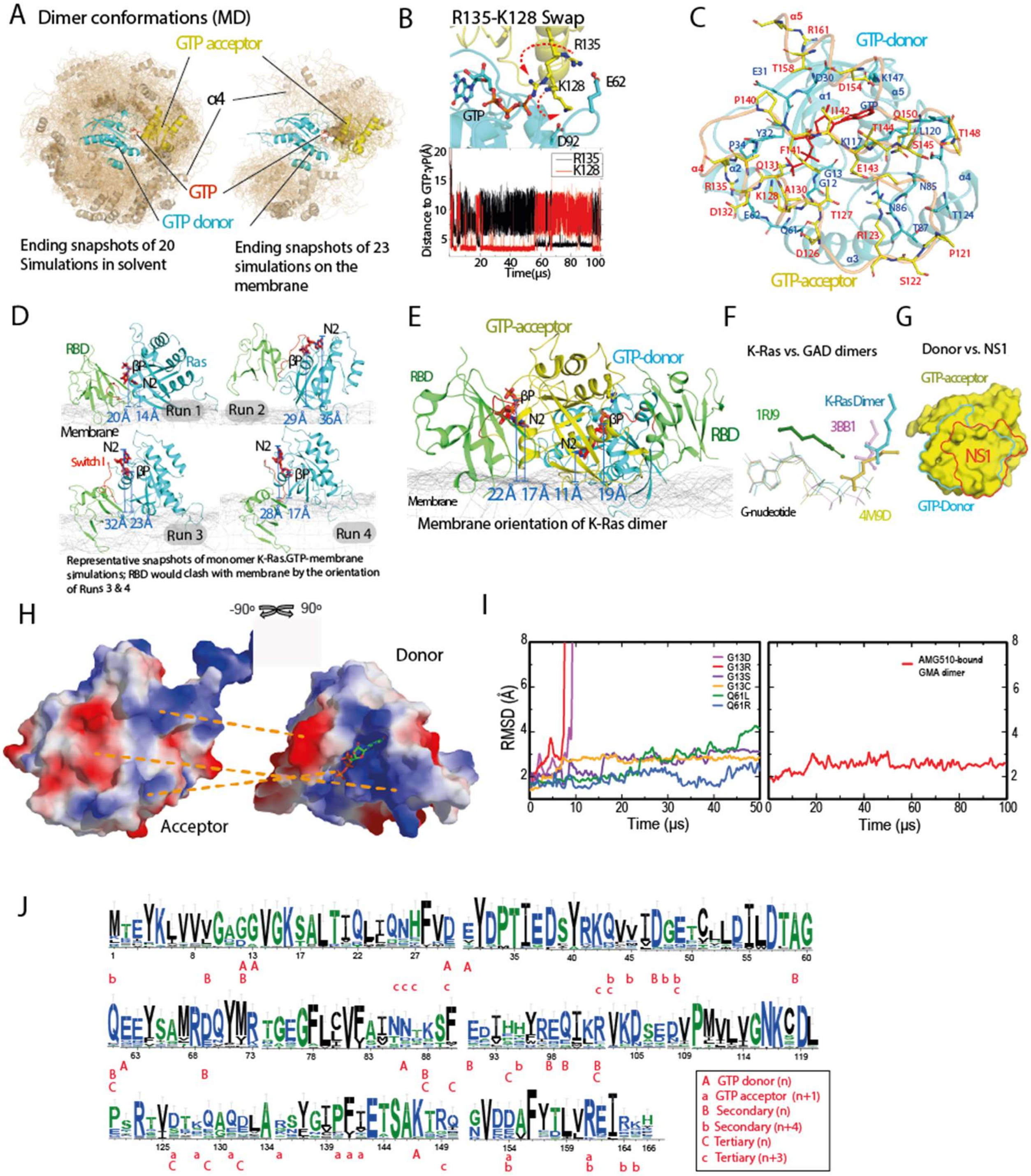


**Figure S2. The GTP-mediated K-Ras dimer model and its orientation on the membrane.**

**A.** Left panel: A collection of snapshots of the simulations starting from two separated K-Ras proteins in solvent. The snapshots are aligned to one K-Ras protein therein (cyan). The GMA dimer model is highlighted. The α4 helix is shown in cartoon representation. Snapshots 10 μs apart were taken from the 20 simulations (which have an aggregate simulation time of 680 μs). Right panel: A collection of snapshots of the simulations of two GTP-bound K-Ras proteins on the membrane; snapshots 10 μs apart were taken from the 23 simulations (which have an aggregate simulation time of 210 μs). Only in one solvent and one membrane simulation was the GMA dimer reached, and in both cases the dimer remained stable. **B.** Upper panel: Acceptor (yellow) interaction with the γ-phosphate of the GTP donor (cyan) in a simulation of a GMA K-Ras dimer. The Arg135 of the acceptor interacts with the γ-phosphate, while the Lys128 interacts with the Glu62 of the donor, and vice versa. Snapshots at 40 μs and 60 μs from the simulation of K-Ras dimerization on the membrane are shown; the motions of Arg135 and Lys128 are shown by dashed red arrows. Lower panel: Distances from Arg135 (black) and Lys128 (red) to the γ-phosphate. **C.** Interface residues of the GMA dimer. **D.** Snapshots at 10 μs from four independent runs of free GTP-bound K-Ras monomer (cyan) on the membrane. To illustrate the relationship between the membrane orientation and RBD binding of K-Ras, the C-Raf RBD (green) is positioned on each snapshot based on the Ras-RBD structure (PDB 4G0N). Shown are the distance of the β phosphorus (βP) and the amine nitrogen of the guanine ring (N2) from the plane of the membrane surface; we use these distances to describe the membrane orientation of a Ras protein (Figure 2G). The Run 1 snapshot is compatible with RBD binding, which corresponds to the Ras membrane orientation of the lower red contours in Figure 2G in the main text. The snapshot from Run 2 corresponds to the right center of the top red contours; this orientation of K-Ras on the membrane positions the RBD away from the membrane. In the Run 3 and Run 4 snapshots, RBD binding leads to a severe steric clash with the membrane; this corresponds to the left center of the top red contour. **E.** Snapshot at 100 μs from the simulation of the GMA dimer formation on the membrane, with two C-Raf RBDs (green) added based on the Ras-RBD pose; the RBDs do not clash with the membrane. The membrane orientation of the donor and acceptor Ras proteins correspond to the lower and upper black contours, respectively, in Figure 2G in the main text. **F.** Examples of G nucleotide–mediated dimerization, in which an arginine interacts with a nucleotide phosphate at the dimerization interface. The Toc34 homodimer[30] (PDB 3BB1), the Ffh-FtsY heterodimer[62] (PDB 1RJ9), and the adenylosuccinate synthetase homodimer (PDB 4M9D) are compared to the GMA K-Ras dimer model (cyan); the host G proteins (not shown) were aligned in this comparison. **G.** The binding site of the synthetic monobody NS1[63] (red) and the donor site (cyan) of an acceptor in the GMA dimer are shown on a Ras protein (yellow). The two binding sites largely overlap. **H.** The electrostatic complementarity at the GMA dimer interface, with contacting areas at the interface connected by dashed lines. **I.** Cα RMSD of the GMA dimer of the wild type and various oncogenic mutants (left panel), and the GMA dimer of the G12C mutant bound with covalent inhibitor AMG510 (right panel) in simulations. **J.** Sequence conservation of K-Ras proteins in various species (see Materials and Methods). A graphical representation of sequence alignment of representative K-Ras proteins in evolution. The residues involved in the K-Ras/K-Ras interactions in the K-Ras helical assembly are labeled. The 32 K-Ras sequences by GenBank IDs are: OLS17184.1, OLS30914.1, OLS23071.1, XP_020603162.1, XP_023347765.1, XP_023347766.1, XP_003378992.1, XP_003377451.1, GBC14959.1, XP_027484897.1, QBM87817.1, XP_013758736.1, EHB13737.1, OQV11758.1, XP_027289840.1, NP_001356715.1, XP_027290137.1, XP_027290057.1, XP_027289839.1, XP_027289566.1, XP_029436565.1, ELK24704.1, XP_027290375.1, XP_023664743.1, XP_021777811.1, XP_021247616.1, XP_025064324.1, XP_006127724.1, NP_001243091.1, RDD38924.1, KPM02442.1, OTF69949.1.

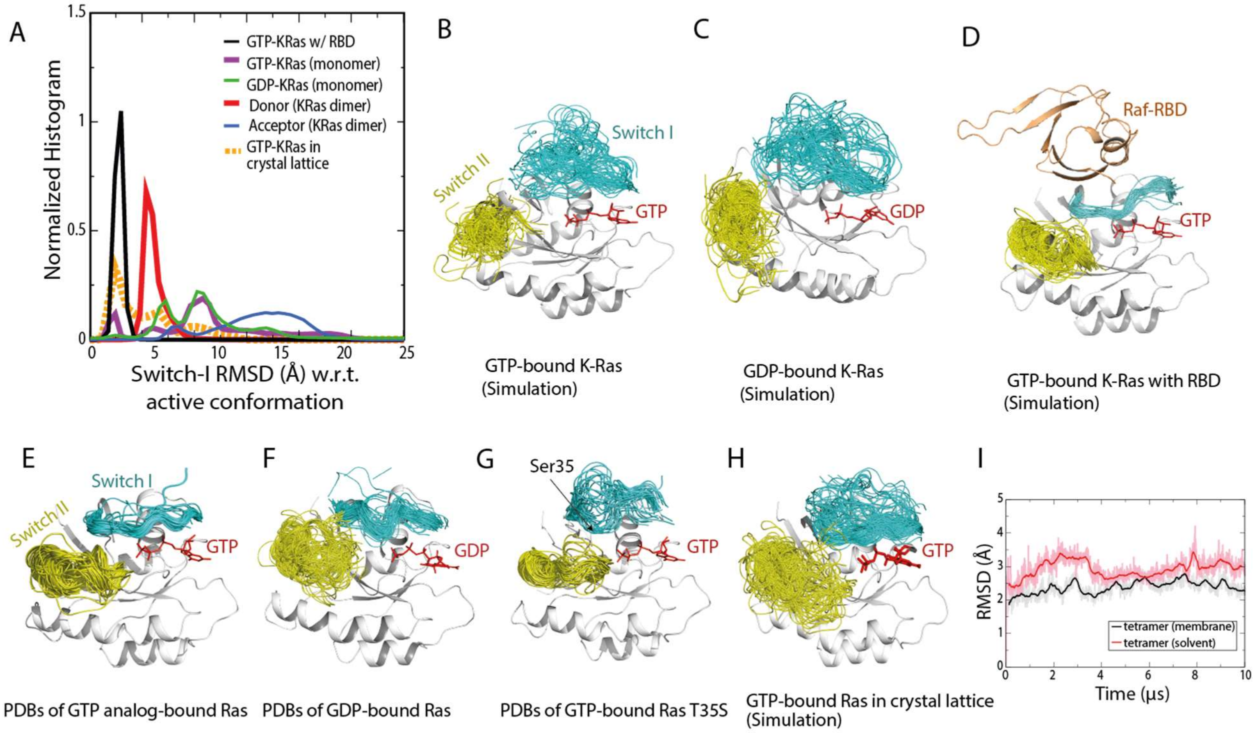


**Figure S3. Stability of the active switch I and switch II conformations. A.** Distributions of the switch I conformation of K-Ras in various simulations. Normalized histograms of the switch I backbone root-mean-square deviation (RMSD) with respect to the active switch I conformation in PDB 4DSN are shown. **B.** Conformations of the switch I (residues 30–38, cyan) and switch II (residue 60–76, yellow) regions in simulations of the GTP-bound K-Ras monomer. **C.** Conformations of the switch I and II regions in simulations of GDP-bound K-Ras monomer. **D.** Conformations of the switch I and II regions in simulations of GTP-bound K-Ras bound with C-Raf RBD. **E.** Crystal structures of Ras bound with GTP or GTP analogs, with the switch I and switch II regions highlighted. The PDB entries included are 1AGP, 1CLU, 1CTQ, 1GNP, 1GNR, 1HE8, 1JAH, 1JAI, 1K8R, 1LF0, 1LFD, 1NVU, 1NVV, 1NVW, 1NVX, 1P2S, 1P2T, 1P2U, 1P2V, 1PLJ, 1PLK, 1QRA, 1RVD, 1ZW6, 2C5L, 2RGA, 2RGB, 2RGC, 2RGD, 2RGE, 2RGG, 2UZI, 2VH5, 3DDC, 3GFT, 3I3S, 3K8Y, 3L8Y, 3L8Z, 3LBH, 3LBI, 3LBN, 3OIU,

3OIV, 3OIW, 3RRY, 3RRZ, 3RS0, 3RS2, 3RS3, 3RS4, 3RS5, 3RS7, 3RSO, 3TGP, 3V4F, 4DLR, 4DLS, 4DLT, 4DLU, 4DLV, 4DLW, 4DLX, 4DLY, 4DLZ, 4DSN, 4DSO, 4DST, 4EFL, 4EFM, 4EFN, 4G0N, 4G3X, 4K81, 4L9W, 4NMM, 4NYI, 4NYJ, 4NYM, 4RSG, 4XVQ, 4XVR, 5B2Z, 5B30, 5P21, 6Q21, 121P, 421P, 521P, 621P, 721P, 821P, and 221P. **F.** Crystal structures of Ras loaded with GDP. The PDB entries included are 1AA9, 1CRP, 1CRQ, 1CRR, 3LO5, 1IOZ, 1LF5, 1PLL, 1Q21, 1WQ1, 1XD2, 1XJ0, 1ZVQ, 2CE2, 2CLD, 2Q21, 2QUZ, 2X1V, 3CON, 3KUD, 4DSU, 4EPR, 4EPT, 4EPV, 4EPW, 4EPX, 3EPY, 4L8G, 4L9S, 4LDJ, 4LPK, 4LRW, 4LUC, 4LV6, 4LYF, 4LYH, 4LYJ, 4M1O, 4M1S, 4M1T, 4M1W, 4M1Y, 4M21, 4M22, 4OBE, 4PZY, 4PZZ, 4Q01, 4Q02, 4Q03, 4Q21, 4QL3, 4TQ9, 4TQA, 4WA7, and 5F2E. **G.** Crystal structures of the T35S Ras mutant loaded with GTP or GTP analog, with the switch I and switch II regions highlighted. The PDB entries included are 1IAQ, 2LCF, 2LWI, 3KKM, and 3KKN. The hydroxyl of Thr35 in the switch I region in wild-type Ras coordinates the $Mg^{2+}$ ion bound to the GTP, and the T35S mutation is known to disrupt the switch I active conformation. The switch I inactive conformations sampled by the simulations (Figure S3B and S3C) are broadly consistent with the inactive conformations in T35S structures.[64]
**H.** Conformations of the switch I and II regions in simulations of 24 copies of GTP-bound K-Ras in a crystal lattice (of PDB 3GFT). **I.** The Cα atom RMSDs of a GMA K-Ras tetramer (w.r.t. the starting structure) in a 10-μs simulation with membrane and in a 10-μs simulation in water solvent. As shown, the membrane helps stabilize the tetramer structure.

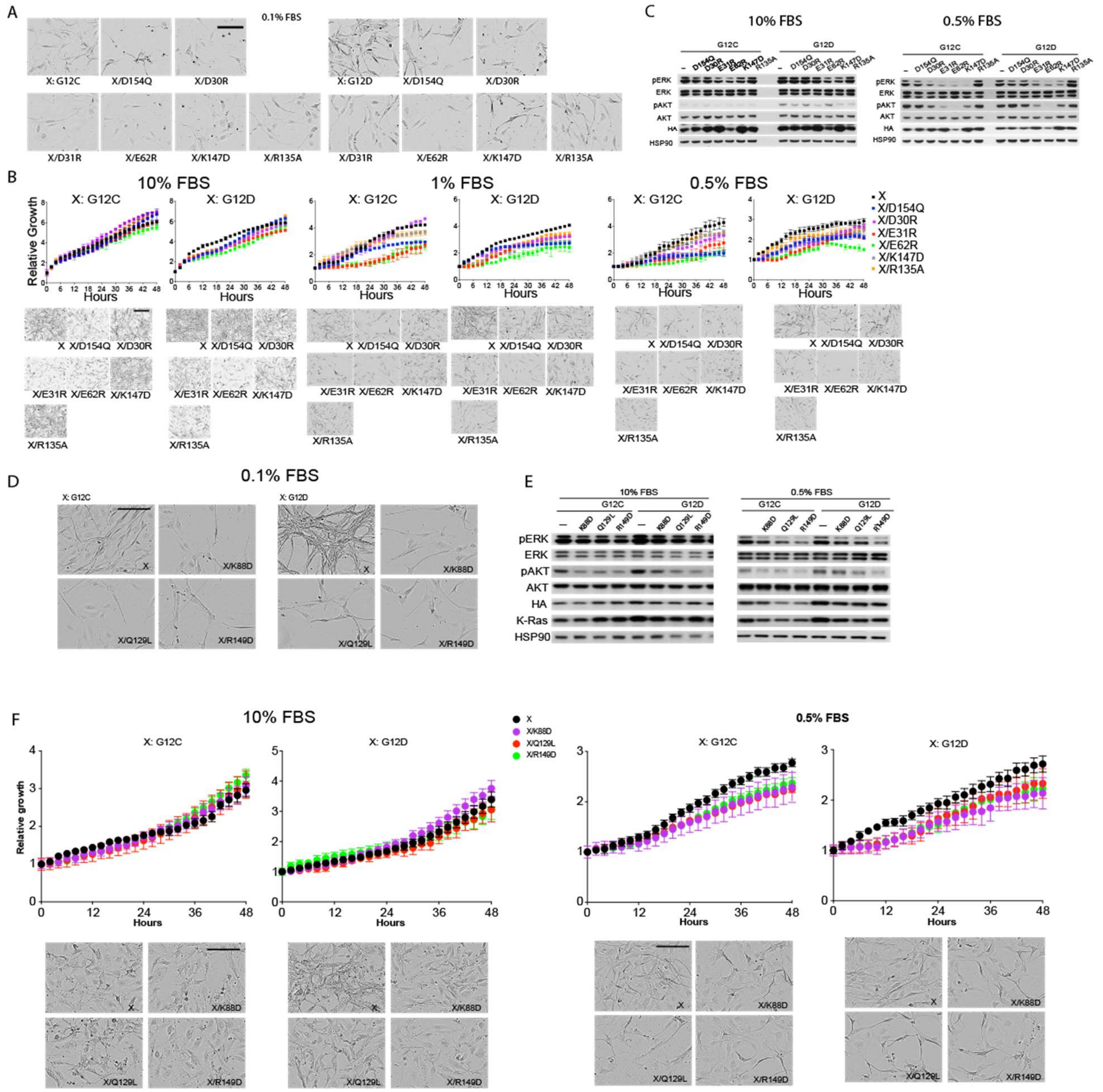


**Figure S4. Cellular validation of the K-Ras signalosome interfaces.** Panels **A**–**C** here refer to the experiments testing mutations at the GMA dimer interface, and panels **D**–**F** refer to the experiments testing secondary/tertiary K-Ras interfaces. **A**. Representative cell images at the endpoint of the experiment testing mutations at the GMA dimer interface (Figure 3F); scale bar:

50 µm. **B.** Growth rates of *K-Ras*$^{lox}$/*K-RAS*$^{MUT}$ cells expressing various K-Ras mutants in 10%, 1%, and 0.5% FBS medium, represented by the confluence value assessed by IncuCyte. Representative pictures at the endpoint are shown in the bottom panels (scale bar: 50 µm). Results are representative from three replicates. **C.** Phosphorylation of ERK and AKT in *K-Ras*$^{lox}$/*K-RAS*$^{MUT}$ cells. Cells were lysed after 48 hours incubation in 0.5% or 10% FBS, as indicated, and analyzed by Western blot. **D.** Representative cell images at the endpoint of the experiment testing mutations at the secondary and tertiary K-Ras/K-Ras interfaces (Figure 6B); scale bar: 50 µm. **E**. Phosphorylation of ERK and AKT in *K-Ras*$^{lox}$/*K-RAS*$^{MUT}$ cells. Cells were lysed after 48 hours incubation in 0.5% or 10% FBS as indicated and analyzed by Western blot. **F**. Growth rates of *K-Ras*$^{lox}$/*K-RAS*$^{MUT}$ cells expressing various K-Ras mutants to test the secondary and tertiary K-Ras/K-Ras interfaces in the background of either G12C or G12D, shown as confluence values measured by IncuCyte. Results are averages of three duplicates with error bars. Representative images at the end point are also shown. Cells were kept in 0.5% or 10% FBS.

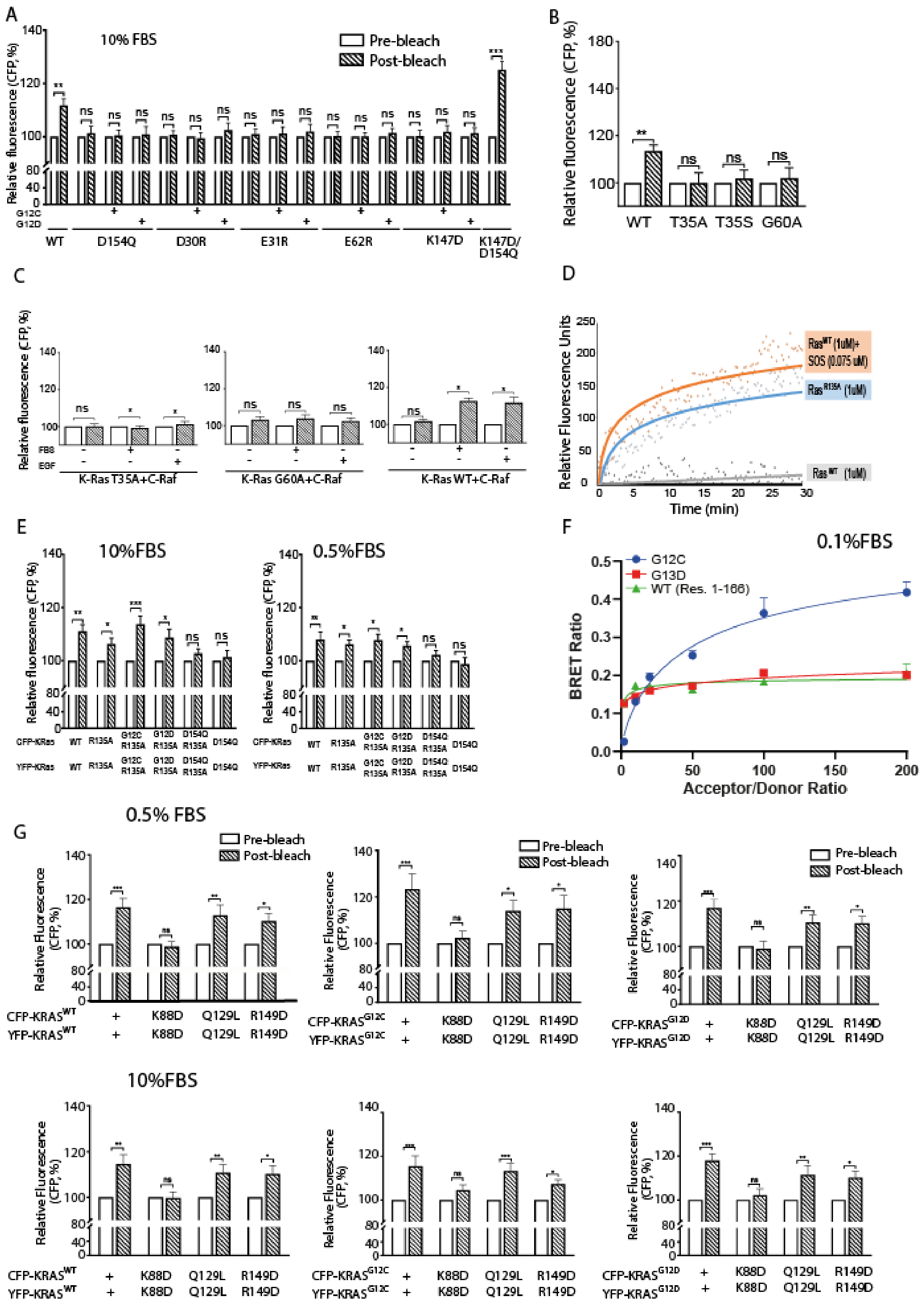


**Figure S5. FRET and BRET validation of K-Ras signalosome interfaces. A**. CFP emission of wild-type K-Ras and various K-Ras mutants to test the K-Ras/K-Ras GMA dimer. HEK293T cells were co-transfected with CFP-K-Ras and YFP-K-Ras, serum starved, and stimulated with 10% FBS. Each K-Ras construct is labeled under its respective plot. **B.** CFP emission of wild-type K-Ras and T35 and G60 mutants to test K-Ras association. **C**. CFP emission of wild-type K-Ras and T35 and G60 mutants with C-Raf to monitor Ras-Raf binding at 10% FBS or 10 ng $mL^{-1}$ EGF. **D.** Nucleotide exchange assay of wild-type K-Ras with (orange) and without (gray) SOS1, and R135A without SOS1 (blue). **E.** CFP emission of wild-type K-Ras and R135A mutants. **F**. BRET signal as an indicator of K-Ras assembly for G12C and G13D mutants, and for the K-Ras construct lacking the membrane-anchoring HVR tail (residues 1–166). Co-transfection of increasing ratios of donor and acceptor plasmids enables discrimination between specific and non-specific (random collision) protein-protein interactions. **G.** CFP emission of wild-type K-Ras and K88D, Q129L, and R149D mutants. In (A), (B), (C), (E), (F), and (H), error bars represent mean ± SD, * denotes $p < 0.05$, ** denotes $p < 0.01$, and *** denotes $p < 0.001$ by the unpaired Student's t-test; ns stands for not significant.

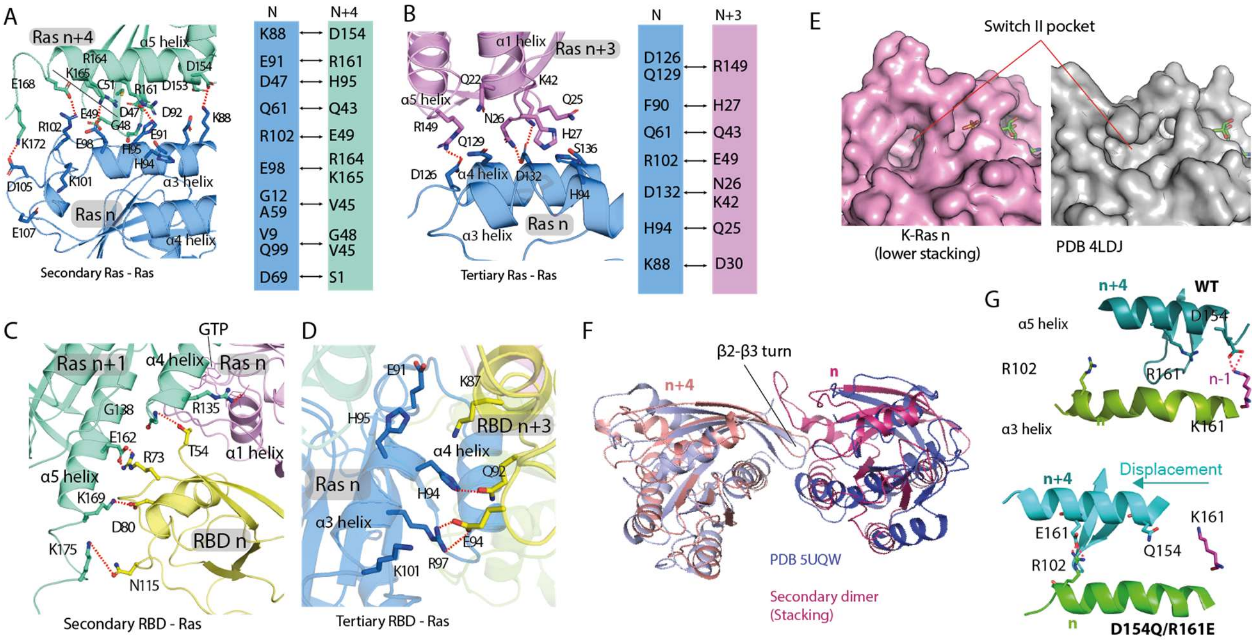


**Figure S6. Structural details of secondary and tertiary Ras-Ras and Ras-RBD interactions.** **A.** Key contacts of the secondary (stacking) Ras-Ras interaction of K-Ras *n* (blue) and K-Ras *n+4* (green). A series of salt bridges form intermittently in MD simulations between the Lys88, Glu91, Glu98, Arg102, and Asp105 residues on the α3 helix of K-Ras *n* and the Asp154, Arg161, Lys165, Glu168, and Lys172 residues, respectively, on the α5 helix of K-Ras *n+4*. **B.** Key contacts at the interface of K-Ras *n+3* (purple) and K-Ras *n* (blue). **C.** Key contacts of the secondary Ras-RBD interaction of C-Raf RBD *n* (yellow) with K-Ras *n+1* (green), which is the GTP acceptor of K-Ras *n* (purple), the primary binding partner of the RBD. The RBD also interacts with the HVR of K-Ras *n+1*. Residues 1–53 of C-Raf are structurally unresolved but are likely to make additional interactions at this secondary Ras-RBD interface. **D.** Key contacts of the tertiary Ras-RBD interaction of Raf RBD *n+3* (yellow) and K-Ras *n* (blue). **E.** Unoccupied Switch-II pocket in a crystal structure of K-Ras (PDB 4LDJ) compared with the

Switch-II pocket of K-Ras *n* in the stacking interaction. **F.** Superposition of stacking (n and n+4) K-Ras proteins from the signalosome model with a crystal dimer of K-Ras (PDB 5UQW); in both cases the dimer interaction is mediated by a β2- β3 turn inserting into a Switch-II pocket. **G.** Comparison of D154Q/R161E with the WT in K-Ras oligomer simulations. As shown, the double mutation disrupts the D154-K147 salt bridge at the GMA dimer (n/n-1), and as compensation, allows an E161-R102 salt bridge in stacking (n/n+4) by shifting the relative position of the α3 and α5 helices.

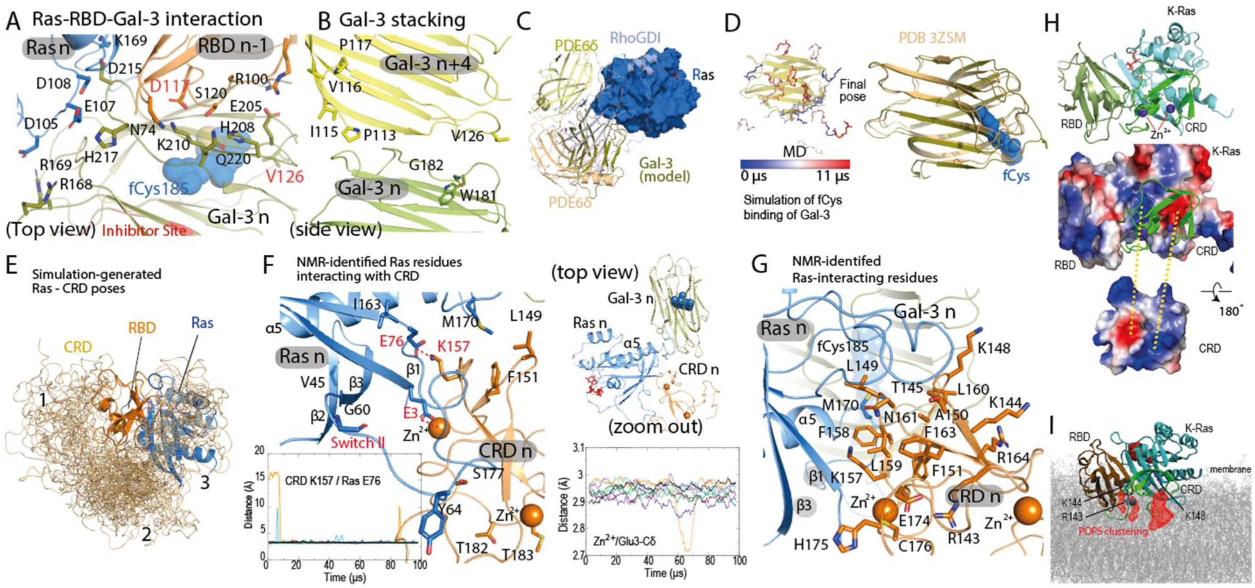


**Figure S7. Structural details of interactions of K-Ras, Gal-3 and the C-Raf CRD. A.** Gal-3 *n* (olive) binds to the HVR fCys185 of Ras *n* (blue) and to C-Raf RBD *n−1* (orange). The thiodigalactoside inhibitor binding site of Gal-3[65] (PDB 4JC1) is shown in red. **B.** Gal-3 stacking; Gal-3 *n* (green) and Gal-3 *n+4* (yellow) are stacked in parallel with K-Ras stacking. **C.** Position of Gal-3 (olive) with respect to K-Ras (blue), in comparison with that of PDE6δ (orange) with respect to farnesylated K-Ras[66] (PDB 5TAR), PDE6δ (yellow) with respect to farnesylated Rheb[67] (PDB 3T5G), and RhoGDI (cyan) with respect to geranylgeranylated CDC42[68] (PDB 1DOA). The Gal-3 position somewhat resembles the PDE6δ complex in the overall pose. The insert shows the simulation-generated Gal-3 binding pose of fCys with the Gal-3 crystal structure (PDB 3ZSM) overlaid for comparison. **D.** Left: snapshots of the simulation in which fCys enters the Gal-3 (yellow) pocket, with color coding on fCys indicating simulation time; right: the final Gal-3/fCys model superimposed on the Gal-3 structure used to initiate the simulation (PDB 3ZSM). **E.** Final snapshots from 55 simulations of the three-domain system of the CRD (light orange) tethered to a K-Ras-bound RBD (orange). The snapshots were aligned with respect to the K-Ras protein. Of the 55 simulations, 24 were in solvent (80 µs in aggregate), and the other 31 were with the RBD and the membrane (117 μs in aggregate). The simulation-generated CRD poses are grouped into three clusters: 1) clashing with the GMA K-Ras dimer; 2) clashing with the membrane; and 3) near the C-terminal of the α5 helix and the β1, β2, and β3 strands of K-Ras. **F.** Details of the CRD-Ras interface of a CRD pose selected from the third cluster. NMR-identified Ras residues involved in Ras-CRD interaction are shown. The Glu3 residue on the β1 strand of the K-Ras protein interacts stably with a CRD-bound $Zn^{2+}$ ion (lower right panel). The Lys157(CRD)/Glu76(K-Ras) distance in simulations is shown to indicate the stability of the Ras-CRD pose (lower left panel). The upper right panel shows the relative positions of K-Ras (blue), CRD (orange), Gal-3 (olive), and GTP (red) in a top view. **G.** CRD residues identified by an NMR study[49] as part of the K-Ras interface. K-Ras M170, fCys185, and CRD $Zn^{2+}$ ions are also shown. **H.** The surfaces of RBD-bound K-Ras and CRD colored by their electrostatic potential (calculated using PyMOL). A cartoon image is used to show the orientation of the Ras-CRD complex. The CRD-bound $Zn^{2+}$ ions are shown in purple. **I.** A C-Raf CRD at the base tier with RBD and K-Ras. Arg143, Lys144, Lys148, and the two coordinating $Zn^{2+}$ ions of the CRD are shown, together with the occupancy map of POPS lipids in contact with the three amino acids shown in red mesh.

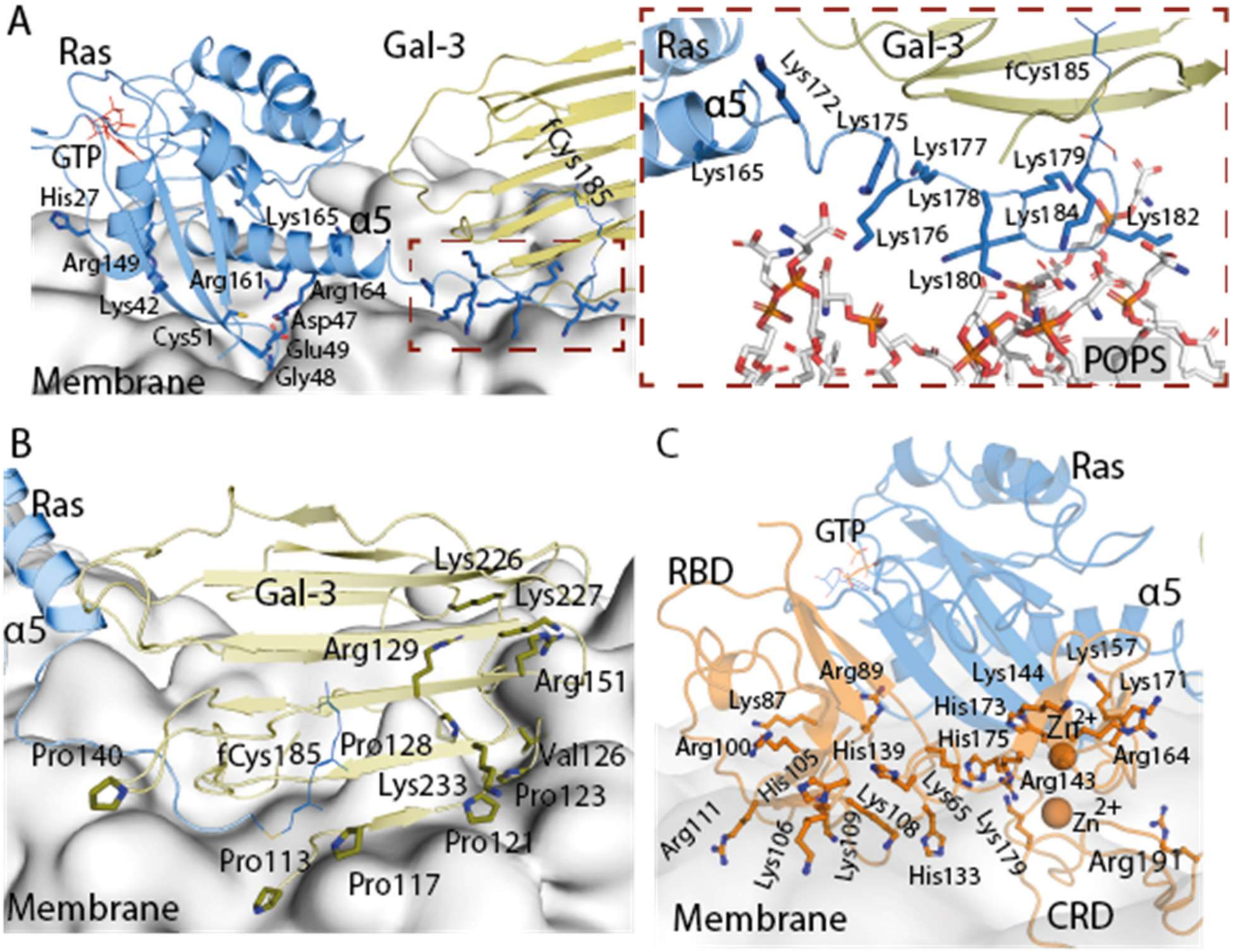


**Figure S8. Membrane interface of K-Ras, Gal-3, and C-Raf. A.** Positively charged residues of a base-tier K-Ras protein proximal to the membrane in the signalosome model. The inset shows the direct interaction between the HVR lysines of a K-Ras protein at the base tier and the phosphotidylserine lipids. **B.** Positively charged residues and prolines of Gal-3 proximal to the membrane at the base tier. A Val126 residue that has been suggested to be involved in Gal-3 membrane interaction[45] is also shown. **C.** Positively charged residues of the C-Raf RBD and CRD and the two CRD-bound $Zn^{2+}$ ions proximal to the membrane.

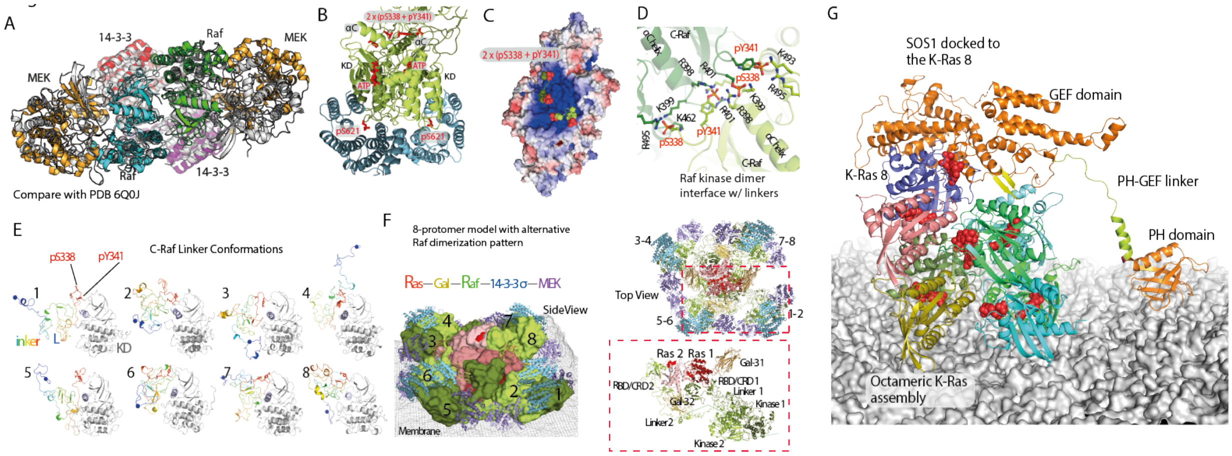


**Figure S9. Additional interactions between Raf, 14-3-3σ, and MEK1. A.** Superposition of the structural model (gray) of the Raf/MEK/14-3-3 complex with the separately resolved cryo-EM structure (PDB 6Q0J). **B.** Interaction between the Raf kinase domain dimer (green) and 14-3-3σ dimer (cyan). The linker residues pSer338 and pTyr341 are located between the two αC helices of the kinase dimer, and pSer621 is bound to a 14-3-3σ protein. **C.** Electrostatic potential surface of the C-Raf kinase domain dimer; the region of the two αC helices, which is where pSer339 and pTyr341 (shown) are located, is highly positively charged. **D.** Interaction of phosphorylated residues pSer338 and pTyr341 with the positively charged residues of the two αC helices of the C-Raf KD dimer. **E.** Samples of conformations of the C-Raf linker (rainbow colors) in the Ras-Raf signalosome model. The C-Raf KD (gray) is attached to the linker for reference. **F.** Top and side views of an alternative eight-protomer signalosome model, wherein C-Raf *n* and C-Raf *n+1* form a dimer, as opposed to the model shown in Figure 1D and 1E, wherein C-Raf *n* and C-Raf *n+4* form a dimer. The color scheme and protein representation are similar to those in Figure 1D. The inset shows that in this alternative model a C-Raf linker (linker 2) has to circumvent a Gal-3 protein (Gal-3 2) to engage the other C-Raf linker in a C-Raf dimer. **G.** An illustration of how the GEF domain of SOS1 can reach the K-Ras *8*. The modeling was based on PDB entry 1NVU for the K-Ras/SOS1 pose, and PDB entry 1XD4 for the structures of the PH domain and the PH-GEF linker; the linker orientation with respect to the PH and the GEF domains was manually adjusted and energetically minimized in the modeling. The helical hairpin of SOS1 is shown in yellow.

#### Supplemental Movie Legends

**Movie S1, related to Figure 1. The architecture of the Ras-Raf signalosome model.** Illustration of the 16-protomer signalosome, in which a 16-member helical assembly of K-Ras on the membrane was assembled, followed by sequential addition of Gal-3, the RBD, CRD, the linker and KD of C-Raf, the 14-3-3σ dimer, and finally MEK1 kinase. Colors are consistent with those in Figure 1D.

**Movie S2, related to Figure S2A (left panel). Simulation of K-Ras dimer formation in solvent.** An unbiased simulation starting with two spatially separated GTP-bound K-Ras proteins in solvent, in which the GTP-mediated asymmetric dimer formed spontaneously at approximately 16 μs. The simulation time is marked. The GTP donor is colored in cyan and the GTP acceptor in pink. At the end of the movie, this dimer is superimposed with the GTP-mediated asymmetric dimer that was generated by simulating K-Ras dimerization on a membrane.

**Movie S3, related to Figure 2A. Simulation of K-Ras dimer formation on a membrane.** An unbiased simulation starting with two spatially separated GTP-bound K-Ras proteins on a membrane, in which the GTP-mediated asymmetric dimer formed spontaneously at approximately 6.5 μs. The simulation time is marked. The GTP donor is colored in cyan and the GTP acceptor in pink. The membrane is shown in gray in the background; GTP molecules are also shown.

#### Supplemental Structure File Legends

**Dataset S1, related to Figure 1D.** Atomic coordinates of the structural model of the eight-protomer Ras-Raf signalosome with K-Ras (chains A, B, P, Q, W, Y, E, G), C-Raf (chains C, D, R, S, X, Z, F, H), Gal-3 (chains N, O, V, U, I, J, K, T), 14-3-3σ (chains n, o, v, u, i, j, k, t), and MEK1 (chains c, d, r, s, x, z, f, h).

**Dataset S2, related to Figure 1A.** Atomic coordinates of the GMA K-Ras dimer on a membrane.
